# Database and deep learning toolbox for noise-optimized, generalized spike inference from calcium imaging

**DOI:** 10.1101/2020.08.31.272450

**Authors:** Peter Rupprecht, Stefano Carta, Adrian Hoffmann, Mayumi Echizen, Antonin Blot, Alex C. Kwan, Yang Dan, Sonja B. Hofer, Kazuo Kitamura, Fritjof Helmchen, Rainer W. Friedrich

## Abstract

Calcium imaging is a key method to record patterns of neuronal activity across populations of identified neurons. Inference of temporal patterns of action potentials (‘spikes’) from calcium signals is, however, challenging and often limited by the scarcity of ground truth data containing simultaneous measurements of action potentials and calcium signals. To overcome this problem, we compiled a large and diverse ground truth database from publicly available and newly performed recordings. This database covers various types of calcium indicators, cell types, and signal-to-noise ratios and comprises a total of >35 hours from 298 neurons. We then developed a novel algorithm for spike inference (CASCADE) that is based on supervised deep networks, takes advantage of the ground truth database, infers absolute spike rates, and outperforms existing model-based algorithms. To optimize performance for unseen imaging data, CASCADE retrains itself by resampling ground truth data to match the respective sampling rate and noise level. As a consequence, no parameters need to be adjusted by the user. To facilitate routine application of CASCADE we developed systematic performance assessments for unseen data, we openly release all resources, and we provide a user-friendly cloud-based implementation.

## INTRODUCTION

Imaging of somatic calcium signals using organic or genetically encoded fluorescent indicators has emerged as a key method to measure the activity of many identified neurons simultaneously in the living brain^1, 2^. However, calcium signals are only an indirect, often non-linear and low pass-filtered proxy of the more fundamental variable of interest, somatic action potentials (spikes)^3–5^. The relationship between calcium signals and spike rates is ideally assessed directly by simultaneous electrophysiological recordings — preferably in the minimally disruptive juxtacellular configuration — and optical imaging of a calcium indicator signal in the same neuron. These dual recordings can serve as ground truth to calibrate and optimize algorithms for the inference of spike rates from other calcium imaging data (Fig. 1a). Based on such ground truth datasets, various model-based methods^6–16, 12^ as well as supervised machine learning algorithms^15, 17–20^ for spike inference have been developed.

**Figure 1.**
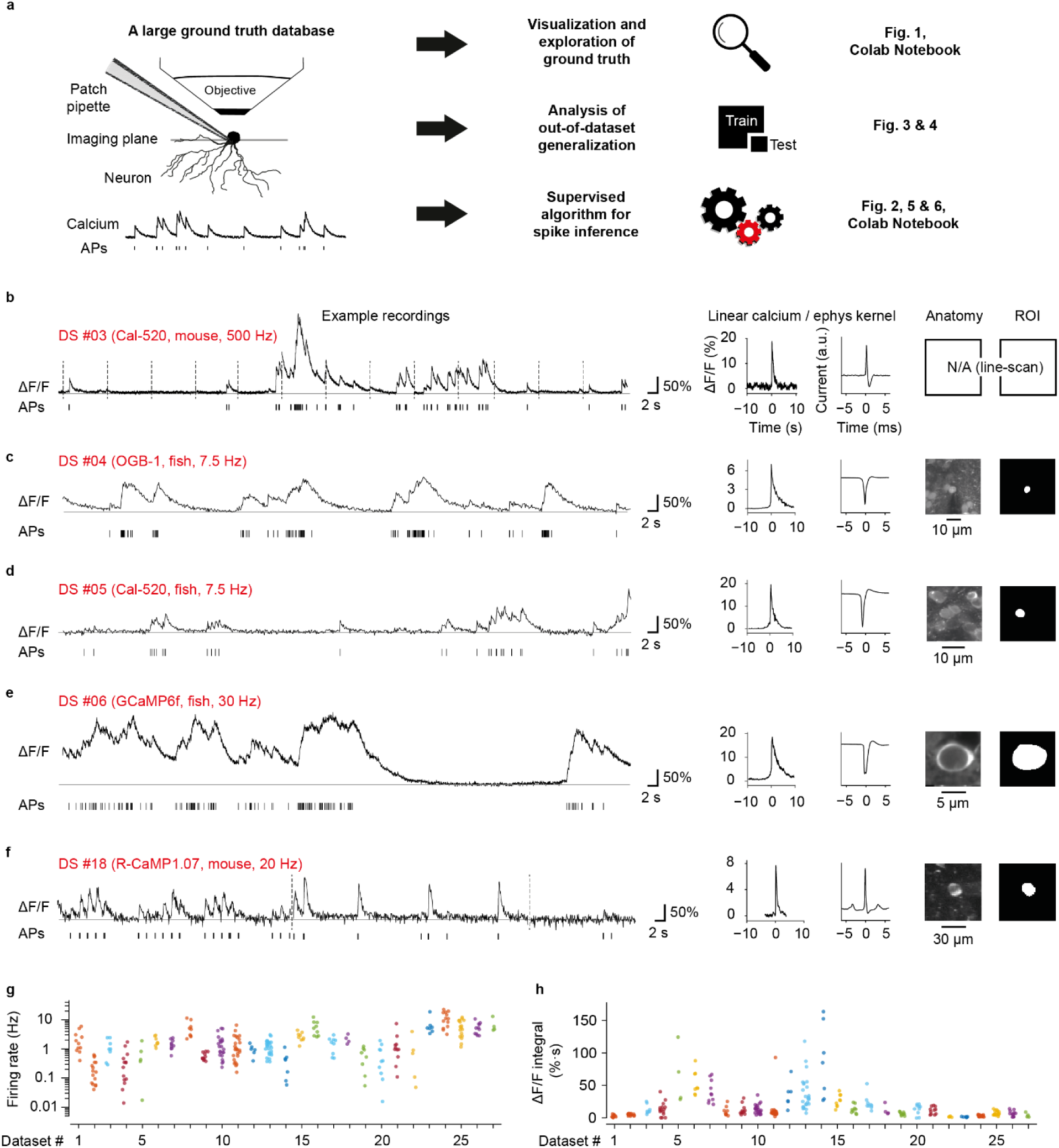
Ground truth datasets. **a,** A large and diverse ground truth database obtained by simultaneous calcium imaging and juxtacellular recording (left) can be used 1) for the exploration of the ground truth by a user, 2) for the analysis of the out-of-dataset generalization of spike inference and 3) for the training of a supervised algorithm for spike inference. The right column refers to relevant figures. *Colab Notebook* refers to relevant cloud-based tools accompanying this paper. **b-f,** Examples of ground truth recordings with different indicators, different brain regions and species. Left: calcium signal traces (ΔF/F) are shown together with the detected action potentials (APs). Dashed lines indicate breaks during recordings. Middle: linear kernels of ΔF/F (time scale in seconds) and electrophysiological data (time scale in milliseconds) triggered by single spikes. Right: fluorescence image of the respective neuron, together with the ROI for fluorescence extraction. **g,** Average spike rate for each neuron of the ground truth database (log scale). 27 datasets (DS) were included in total. Interneuron datasets comprise DS#22-27. **h,** Integral ΔF/F of the spike kernel (first 2 s) for each neuron. Lowest values are observed in PV+ interneurons (DS#23 and #24). See Fig. S1 for the underlying kernels.

Ideally, an algorithm should be applicable to infer spike rates in unseen calcium imaging datasets for which no ground truth is available. The relationship between spikes and the evoked calcium signals depends on multiple factors including the neuron type, the type of calcium indicator and its concentration, the optical resolution, the sampling rate and the noise level. Many of these parameters can vary substantially between experiments and even from neuron to neuron within the same experiment. As a consequence, experimental conditions of novel datasets are often not well matched to those of available ground truth data. It is therefore not clear how an algorithm based on a specific ground truth dataset generalizes to other datasets, which causes problems for the inference of spike rates from calcium imaging data under most experimental conditions^12,13,21,22^.

Here, we address the issue of generalization systematically. To assemble a large ground truth database, we performed juxtacellular recordings and two-photon calcium imaging using different calcium indicators and in different brain regions of zebrafish and mice. This database was then augmented with a carefully curated selection of publicly available ground truth datasets. Using this large database, we developed a supervised method for **<u>ca</u>**librated **<u>s</u>**pike inference of **<u>ca</u>**lcium data using **<u>de</u>**ep networks (CASCADE). CASCADE includes methods to resample the original ground truth datasets in order to match their sampling rate and noise level to a specific calcium imaging dataset of interest. This procedure allowed us to train machine learning algorithms upon demand on a broad spectrum of resampled ground truth datasets, matching a wide range of experimental conditions. Finally, we tested the performance of CASCADE systematically when applied to unseen data. CASCADE was robust with respect to any hyper-parameter choices and performed better than existing algorithms in benchmark tests performed across all ground truth datasets and across noise levels. The CASCADE algorithm can be used directly via a cloud-based web application and is also available together with the ground truth datasets as a simple and user-friendly Python-based toolbox.

## RESULTS

### A large dataset of curated ground truth recordings

To extend the spectrum of existing ground truth datasets, we performed simultaneous electrophysiological recordings and calcium imaging in adult zebrafish and mice (Fig. 1b-h; Table 1). In zebrafish, a total of 47 neurons in different telencephalic regions were recorded in the juxtacellular configuration in an explant preparation of the whole adult brain^23^ using the synthetic calcium indicators Oregon Green BAPTA-1 (OGB- 1) and Cal-520 as well as the genetically encoded calcium indicator GCaMP6f. Additional ground truth recordings were performed in head-fixed, anesthetized mice in hippocampal area CA3 using the genetically encoded indicator R-CaMP1.07 ^24^. Furthermore, we extracted ground truth recordings from raw data of previous publications using Cal-520 and R-CaMP1.07, respectively, in mouse primary somatosensory cortex (S1; total of 21 neurons)^25, 26^, and from interneurons in mouse primary visual cortex (V1) using OGB- 1 *in vivo* and GCaMP6f in slices (total of 69 neurons)^27, 28^, complemented with a small new *in vivo* interneuron dataset using GCaMP6f (4 neurons). In addition, we surveyed openly accessible datasets and extracted ground truth from raw movies (when available) or preprocessed calcium imaging data^15, 18, 29–33^. Rigorous quality control (Methods) reduced the original number from a total of 193 available neurons to 157 neurons. Together with our own recordings, we have assembled 27 datasets comprising a total of 298 neurons, 8 calcium indicators and 9 brain regions in 2 species, totaling ∼38 hours of recording and 495,077 spikes.

**Table 1.** Overview of all ground truth datasets. Standardized noise of calcium ΔF/F was determined as described in Methods. Frame rate is given as the mean across experiments if the frame rates varied (typically only slightly) across experiments within a single dataset. Noise levels and spike rates are given as mean ± s.d. across neurons.

| Dataset identifier | Calcium indicator | Induction method | Animal model | Brain region | Frame rate [Hz] | Standardized noise [%-Hz<sup>-1/2</sup>] | Spike rate [Hz] | # of neurons | Recording duration [min] | Source paper |
| --- | --- | --- | --- | --- | --- | --- | --- | --- | --- | --- |
| #1 | OGB-1 | acute injection | Mouse | V1 | 11.3 | 0.7±0.2 | 5.5±1.5 | 11 | 83 | Theis et al., 2016 |
| #2 | OGB-1 | injection +<br>tg(tdTomato-CaMKIIα) | Mouse | V1 | 15.6 | 0.5±0.1 | 0.2±0.2 | 16 | 116 | Kwan and Dan, 2012 |
| #3 | Cal-520 | acute injection | Mouse | S1 | 500.0 | 0.3±0.1 | 1.2±0.8 | 8 | 23 | Tada et al., 2014 |
| #4 | OGB-1 | acute injection | Zebrafish | pDp | 7.7 | 1.0±0.2 | 0.4±0.5 | 15 | 81 | this paper |
| #5 | Cal-520 | acute injection | Zebrafish | pDp | 7.8 | 2.0±1.3 | 1.3±2.1 | 5 | 31 | this paper |
| #6 | GCaMP6f | tg(NeuroD) | Zebrafish | aDp | 30.0 | 1.3±0.8 | 1.9±0.7 | 8 | 46 | this paper |
| #7 | GCaMP6f | tg(NeuroD) | Zebrafish | dD | 30.0 | 0.6±0.1 | 1.5±0.6 | 10 | 69 | this paper |
| #8 | GCaMP6f | tg(NeuroD) | Zebrafish | OB | 30.0 | 0.8±0.2 | 5.3±3.3 | 9 | 45 | this paper |
| #9 | GCaMP6f | AAV | Mouse | V1 | 60.1 | 0.4±0.1 | 0.6±0.2 | 11 | 129 | Chen et al., 2013 |
| #10 | GCaMP6f | tg(Emx1) | Mouse | V1 | 160.1 | 0.5±0.2 | 1.6±1.4 | 23 | 72 | Huang et al., 2019 |
| #11 | GCaMP6f | tg(Cux2) | Mouse | V1 | 158.3 | 0.5±0.2 | 1.5±1.5 | 25 | 78 | Huang et al., 2019 |
| #12 | GCaMP6s | tg(tetOs) | Mouse | V1 | 151.6 | 0.8±0.1 | 1.0±0.4 | 6 | 13 | Huang et al., 2019 |
| #13 | GCaMP6s | tg(Emx1) | Mouse | V1 | 157.5 | 0.5±0.2 | 1.3±0.7 | 26 | 62 | Huang et al., 2019 |
| #14 | GCaMP6s | AAV | Mouse | V1 | 60.1 | 0.5±0.2 | 0.4±0.4 | 7 | 70 | Chen et al., 2013 |
| #15 | GCaMP6s | AAV | Mouse | V1 | 59.1 | 0.7±0.2 | 6.2±3.5 | 9 | 77 | Theis et al., 2016 |
| #16 | GCaMP6s | AAV | Mouse | V1 | 59.1 | 0.9±0.2 | 5.8±3.3 | 9 | 25 | Theis et al., 2016 |
| #17 | GCaMP5k | AAV | Mouse | V1 | 50.0 | 0.5±0.2 | 1.6±0.9 | 9 | 29 | Akerboom et al., 2012 |
| #18 | R-CaMP1.07 | tg(Grik4-cre) + AAV | Mouse | CA3 | 20.0 | 1.6±0.3 | 2.2±0.8 | 4 | 33 | Schoenfeld et al., 2021 |
| #19 | R-CaMP1.07 | AAV | Mouse | S1 | 15.0 | 0.6±0.2 | 0.9±1.0 | 9 | 50 | Bethge et al., 2017 |
| #20 | jRCaMP1a | AAV | Mouse | V1 | 15.0 | 1.3±0.5 | 0.6±0.6 | 10 | 88 | Dana et al., 2016 |
| #21 | jRGECO1a | AAV | Mouse | V1 | 29.8 | 1.0±0.3 | 1.6±2.0 | 11 | 118 | Dana et al., 2016 |
| #22 | OGB-1 | injection + tg(GFP-GIN) | Mouse | V1 (SST) | 15.6 | 0.6±0.1 | 1.1±1.6 | 5 | 49 | Kwan and Dan, 2012 |
| #23 | OGB-1 | injection +<br>tg(tdTomato-PV) | Mouse | V1 (PV) | 15.6 | 0.6±0.1 | 6.9±5.4 | 7 | 17 | Kwan and Dan, 2012 |
| #24 | GCaMP6f | tg(PV-cre) + AAV | Mouse (in vitro) | V1 (PV) | 26.6 | 0.5±0.2 | 11.6±5.4 | 13 | 215 | Khan et al., 2018 |
| #25 | GCaMP6f | tg(SOM-cre) + AAV | Mouse (in vitro) | V1 (SST) | 26.6 | 0.6±0.2 | 5.9±3.5 | 17 | 375 | Khan et al., 2018 |
| #26 | GCaMP6f | tg(VIP-cre) + AAV | Mouse (in vitro) | V1 (VIP) | 26.6 | 2.1±1.3 | 5.5±2.5 | 11 | 252 | Khan et al., 2018 |
| #27 | GCaMP6f | Td(tdTomato-PV)<br>+AAV | Mouse | V1 (PV) | 30.0 | 2.8±1.1 | 7.0±4.1 | 4 | 30 | this paper |

Recording durations, imaging frame rates and spike rates varied greatly across ground truth datasets (Table 1). Typical spike rates spanned more than an order of magnitude, ranging from 0.4 to 11.6 Hz, and frame rates varied between 7.7 Hz and >160 Hz (Table 1; Fig. 1g). We used regularized deconvolution to compute the linear ΔF/F kernel evoked by the average spike and found that the area under the kernel curve varied substantially across datasets, even when data were obtained with the same indicator, and was substantially smaller for interneuron datasets, especially for parvalbumin-positive (PV) interneurons (Fig. 1h). Interestingly, kernels varied substantially even across neurons within the same dataset (Fig. 1h, Fig. S1). This diversity highlights the challenge faced by any algorithm that is supposed to generalize to unseen data.

### Inference of spike rates with a deep convolutional network

Several favorable properties make supervised deep learning approaches well suited for spike inference from calcium imaging data. First, deep learning generally tends to outperform other classification or regression methods if the amount of training data is sufficiently high (typically >1000 data points for each category in classification tasks) ^34–36^. Second, the cost function can easily be modified to optimize the metric of interest, *e.g.*, correlation with ground truth or mean squared error, without changing network architecture. Third, the temporal extent of receptive fields of deep networks can be adapted to account for history-dependent effects such as the dependence of action potential-evoked calcium transients on previous activity (see Fig. S2 for an example). Finally, deep networks are intrinsically non-linear, allowing to fit non-linear behaviors of calcium indicators.

We designed a simple convolutional network that uses a segment of the calcium signal trace (expressed as percentage fluorescence change ΔF/F) around a time point *t* to infer the spike rate at *t*. Compared to two-dimensional image classification and object labeling^34, 37, 38^, requirements on computational hardware are low because datasets are small and the inference task is only one-dimensional (time). For example, ImageNet^39^, a dataset used for visual object identification and detection in the deep learning field, is typically used at a resolution of 256 x 256 = 65,536 data points per sample, whereas the input used for spike inference in this study was smaller by approximately three orders of magnitude, typically consisting of a segment of the ΔF/F trace with 64 data points.

We used a network architecture with a standard convolutional design, consisting of rectifying linear units (ReLUs) that were distributed across three convolutional layers, two pooling layers and a single dense layer. The final dense layer projected to a single output unit that reported the estimated spike rate for the current time *t* (Fig. 2a; see Methods for more details).

**Figure 2.**
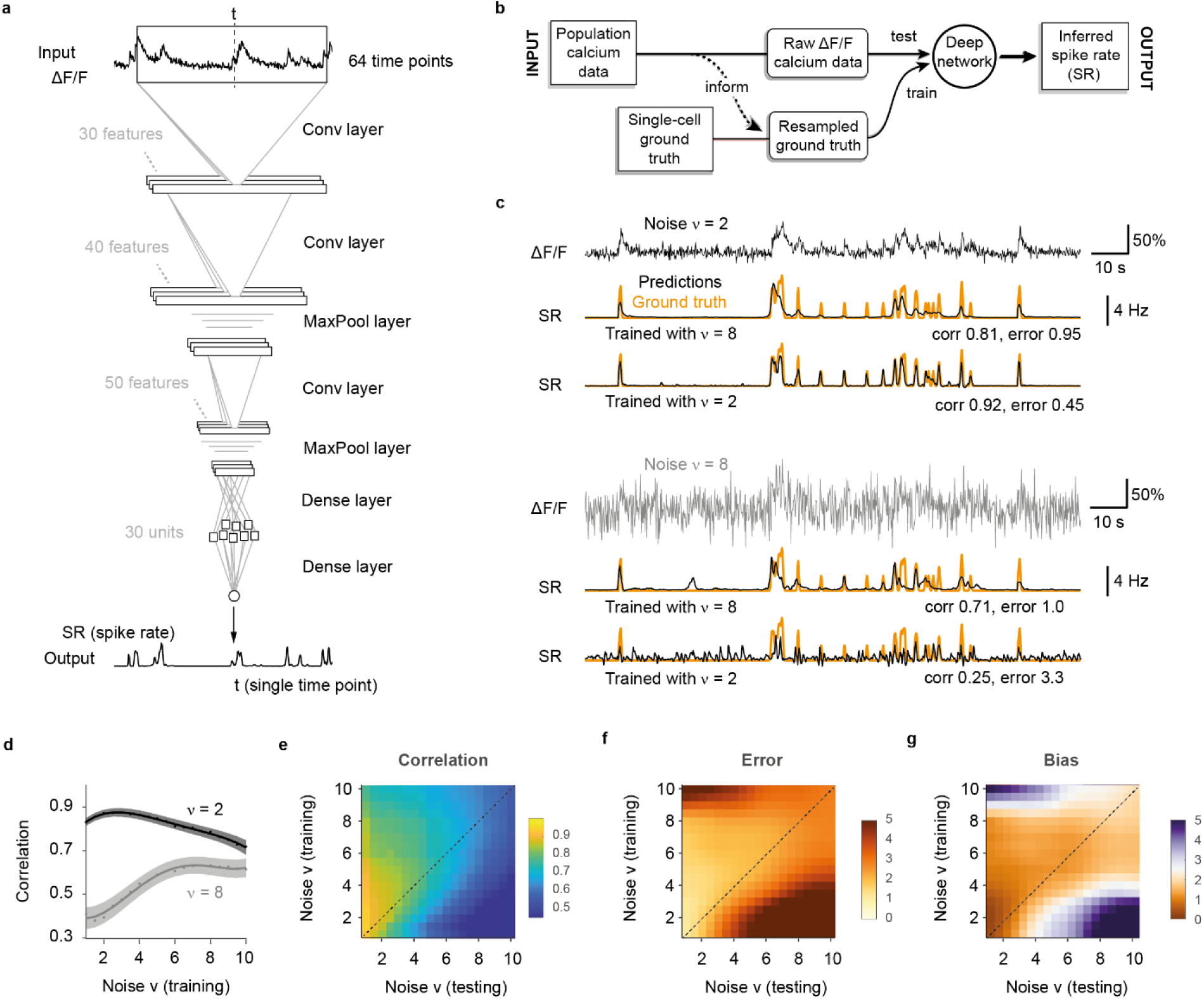
Training a deep network with noise-matched ground truth improves spike inference. **a,** The default deep network consists of an input time window of 64 time points centered around the time point of interest. Through three convolutional layers, two pooling layers and one small dense layer, the spiking probability is extracted from the input time window and returned as a single number for each time point. **b,** Properties of the population data (frame rate, noise level; dashed line) are extracted and used for noise- matched resampling of existing ground truth datasets. The resampled ground truth is used to train the algorithm, resulting in calibrated spike inference of the population imaging data. **c**, Top: a low-noise ΔF/F trace is translated into spike rates (SR; inferred spike rates in black, ground truth in orange) more precisely when low-noise ground truth has been used for training. Bottom: a high-noise ΔF/F trace is translated into spike rates (SR; inferred spike rates in black, ground truth in orange) more precisely when high-noise ground truth has been used for training. ν in units of standardized noise, % · *Hz*^−1/2^. **d,** The spike inference performance for two test conditions (low noise, ν = 2, dark gray; high noise, ν = 8, light gray) is optimal when training noise approximates testing noise levels. **e,** Correlation between predictions and ground truth is maximized if noise levels of training datasets match noise levels of testing sets. **f,** Relative error of predictions with respect to ground truth. **g,** Relative bias of predictions with respect to ground truth. Column-wise normalized versions of (e-g) are shown in Fig. S6.

### Resampling of ground truth data for noise-matching

The key idea underlying our approach is that the ground truth (training data) is as important as the algorithm itself and should match as well as possible the noise level and sampling rate of the unseen population calcium data of interest (test data). We therefore devised a workflow where noise level and sampling rate are extracted from the test data and then used to generate noise- and rate-matched training data from the ground truth database (Fig. 2b) by temporal resampling and addition of noise. To facilitate gradient descent, the ground truth spike rate is smoothed with a Gaussian kernel (σ = 0.2 s, unless otherwise indicated; Methods).

To extract ΔF/F noise levels, we computed a standardized noise metric ν that is robust against outliers and approximates the standard deviation of ΔF/F baseline fluctuations. This metric was normalized by the square root of the frame rate to allow for comparison of noise measurements across datasets. Consequently, ν has units of %·Hz^-1/2^, which for simplicity we usually omit (Methods; Fig. S3). To generate training data with pre-defined ΔF/F noise levels, we explored several approaches based on sub-sampling of ROIs or additive artificial noise (Supplementary Note ‘Noise-matching of resampled ground truth data‘; Fig. S4). We identified the addition of artificial Poisson-distributed noise as the most suitable approach to transform the ground truth data into appropriate training data for the deep network.

To quantify deep network performance, we developed a set of complementary metrics for the accuracy of spike inference (equations and illustrations in Fig. S5). Following previous studies, we calculated the Pearson **correlation** between ground truth spike rates and inferred spike rates^15, 18^. However, this correlation measure of performance leaves the absolute magnitude of the inferred instantaneous spike rate unconstrained. We therefore also computed the positive and negative deviations of the absolute spike rate from the ground truth spike rate. The sum of the absolute deviations was defined as the **error** while the sum of the signed deviations was defined as the **bias** of the inference (Methods; Fig. S5). Error and bias were both normalized by the number of true spikes to obtain relative metrics that can be compared between datasets. Among these three metrics (correlation, error, bias), correlation is arguably the most important one because it estimates the similarity of inferred and true spike rates. Error and bias are relevant for the inference of absolute spike rates because they identify spike rates that are either incorrectly scaled or systematically too large or small.

The performance of the deep network degraded considerably when the noise level of the test dataset deviated substantially from the noise level of the ground truth. As expected by intuition, a network that had only seen almost noise-free data during training failed to suppress fluctuations in noisier recordings. Conversely, we also observed that a network trained on very noisy calcium signals was unable to fully benefit from low-noise calcium recordings, inferring only an imprecise approximation of the ground truth (Fig. 2c). A systematic iteration across combinations of noise levels for training and test datasets showed that for each test noise level the best model had been trained with a similar or slightly higher noise level (Fig. 2d-g; even more apparent when normalizing the metrics for each test noise level in Fig. S6). Very low noise levels (ν < 2) result in a special case (Fig. 2d,e): since some neurons of a given ground truth dataset do not reach the desired noise level even without addition of noise (cf. Table 1), the effective size of the training dataset decreases, resulting in slightly lower performance. In general, however, the results show that it is beneficial to train with noise levels that are adapted to the calcium data for which the algorithm will be applied after training.

### Parameter-robustness of spike inference

Traditional models to infer spiking activity typically contain a small number of parameters^10–12, 14^ that describe biophysical quantities and are adjusted by the user. Deep networks, in contrast, contain thousands or millions of parameters adjusted during training that have no obvious biophysical meanings^13, 15^. The user can modify only a small number of hyper-parameters that define general properties of the network such as the loss function, the number of features per layer, or the receptive field size, *i.e.,* the size of the input window shown in Fig. 2a. We therefore tested how spike inference performance depends on these hyper- parameters.

We found that the performance of the network was robust against variations of all hyper-parameters (Supplementary Note ‘Dependence of performance on hyper-parameters, overfitting and network architecture’; Fig. S7a-e), allowing us to leave all parameters unchanged for all conditions. Moreover, overfitting was moderate despite prolonged training, indicating that the abundance of noise and sparseness of events act as a natural regularizer (same Supplementary Note; Fig. S7f-h). Finally, we tested different deep learning architectures including non-convolutional or recurrent long short-term memory (LSTM) networks. While very large networks tended to slightly overfit the data, most networks performed almost equally well (same Supplementary Note; Fig. S8). Hence, the expressive power of moderately deep networks and the robustness of back-propagation with gradient descent enables multiple different networks to find good models for spike inference irrespective of the network architecture, hyper-parameter settings and the chosen learning procedure. This high robustness of the deep learning approach practically eliminates the need for manual adjustments of hyper-parameters.

### Generalization across neurons within the same dataset

Ideally, the ground truth data used to train a network should match the experimental conditions in the test dataset (calcium indicator type, labeling method, concentration levels, brain region, cell type, etc.). To explore spike inference under such conditions we measured how well spike rates of a given neuron within a ground truth dataset can be predicted by networks that were trained using the other neurons in the dataset. First, all ground truth calcium ΔF/F data were resampled to a common sampling rate and adjusted to the same noise levels by adding Poisson noise. If the initial noise level of a given ground truth neuron was higher than the target noise level, the neuron was excluded from the respective noise level analysis. We then evaluated the performance of CASCADE as a function of the noise levels of the (re-sampled) datasets. As expected, correlations increased and errors decreased for lower noise levels, while average biases seemed not to be systematically affected (Fig. S9a-d). Performance metrics also varied considerably across different neurons within a single dataset when resampled at the same noise level ν. To better understand this variability, we performed additional analyses.

First, we found spike-evoked calcium transients to be variable across neurons from the same dataset (Fig. 1h, Fig. S1). Large errors and biases, as well as low correlations, were observed when spike-evoked calcium transients of a neuron deviated strongly from those of other neurons (red arrow in Fig. S9d; cf. Fig. S1r for the respective linear kernels of DS#18).

Second, spike inference may be complicated by movement artifacts or neuropil contamination. Movement artifacts typically had slow onset- and offset-kinetics (Fig. S10a), or a faster, quasi-periodic temporal structure related to breathing (Fig. S10d-e). Neuropil contamination is often very difficult to distinguish from somatic calcium signals and particularly severe when neurons are tightly packed and densely labeled^1, 40–43^ (Fig. S10b). For a subset of datasets, we tested the effect of simple center-surround subtraction of the neuropil signal^29^. Because subtraction is not perfect, decontaminated datasets still contained residual neuropil signals (Fig. S10b) or negative transients (Fig. S10c). Nonetheless, spike inference was significantly improved by neuropil decontamination (Fig. S11). More detailed inspection of the results showed that CASCADE was able to learn to ignore negative transients and movement artifacts, but only as long as they were distinguishable from true calcium transients. For example, artifacts were only sometimes correctly not associated with spikes (Fig S10a-c), limiting the precision of spike inference.

Third, we found that the activity of sparsely spiking neurons is less well predicted since the calcium signal of single action potentials is more likely to be overwhelmed by shot noise, particularly in the high-noise regime (arrows in Fig. S9a,c). We therefore evaluated conditions required for single-spike precision and observed that either shot noise or other noise sources were too prominent in all ground truth datasets to allow for reliable single-spike detection. The trained network thus systematically underestimated single spikes (Fig. S12). This observation was made using GCaMP indicators, which show a strongly nonlinear relationship between calcium concentration and fluorescence and therefore are less sensitive to isolated single spikes occurring during low baseline activity, but also using synthetic dyes (Fig. S12). These observations indicate that the network needs to learn a tradeoff between false-positive detections of noise events and false-negative detections of single spikes. Further details related to single-spike precision and the possibility to discretize inferred spike rates are discussed in the Supplementary Note ‘Discrete spikes and single-spike precision.

In summary, we showed that CASCADE is able to generalize to unseen neurons from the same ground truth training set. Not surprisingly, the precision of this generalization decreases for higher noise levels, in particular when spike rates are low. The precision is fundamentally limited by the variability of calcium kernels across neurons and probably also by the non-linearity of GCaMP-like indicators, and precision is further reduced when additional noise (motion artifacts, neuropil contamination) is prominent.

### Generalization across datasets

We next explored how spike inference by a network trained on one ground truth dataset generalizes to other datasets. Using all available datasets, we quantified the median performance metrics across all possible combinations of datasets for training and testing (Fig. 3a,c,e) and analyzed the performance of each trained model across test datasets (Fig. 3b,d,f). In most training/test combinations, correlations were high whereas errors and biases remained low. However, models trained with some datasets showed low performance across datasets (*e.g.,* DS#01), and some datasets, often with motion or neuropil contamination artifacts that were larger than typical calcium transients, could not be predicted well by any model trained on another dataset (*e.g.,* DS#01-02, DS#21-23).

**Figure 3.**
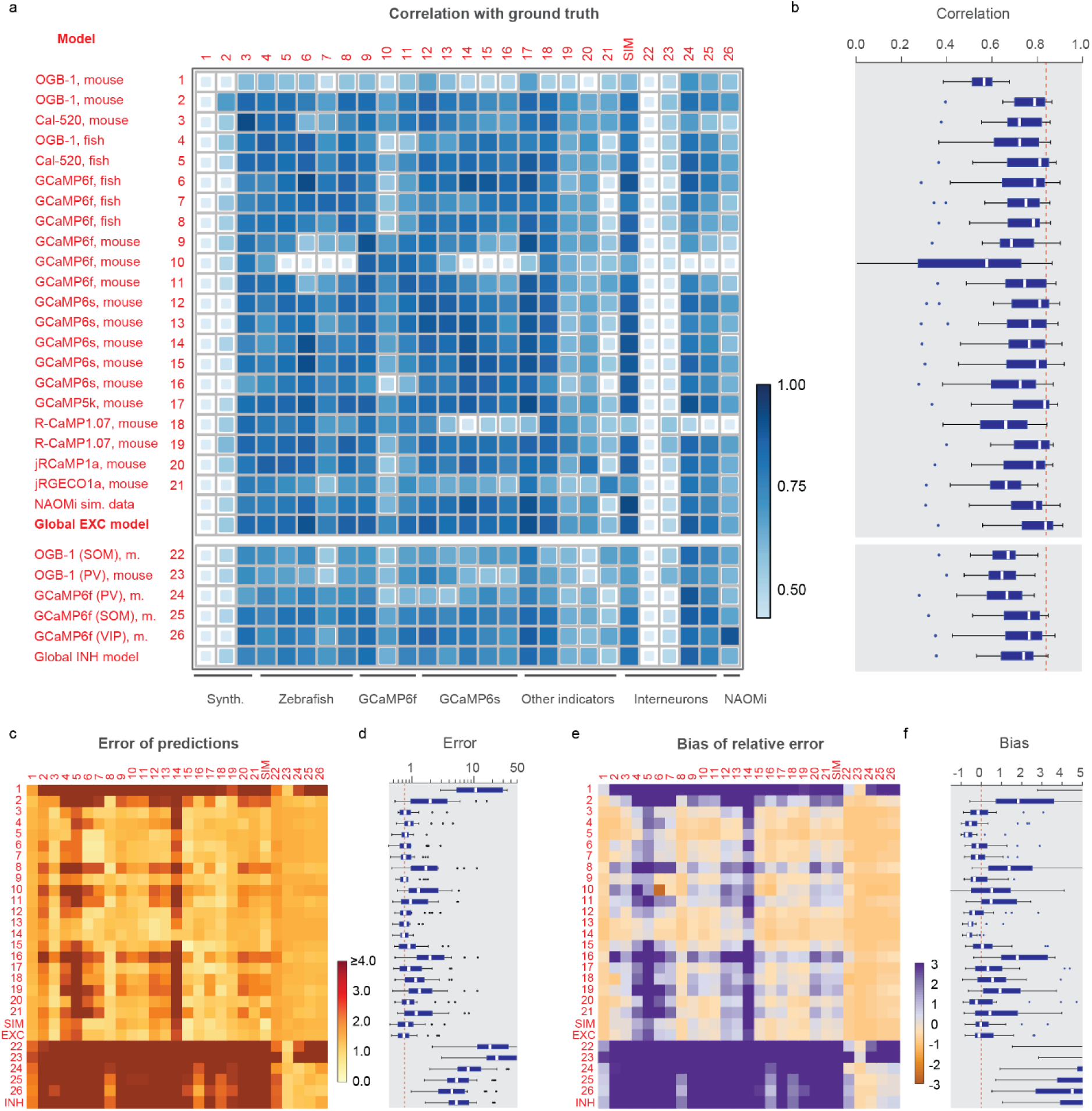
Generalization across datasets. The network was trained on a given dataset (indicated by the row number) and tested on each other ground truth dataset (column). Diagonal values correspond to metrics shown in Fig. 3e. „NAOMi“ is a model trained on simulated GCaMP6f data based on Charles et al. (2019). Rows 21-24 are networks trained on datasets with interneurons. „global EXC model“ and „global INH model“ are globally trained on all excitatory or inhibitory datasets (except datasets #01 and and the respective test dataset). **a,** Correlation of predictions with the ground truth. The size and color of the squares scale with correlation. **b,** Distribution of the performance of each trained network (row) across all datasets. The dashed line highlights the median of the best-performing model (‘global EXC model’). **c-d,** Relative error of predictions compared to the ground truth. The dashed line in (d) highlights the median of the best-performing mode (‘global EXC model’). **e-f,** Relative bias of predictions compared to the ground truth. All datasets were re-sampled at a frame rate of 7.5 Hz, with a standardized noise level of 2.

The entries of the matrix in Fig. 3 remained highly similar when parameters such as the resampling rate, temporal smoothing of the ground truth or the noise level were modified (Fig. S13). Interestingly, models trained on datasets of excitatory neurons (DS#01-21) also produced high quality-predictions of spike rate variations in interneurons (DS#22-26) (Fig. 3a,b). However, the separate analysis of error and bias revealed that absolute spike rates were substantially underestimated (Fig. 3c-f).^44^

Near-maximal correlation for a given dataset was often achieved by multiple models (Fig. 3a). In some datasets, the highest correlation was even achieved when the model was trained on ground truth from another dataset, rather than from the same dataset. Interestingly, the performance of training/testing combinations showed no obvious clustering related to indicator type (*e.g.,* genetically encoded vs. organic indicators) or species (zebrafish vs. mouse). An attempt to explain the mutual predictability of datasets by more refined statistical dataset descriptors such as the mean spike rate or decay times was not very successful (Fig. S14). It is therefore not obvious how to select an optimal training dataset to predict spike rates for an unseen dataset.

To optimize dataset selection and network training for practical applications, we tested an alternative and simpler approach by training a model on all excitatory datasets except DS#01, hence called the ‘global EXC model’. We found that this global model performed better than all other models in cross-dataset tests (Fig. 3a-f; test dataset was always excluded from training data), not only due to the size but also due to the diversity of the training set (Fig. S15). Compared to randomly selecting a single dataset with excitatory neurons for training, correlations were increased by 0.05±0.05, errors were reduced by 0.05±0.05, and absolute biases were reduced by 0.25±0.90 (median ± s.d.). In addition, the global EXC model performed better than any of the 21 single models in all cross-dataset tests (p < 0.001 for all comparisons, paired signed-rank test). Compared to predictions across neurons within the same dataset (Fig. S9; diagonal elements in Fig. 3), the correlations resulting from the global EXC model were decreased by 0.02±0.04 (p = 0.04, Wilcoxon signed-rank test), errors were increased by 0.33±0.53 (p = 0.01), while the absolute bias was slightly decreased (0.40±0.40, p = 0.002). Hence, using dataset-specific ground truth can yield performance significantly better than the global EXC model. In the absence of such specific calibration data, however, training the algorithm with all available data is a simple and effective strategy to generate a model that generalizes robustly to unseen datasets.

Not surprisingly, a global INH model trained on all interneuron datasets generalized less well than the EXC model (Fig. 3). Indeed it was not more successful than the EXC model in predicting interneuron activity, most likely because the diversity of interneuron types is higher and because the amount of available ground truth was lower.

We also trained a model on a large artificial dataset (250 neurons) that was generated using the calcium imaging simulation environment NAOMi^45^ (Methods). The model performed well but lower than the global EXC model (correlation reduced by 0.05±0.04, p=0.0003; error slightly increased by 0.06±0.22, p=0.0006; bias not significantly changed, p=0.67; Fig. 3). We hypothesize that some relevant sources of variability at the neuronal level (*e.g.,* variable decay times, transient shapes and non-linearities) are captured by experimental ground truth but not by simulated ground truth recordings. For new indicator types, where biophysical parameters are known but experimental ground truth is not available, ground truth generated by NAOMi could nonetheless be a useful way to train supervised spike inference algorithms.

### Comparison with existing methods

To benchmark the performance of CASCADE, we compared it to five other model-based methods. In summary, we found that CASCADE performed better than all other algorithms across datasets, across noise levels, and for different temporal precisions. Moreover, CASCADE showed a less pronounced bias towards underestimating high spike rates.

We compared CASCADE to the fast online deconvolution procedure OASIS with two distinct implementations in CaImAn and Suite2p^14, 42, 46^, to the discrete change-point detection algorithm by Jewell and Witten^47^ (here referred to as Jewell&Witten), and to two more complex algorithms, Peeling and MLSpike. Peeling uses iterative template-subtraction to infer discrete spikes^10^. MLSpike was chosen because it outperformed various other methods in previous applications^11, 15^. Although model-based methods are, in principle, non-supervised, several parameters need to be tuned to achieve maximal performance on a given dataset^12^. To avoid sub-optimally tuned algorithms and to make the comparison with CASCADE as fair as possible, we used extensive grid searches to optimize parameter tuning of each algorithm-dataset combination (Methods; see Table 2 for the best model parameters for each dataset as a function of noise). This procedure allowed us to minimize the same loss function for all algorithms (mean squared error between ground truth and the inferred spike rate), using grid search for model-based approaches and backpropagation for CASCADE. Importantly, the neuron used for testing was always omitted during the training/fitting period (leave-one-out strategy). We refer to these models as “tuned” for specific datasets, as opposed to CASCADE’s “global EXC model” that was trained on other datasets (Fig. 3). The Peeling and Jewell&Witten algorithms infer discrete spikes rather than spike rates, which may result in a slight disadvantage. To convert their output to continuous rates, predicted spikes were convolved with a Gaussian kernel of a width that minimized the mean squared error.

**Table 2.** (supplementary material) Optimal parameters for the model-based spike detection algorithms. . Values were found for each dataset separately using a grid search over the indicated parameters (Methods). Standardized noise level ν in % · *Hz*^−1/2^. Black columns contain optimal parameters when optimizing for the mean squared error of predictions with respect to ground truth (frame rate 7.5 Hz, smoothing with σ=0.2 s). Continued on next page.

| NOISE<br>LEVEL 2 | MLSpike |  | Peeling |  | CalmAn | Suite2p |  | Jewell |  |
| --- | --- | --- | --- | --- | --- | --- | --- | --- | --- |
|  | tau | amplitude | tau | amplitude | gamma | sig_baseline | gamma | penalty | gamma |
| DS01 | 0.1000 | 0.3500 | 0.6360 | 7.2730 | 0.0200 | 0.2500 | 0.0250 | 0.0040 | 0.4730 |
| DS02 | 1.9690 | 0.3500 | 0.2500 | 31.5620 | 0.0200 | 32.5000 | 0.2190 | 0.0200 | 0.7060 |
| DS04 | 1.5000 | 0.1910 | 1.0170 | 16.0000 | 0.0250 | 20.6670 | 0.1000 | 0.0500 | 0.7000 |
| DS05 | 4.0000 | 0.0100 | 2.3750 | 35.0000 | 0.7400 | 33.7500 | 0.6750 | 0.0840 | 0.8500 |
| DS06 | 2.0000 | 0.0900 | 2.3750 | 31.8750 | 0.8030 | 0.2500 | 0.6750 | 0.0070 | 0.9500 |
| DS07 | 2.0000 | 0.2420 | 2.1000 | 35.0000 | 0.8180 | 0.2500 | 0.6900 | 0.0160 | 0.9200 |
| DS08 | 1.2220 | 0.0300 | 2.0000 | 8.3330 | 0.1890 | 0.2500 | 0.0920 | 0.0040 | 0.6560 |
| DS09 | 1.0000 | 0.2500 | 0.9320 | 28.6360 | 0.4180 | 4.3180 | 0.2000 | 0.0500 | 0.5090 |
| DS10 | 0.7500 | 0.1610 | 0.8040 | 35.0000 | 0.5540 | 0.2500 | 0.2830 | 0.0100 | 0.8000 |
| DS11 | 0.7600 | 0.1300 | 0.5300 | 29.4000 | 0.1130 | 0.3100 | 0.1000 | 0.0150 | 0.7180 |
| DS12 | 3.6670 | 0.3500 | 2.4170 | 35.0000 | 0.7730 | 2.1250 | 0.5500 | 0.0400 | 0.8000 |
| DS13 | 2.0000 | 0.3500 | 2.9810 | 35.0000 | 0.8170 | 0.2500 | 0.7920 | 0.0200 | 0.9000 |
| DS14 | 1.0000 | 0.2500 | 0.9320 | 28.6360 | 0.4180 | 4.3180 | 0.2000 | 0.0500 | 0.5090 |
| DS15 | 1.5000 | 0.0300 | 2.4440 | 15.5560 | 0.6220 | 0.2500 | 0.3000 | 0.0070 | 0.9000 |
| DS16 | 0.7500 | 0.0100 | 1.1110 | 14.4440 | 0.0420 | 0.2500 | 0.0500 | 0.0040 | 0.7110 |
| DS17 | 0.8330 | 0.0830 | 1.5000 | 20.0000 | 0.5040 | 0.3610 | 0.2000 | 0.0100 | 0.8000 |
| DS18 | 1.0630 | 0.0700 | 1.1250 | 18.7500 | 0.3200 | 0.2500 | 0.1810 | 0.0040 | 0.9130 |
| DS19 | 1.2500 | 0.1060 | 0.5280 | 18.8890 | 0.0200 | 2.5000 | 0.0250 | 0.0190 | 0.5890 |
| DS20 | 2.5000 | 0.0860 | 1.9440 | 10.5560 | 0.1620 | 4.7220 | 0.1220 | 0.0320 | 0.7890 |
| DS21 | 0.4050 | 0.2460 | 0.5000 | 35.0000 | 0.0200 | 4.0910 | 0.0390 | 0.0550 | 0.2320 |
| DS22 | 1.6200 | 0.3500 | 0.4000 | 10.0000 | 0.1880 | 4.6000 | 0.0250 | 0.0060 | 0.2400 |
| DS23 | 2.1570 | 0.1790 | 0.2860 | 2.5000 | 0.0200 | 0.2500 | 0.0250 | 0.0010 | 0.1000 |
| DS24 | 0.2000 | 0.0130 | 2.0390 | 2.5000 | 0.0200 | 0.2500 | 0.0250 | 0.0030 | 0.1810 |
| DS25 | 0.2000 | 0.0290 | 1.8530 | 8.8240 | 0.0690 | 0.2500 | 0.0290 | 0.0030 | 0.5650 |
| DS26 | 0.2000 | 0.0550 | 1.1250 | 9.3750 | 0.0200 | 0.2500 | 0.0250 | 0.0040 | 0.4000 |

| NOISE<br>LEVEL 9 | MLSpike |  | Peeling |  | CalmAn | Suite2p |  | Jewell |  |
| --- | --- | --- | --- | --- | --- | --- | --- | --- | --- |
|  | tau | amplitude | tau | amplitude | gamma | sig_baseline | gamma | penalty | gamma |
| DS01 | 0.1000 | 0.3500 | 0.3640 | 19.0910 | 0.2450 | 5.4550 | 0.0250 | 0.0250 | 0.1000 |
| DS02 | 0.1130 | 0.3500 | 4.9380 | 35.0000 | 0.7600 | 50.0000 | 2.8940 | 0.2500 | 0.9000 |
| DS04 | 2.0000 | 0.3500 | 0.5000 | 29.3330 | 0.0200 | 30.0000 | 0.5130 | 0.2500 | 0.3500 |
| DS05 | 2.6250 | 0.2650 | 2.2500 | 35.0000 | 0.7150 | 45.0000 | 1.2250 | 0.5630 | 0.7250 |
| DS06 | 1.9380 | 0.0300 | 2.0000 | 34.3750 | 0.7830 | 4.3750 | 0.8000 | 0.1190 | 0.8000 |
| DS07 | 2.9000 | 0.1580 | 2.5000 | 35.0000 | 0.8160 | 3.0000 | 1.0300 | 0.2500 | 0.8000 |
| DS08 | 1.7220 | 0.0830 | 2.2780 | 8.3330 | 0.1760 | 0.3610 | 0.1830 | 0.0300 | 0.1170 |
| DS09 | 1.4770 | 0.3500 | 0.8860 | 33.1820 | 0.5380 | 50.0000 | 0.6450 | 0.2500 | 0.5000 |
| DS10 | 0.7500 | 0.1600 | 1.0000 | 35.0000 | 0.5530 | 3.4780 | 0.3740 | 0.1000 | 0.2480 |
| DS11 | 0.5000 | 0.2880 | 1.0000 | 20.0000 | 0.1460 | 5.2000 | 0.2000 | 0.1000 | 0.1900 |
| DS12 | 1.2920 | 0.3500 | 2.4170 | 35.0000 | 0.7770 | 3.8330 | 1.1330 | 0.2250 | 0.7000 |
| DS13 | 1.5000 | 0.2270 | 3.0000 | 35.0000 | 0.8450 | 0.9620 | 1.3270 | 0.1810 | 0.7460 |
| DS14 | 1.4770 | 0.3500 | 0.8860 | 33.1820 | 0.5380 | 50.0000 | 0.6450 | 0.2500 | 0.5000 |
| DS15 | 1.5000 | 0.0300 | 2.3890 | 15.5560 | 0.5910 | 1.5000 | 0.3890 | 0.1000 | 0.6670 |
| DS16 | 1.5000 | 0.0300 | 1.0000 | 13.0560 | 0.0200 | 0.3060 | 0.1110 | 0.0250 | 0.1610 |
| DS17 | 0.9720 | 0.0720 | 1.7220 | 19.4440 | 0.5000 | 17.2220 | 0.2560 | 0.1000 | 0.5000 |
| DS18 | 1.0630 | 0.3200 | 1.6250 | 14.3750 | 0.3200 | 1.1250 | 0.2880 | 0.0590 | 0.4630 |
| DS19 | 1.8890 | 0.3480 | 0.5000 | 18.8890 | 0.0200 | 18.5710 | 0.2140 | 0.1000 | 0.2640 |
| DS20 | 2.5000 | 0.2260 | 2.0560 | 15.0000 | 0.1130 | 18.8890 | 0.6000 | 0.2500 | 0.5670 |
| DS21 | 0.4320 | 0.3010 | 0.5000 | 33.1820 | 0.0200 | 29.0910 | 0.0950 | 0.1140 | 0.1000 |
| DS22 | 0.1000 | 0.3500 | 3.2500 | 33.0000 | 0.3200 | 34.0000 | 1.5400 | 0.1500 | 0.5100 |
| DS23 | 0.1000 | 0.3500 | 2.4290 | 3.5710 | 0.1290 | 0.2500 | 0.0290 | 0.0000 | 0.2290 |
| DS24 | 0.3500 | 0.0300 | 2.0770 | 2.5000 | 0.0200 | 0.2500 | 0.0250 | 0.0030 | 0.1000 |
| DS25 | 0.3590 | 0.0310 | 1.5880 | 9.4120 | 0.0550 | 0.6320 | 0.0570 | 0.0200 | 0.1000 |
| DS26 | 0.1000 | 0.0700 | 1.0000 | 10.0000 | 0.0200 | 0.5630 | 0.0250 | 0.0100 | 0.1000 |

The tested algorithms showed systematic differences in performance (Fig. 4a, Fig. S16). A quantitative comparison across all datasets for a fixed noise level revealed that performance varied strongly across ground truth datasets, single neurons, and algorithms (Fig. 4b, Fig. S17). Neurons that could be predicted well by one algorithm could often also be predicted well by other algorithms, in particular with respect to errors and biases (Fig. S17), suggesting that outlier neurons within datasets exhibit unusual properties that lead to biased predictions (cf. Fig. S9). The tuned CASCADE model and CASCADE’s global EXC model produced good predictions for the broadest set of neurons across datasets. High-level performance of the model-based algorithms was observed in fewer datasets. For example, in multiple neurons from diverse datasets, the performance of MLSpike was lower than that of CASCADE (Fig. 4b; datasets #7-8 [GCaMP6f in fish], #15-16 [GCaMP6s in V1], #24-26 [GCaMP6f in interneurons]). These datasets had relatively high (Table 1) and slowly changing spike rates rather than discrete bursts. Interestingly, the Peeling algorithm performed relatively well on some of these datasets. To more directly compare the performances across neurons with CASCADE, we calculated the difference in correlation achieved by CASCADE and other algorithms for each neuron. The resulting distributions (Fig. 4c) show that CASCADE yields better inferences for the majority of neurons across all compared algorithms (p<10^-10^ for all comparisons with other algorithms; p=0.068 when compared with CASCADE’s global EXC model; paired Wilcoxon signed-rank test).

**Figure 4.**
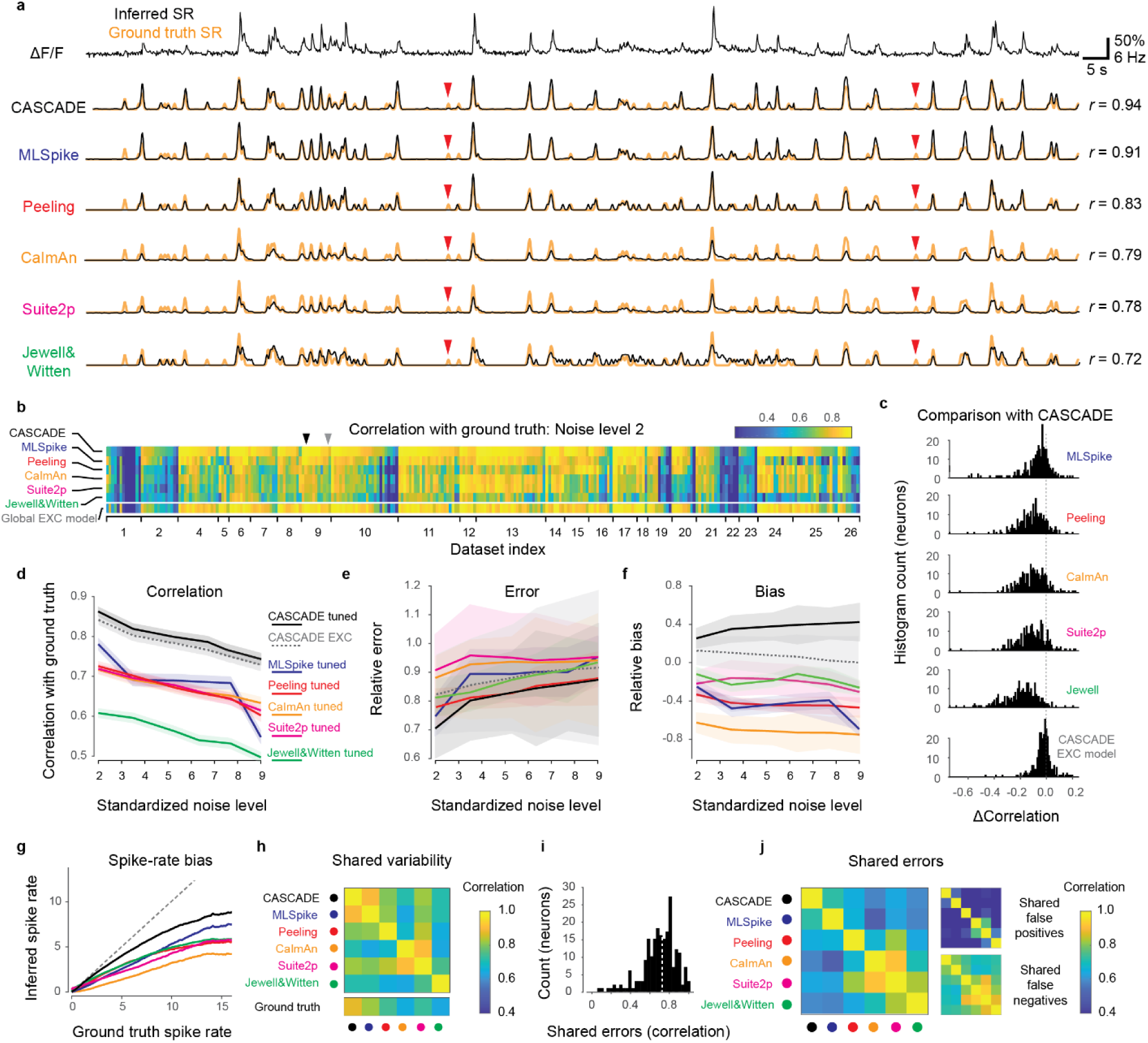
Comparison with model-based algorithms. **a,** Example calcium imaging recording and corresponding predictions from the deep-learning based method (CASCADE) and five model-based algorithms (MLSpike, Peeling, CaImAn, Suite2p, Jewell&Witten). Respective predictions are in black, ground truth in orange. *r* indicates correlation of predictions with ground truth. Clear false negative detections are labeled with red arrowheads. **b**, Heat map of the performance (correlation) of each algorithm for each dataset and neuron, calculated at standardized noise level 2 % Hz^-1/2^. All algorithms, except for CASCADE‘s global EXC model (cf. Fig. 3), were tuned to the respective dataset by the mean squared error between ground truth and inferred spike rate. Arrowheads highlight the example neurons shown in Fig. 4a (black) and Fig. S16 (grey). **c,** Direct comparison of performance (b) between CASCADE and other algorithms on a single-neuron basis. The difference in performance (correlation) is shown as a histogram across all neurons. ‘Global EXC model‘ as defined in Fig. 3. **d-f,** Comparison of correlation, error and bias across all algorithms and noise levels. ν in units of standardized noise, % · *Hz*^−1/2^. Solid/dashed lines indicate the mean across all neurons, shaded areas represent the SEM. **g**, Spiking activity in 2 s-bins, ground truth vs. Predictions. Lines indicate medians across distributions. Algorithms are color-coded as before. Underlying distributions are shown in Fig. S16. The unity relationship is shown as dashed line. **h**, Variability shared across algorithms, measured by the correlation between predictions. **i**, Histogram of error shared between CASCADE and MLSpike, quantified as the correlation between the unexplained variances for each neuron. Dashed line indicates the median. **j**, Shared median errors as illustrated in (i) for all pairs of algorithms. The smaller matrices to the right break the shared errors down into false positives and false negatives. All quantifications were performed with ground truth datasets resampled at 7.5 Hz with a noise level of 2 unless otherwise indicated. Dataset #03 was omitted for all comparisons in Fig. 4 since the short recordings (<10 s) could not be processed by all algorithms.

We found that the performance of CASCADE was consistently better across different recording conditions. First, based on the finding that noise levels affect spike inference more strongly than other parameters (Fig. S13), we repeated the benchmarking in Fig. 4b across multiple noise levels and found that the performance ranking across algorithms was largely maintained (Fig. 4d), with the global EXC model achieving performance close to the tuned CASCADE model (significant difference: p=0.039, signed-rank test), followed by MLSpike, Peeling, Suite2p and CaImAn with similar performances, then followed by Jewell&Witten (p<10^-10^ for all algorithms). Although the error computed from CASCADE’s predictions was significantly lower than for most other algorithms (p<0.005 for Jewell&Witten, Suite2p, MLSpike and CASCADE’s global EXC model; p<0.01 for CaImAn; but p>0.05 for Peeling; paired Wilcoxon signed-rank), variability was high and relative effect sizes were low (Fig. 4e). Therefore, errors are not a very sensitive readout of performance. Finally, biases of predictions were negative (indicating underestimates of true spike rates) for all tuned model-based algorithms except for the tuned CASCADE model (Fig. 4f). CASCADE’s global EXC model exhibited the smallest overall bias.

We further found that all algorithms systematically underestimated spike rates as spike rates increased, but this effect was smallest for CASCADE. To visualize these results, we plotted the number of spikes for ground truth and predictions within each 2-s time bin (Fig. S18) and extracted the median lines of these distributions (Fig. 4g). An underestimate of high spike rates may be expected since periods of high spike rates are less frequent. False positive predictions of high spike rates would thus lead to larger performance drops than false negative omissions of rare events.

For spike inference evaluated at higher temporal precision, the performance (correlation with ground truth) dropped for all algorithms, but this effect was more modest for CASCADE than for all model-based algorithms (Fig. S19). We trained all algorithms to a ground truth that was smoothed in time to a variable degree (Gaussian smoothing kernel between σ = 0 ms and σ = 333 ms; default: σ = 200 ms). Example predictions highlight that several algorithms make impressive predictions also under these more difficult conditions (Fig. S19a), but some algorithms, in particular those based on discrete events (Peeling, Jewell&Witten) were not able to include graded certainties about spike times and therefore performed less well (Fig. S19b). However, also the performance of MLSpike, CaImAn and Suite2p dropped faster than the performance of CASCADE when spike rates were evaluated with increasing temporal precision (Fig. S19c).

Overall, predictions across algorithms were not only similar on the level of neurons (Fig. 4b, Fig. S17), but also exhibited highly correlated predictions and errors over the time course of single neurons (Fig. 4h-j). The shared variability, measured as the median correlation between predictions, was particularly high for the two closely related algorithms, Suite2p and CaImAn, but also high between CASCADE and MLSpike. Indeed, the correlation between CASCADE and MLSpike was as high as the correlation between CASCADE and the ground truth (Fig. 4h, bottom). To better understand these similarities, we explored false predictions shared by algorithms. To this end, we computed the similarities (correlation) of the unexplained, residual variances across algorithms. These shared errors were relatively prominent across algorithms (Fig. 4i,j). In particular, errors made by CaImAn, Suite2p, Peeling and Jewell&Witten were often correlated, but CASCADE and MLSpike also shared a relatively large fraction of unexplained variance. We further divided the unexplained variance in false positives (predictions were higher than the ground truth) and false negatives (predictions were lower than the ground truth). False negatives but not false positives were highly correlated across most algorithms, with the exception of Suite2p and CaImAn, which also shared false positives (Fig. 4j, right). Shared false negatives are clearly visible in typical predictions (red arrows in Fig. 4a, and more prominently in Fig. S16). Together, these analyses show highly similar predictions and similar missed spike events across algorithms.

Finally, we compared practical aspects arising during the application of different algorithms. With respect to processing speed we found that CASCADE (based on a GPU), CaImAn, and Jewell&Witten performed similarly fast (200k-300k samples/s). They were only outperformed by Suite2p (more than 5M samples/s). Peeling (ca. 5k samples/s) and in particular MLSpike (ca. 0.8k samples/s) were much slower, owing to their more complex fitting procedure. For optimization, CASCADE uses backpropagation, which is almost equally fast as inference, resulting in a total training time of <10 min for a typical ground truth dataset with 5M data points and a realistic number of 20 iterations (epochs) through the dataset. For model-based algorithms, extensive grid searches across parameters must be performed. We usually performed grid search in a 2D parameter space with 100-500 parameter combinations, which is feasible within minutes for Suite2p, CaImAn and Jewell&Witten. For Peeling and MLSpike, this procedure would take several days for a single model. In our analyses, we therefore reduced the number of training samples for MLSpike and Peeling such that the respective grid searches lasted approximately 2 hours per model for MLSpike. Furthermore, we found that best fit parameters for model-based approaches changed systematically with different noise levels, suggesting that new models have to be fit for each noise level (Table 2, supplementary material), an effect that was more pronounced for algorithms that do not use the noise level as an input (Suite2p and Jewell&Witten). As another possible drawback of some model-based algorithms, we found that inferred spike rates were often temporally shifted to later time points, and this delay was variable across datasets (0.16±0.14 s for MLSpike, 0.03±0.09 s for Peeling, 0.31±0.23 s for CaImAn, 0.29±0.22 s for Suite2p and 0.27±0.19 s for Jewell&Witten; mean delay ± s.d. across datasets). We corrected these shifts for all analyses presented here. However, using algorithms without this correction could result in misleading interpretations of the temporal dynamics of the inferred spike rates. This can be prevented by using an algorithm that does not induce strong delays (*e.g.,* Peeling) or a model-free algorithm that directly learns the best delay from ground truth (CASCADE). Together, these aspects reflect that, unlike model-based algorithms, CASCADE can make use of ground truth datasets in an efficient and natural way.

### Application to large-scale population calcium imaging datasets

A transformation of calcium signals into estimates of spike rates may be desired for multiple reasons. First, the reconstruction of spike rates can recover fast temporal structure in neuronal activity patterns that is obscured by slower calcium signals^4, 5^. Second, calcium signals contain shot noise and potentially other forms of noise that are unrelated to neuronal activity. A method that infers spike rates while ignoring noise can therefore de-noise activity measurements without the detrimental effects of over-expressed indicators^1, 48^ and without compromising temporal resolution. Third, while calcium signals usually represent relative changes in activity, spike rates provide absolute activity measurements that can be compared more directly across experiments. With these potential goals in mind, we applied CASCADE to different large-scale calcium imaging datasets.

In a brain explant preparation of adult zebrafish^23^, we measured odor-evoked activity in the posterior part of telencephalic area Dp (pDp), the homolog of piriform cortex, using OGB-1. Multi-plane two-photon imaging^49^ was performed with the same recording conditions as in DS#04 at a noise level of 2.36±0.97 (% · *Hz*^−1/2^; median ± s.d.) across 1,126 neurons. Under these conditions, predictions are expected to be highly accurate (Fig. S9a,e; correlation to ground truth: 0.87±0.06 for a standardized noise level of 2, median ± s.d.; Gaussian smoothing of the ground truth with σ=0.2 s). Consistent with previous electrophysiological recordings^50^, spiking activity estimated by CASCADE with a model trained on DS#04 was sparse (0.6±1.1 spikes during the initial 2.5 s of the odor response; mean ± s.d.; Fig. 5a) and variable across neurons (Fig. 5b), with a clear difference between the anatomically distinct dorsal and ventral regions of pDp (0.07±0.11 vs. 0.21±0.11 Hz; entire recording).

**Figure 5.**
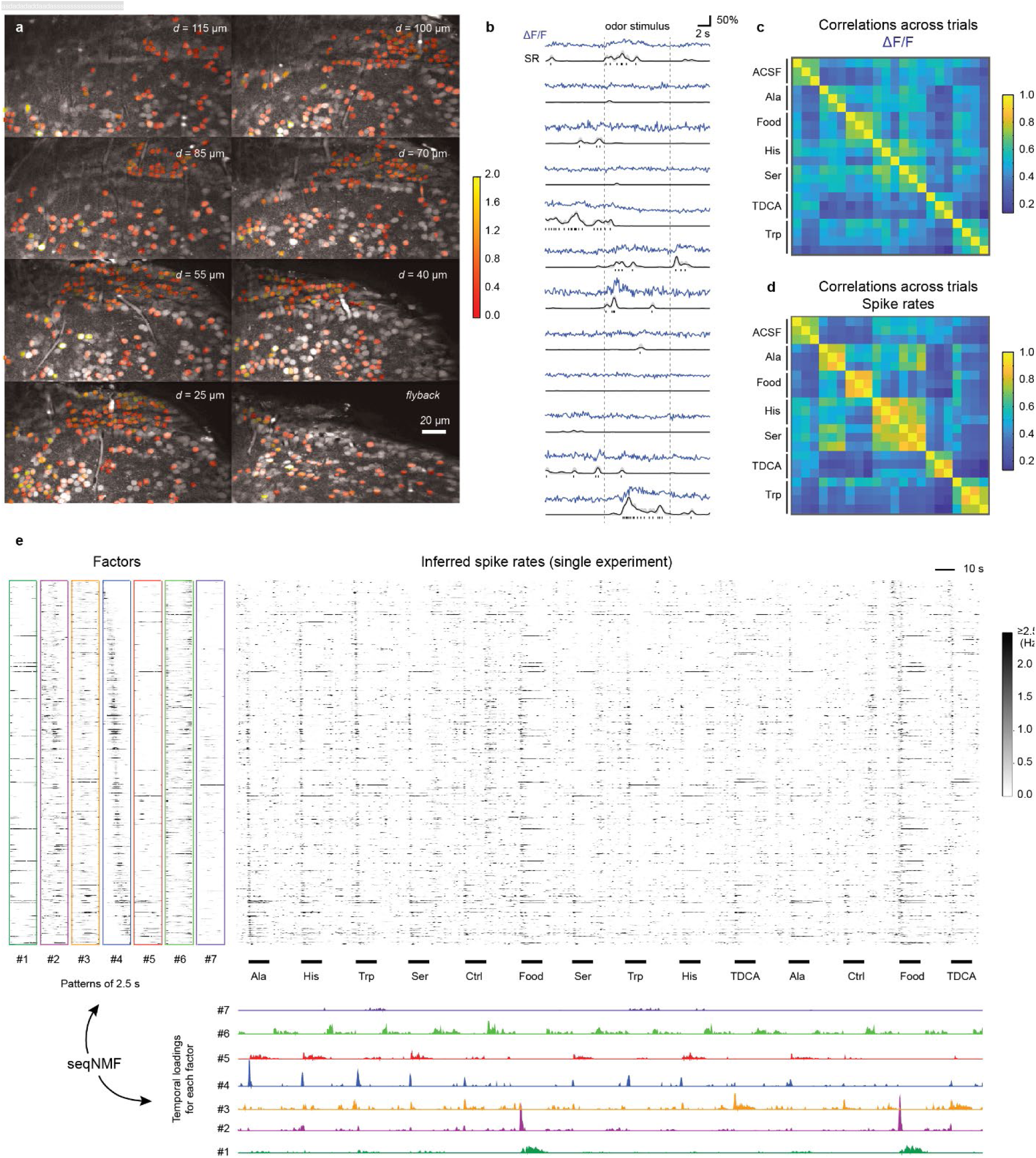
Inference of spiking activity with CASCADE from population calcium imaging across >1100 neurons in adult zebrafish. **a,** Multiple planes were imaged simultaneously. The ROIs are colored with the average number of inferred stimulus-evoked spikes (colorbar). Non-active neurons were left uncolored. **b,** Randomly selected examples of calcium traces (ΔF/F, blue), inferred spike rates (SR, black) and inferred discrete spikes, highlighting the de-noising through spike inference. **c,** Correlation of odor-evoked responses across trials, based on ΔF/F data during the initial 2.5 s of the odor response. **d,** Correlation of odor-evoked responses across trials, based on inferred spiking probabilities. **e,** Unsupervised detection of sequential factors (left) and their temporal ‘loading’ (bottom), shown together with the inferred spiking probabilities (center) across a subset of stimulus repetitions. The temporal loadings indicate when a given factor becomes active. All neurons were ordered according to highest activity in pattern #4, highlighting the sequential activity pattern that is evoked by stimuli at multiple times.

The comparison of ΔF/F signals and inferred spike rates showed that CASCADE detected phases of activity but effectively suppressed small irregular fluctuations in activity traces, indicating that spike inference suppressed noise. Consistent with this interpretation, spike inference by CASCADE increased the correlation between time-averaged population activity patterns evoked by the same odor stimuli in different trials (Fig. 5c,d).

Previous studies showed that odor-evoked population activity in pDp is dynamic^50, 51^ but the fine temporal structure has not been explored in detail. We applied an unsupervised non-negative matrix factorization method for sequence detection (seqNMF, ref. ^52^) to the inferred spike rate patterns to identify recurring short (2.5 s) sequences of population activity (hereafter referred to as factors) embedded in the overall population activity. This analysis required a high effective temporal resolution because factors exhibited rich temporal structure on a sub-second timescale (Fig. 5e). We found multiple factors that occurred with high precision and in a stimulus-specific manner at different phases of the odor response. For example, factors #2 and #4 in Fig. 5e were transient and associated with response onset, factor #5 persisted during odor presentation, and factor #6 was activated after stimulus offset (Fig. 5e). Odor-evoked population activity in pDp therefore shows complex dynamics on timescales that cannot be resolved without temporal deconvolution of calcium signals. The transformation of calcium signals into spike rate estimates by CASCADE therefore provides interesting opportunities to use calcium imaging for the analysis of fast network dynamics.

We next analyzed the Allen Brain Observatory Visual Coding dataset, comprising >400 experiments in mice with transgenic GCaMP6f expression, each consisting of approximately 100-200 neurons recorded at very low noise levels (0.94±0.25 % · *Hz*^−1/2^; mean ± s.d.; Fig. 6a)^53^. Using the global EXC model of CASCADE we estimated the absolute spike rates across all neurons for different transgenic lines (Fig. 6b; Fig. S20; Gaussian smoothing of the ground truth with σ=0.05 s). Spike rates across all 38,466 neurons were well described by a lognormal distribution centered around 0.1-0.2 Hz (Fig. 6c). Given the sampling rate (30 Hz) and noise level of this dataset we expect a correlation of 0.89±0.18, an error of 0.70±0.96 and a bias of 0.27±1.00 (median ± s.d. across neurons), based on our previous cross-dataset comparisons which also covered ground truth for transgenic lines included in this population imaging dataset (Fig. 3). Since we could not broadly test the generalization across a large set of interneuron datasets (Fig. 3), we did not include interneuron experiments in our analysis. Spike rates varied systematically across cortical layers, with highest activity in layer 5 (Fig. 6d,e), across transgenic lines (Fig. 6d) and across stimuli presented, with highest activation during naturalistic stimuli (natural scenes or movies; Fig. 6e). These results provide a comprehensive description of neuronal activity in the visual system of the mouse and reveal systematic differences in neuronal activity across cell types, brain areas, cortical layers, and stimuli.

**Figure 6.**
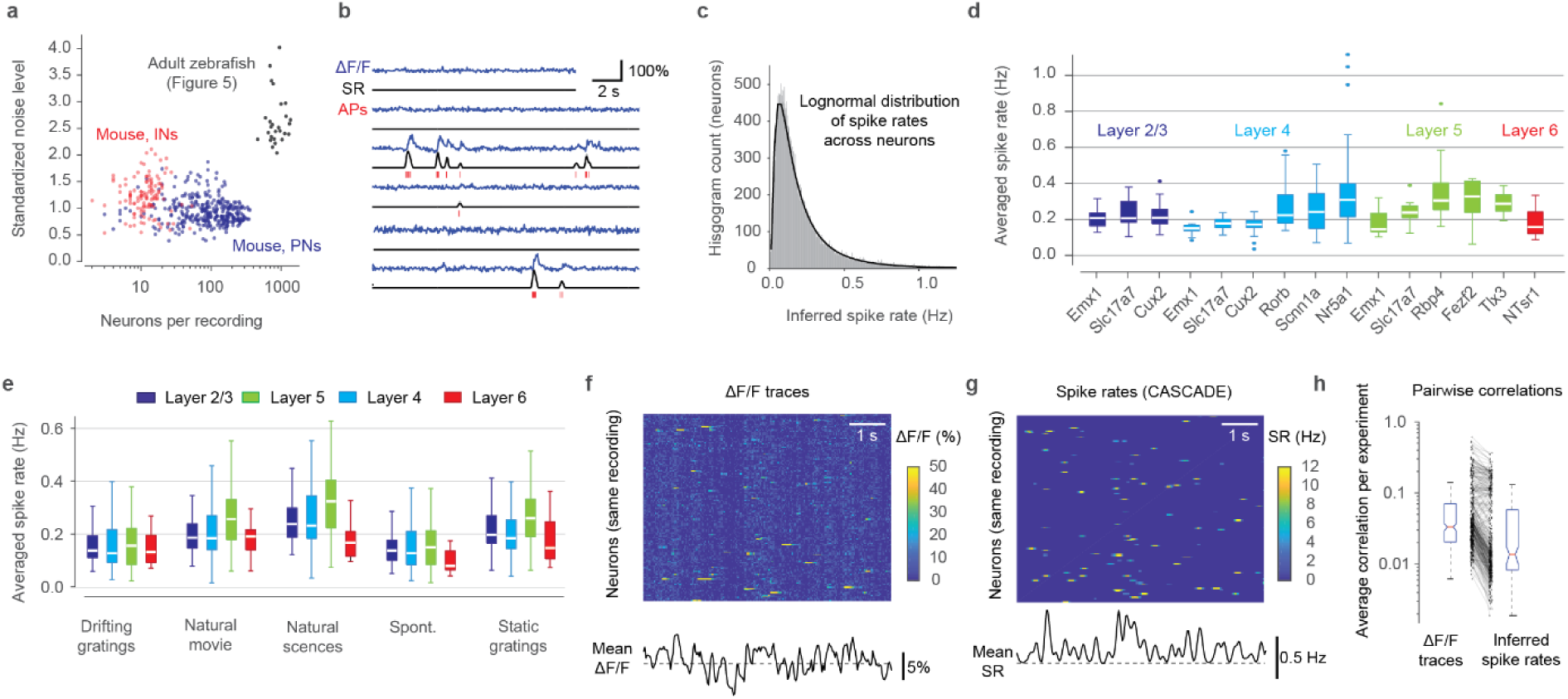
Inference of spiking activity with CASCADE for the Allen Brain Observatory dataset in mice. **a,** Number of recorded neurons vs. standardized noise levels (in % · *Hz*^−1/2^) for all experiments from dataset from excitatory (blue) and inhibitory (red) datasets; population imaging datasets in zebrafish (Fig. 5) in black for comparison. **b,** Example predictions from calcium data (blue). Discrete inferred spikes are shown in red below the inferred spike rates (black). See Fig. S20 for more examples. **c,** Spike rates across the entire population are well described by a log-normal distribution (black fit). n = 38,466 neurons**. d,** Inferred spike rates across all neurons for recordings in different layers (colors) and for different transgenic driver lines of excitatory neurons. Each underlying data point is the mean spike rate across an experiment. Total of 336 experiments. **e,** Average spike rates for different stimulus conditions (x-labels) across layers (colors). Each data point is the mean spike rate across one experiment. **f,** Excerpt of raw ΔF/F traces of a subset of neurons of a single experiment (L2/3-Slc17a7, experiment ID ‘652989705’). Correlated noise is visible as vertical striping patterns. **g,** Same as (f), but with inferred spike rates. **h,** Average correlation between neuron pairs within an experiment, computed from raw ΔF/F traces (left) and inferred spike rates (right).

Raw ΔF/F often exhibited correlated noise, visible as a vertical striping in matrix plots, which was small for individual neurons but tended to dominate the mean ΔF/F across neurons, possibly due to technical noise or neuropil signal (Fig. 6f). CASCADE effectively eliminated these artifacts (Fig. 6g). As a consequence, correlations between activity traces of different neurons were reduced across all experiments by 38±43% (mean ± s.d.; Fig. 6h; p < 10^-15^, paired signed-rank test). Many analyses of neuronal population activity require accurate measurements of pairwise neuronal correlations^53–56^. Noise suppression by spike inference can therefore help to make these analyses more reliable.

Together, these examples illustrate how calibrated spike inference by CASCADE can be applied to perform comprehensive analyses of neuronal activity, to identify complex temporal structure in neuronal population dynamics, and to remove shot noise and other noise from calcium imaging data.

### A user-friendly toolbox for spike inference

The deployment of spike inference tools often suffers from practical problems. First, the difficulty to set up a computational pipeline might prevent wide-spread use. To address this problem, we generated a cloud-based solution using Colab Notebooks that can be applied without local installations. We also set up a well-documented Github repository (https://github.com/HelmchenLabSoftware/Cascade) containing ground truth datasets, pre-trained models, notebooks and demo scripts that can be easily integrated into existing analysis pipelines such as CaImAn, SIMA or Suite2P^42, 46, 57^. Since the algorithm works on regular laptops and workstations without GPU support, the main installation difficulties of typical deep learning applications are circumvented.

In a typical workflow, first the noise level for each neuron in a calcium imaging dataset is determined. Then, a pre-trained model that has been trained on noise-matched resampled ground truth data is loaded from an online model library and applied to the ΔF/F data without any need to adjust parameters. In addition, CASCADE can be easily modified and retrained to address additional specific needs, for example, more complex loss functions^22^ or a modified architecture. Also, the resampled ground truth can be adapted directly if desired. For example, we used a Gaussian kernel to smooth the ground truth spike rate, but this standard procedure can be disadvantageous to precisely determine the onset timing of discrete events. In CASCADE, it is simple to replace the Gaussian kernel by a causal smoothing kernel to circumvent this problem (Fig. S21).

A second problem is that experimenters may need additional tools and documentation for interpretation of the results. To address this issue, we included graphical outputs and guiding comments that are accessible also for non-specialists throughout the demo scripts. Together with existing literature on the appropriate interpretation of raw calcium data^4,5, 22, 45, 58^, these subsidiary tools will help to focus the attention on data quality and make users aware of the potentials and limitations of raw and deconvolved data.

## DISCUSSION

We have created new tools and resources for spike inference from calcium imaging data. Any spike inference approach, in particular methods based on deep learning, critically depend on the availability and quality of ground truth datasets. We therefore created a ground truth database that is larger and more diverse than previous datasets^15, 18^, spanning multiple calcium indicators, brain regions and species (Fig. 1). Moreover, we developed CASCADE, a novel algorithm for spike inference based on deep learning. The central idea of CASCADE is to optimize the match between the training data and experimental datasets, rather than to invest primarily into the optimization of the inference algorithm itself. Previous supervised spike inference algorithms typically trained models with an existing, immutable ground truth^15, 18, 19^. Our algorithm, in contrast, resamples the ground truth datasets upon demand to match both frame rate and noise level in an automated fashion for each neuron (Fig. 2). Training with an appropriately resampled and noise-matched ground truth resulted in improved inference, highlighting the importance of training the algorithm not only with realistic calcium signals but also with realistic noise patterns.

The generalization of spike inference methods across unseen datasets had been investigated sporadically^12, 13, 21^ but never systematically in previous studies, presumably due to the lack of extensive ground truth data. Taking advantage of our diverse ground truth database, we explored how predictions depend on species (zebrafish or mouse), indicator type and brain region (Fig. 3) as well as on other experimental parameters that are presumed to strongly influence spike inference. Surprisingly, we found that some training datasets allowed for efficient generalization across these parameters, and a combined training dataset achieved very high performance across all test data. This result was obtained for both excitatory neurons and interneurons, although absolute spike rates were underestimated for interneuron data. Moreover, some datasets performed poorly as training sets while others performed poorly as test sets, even when compared against datasets with a similar indicator and/or from the same brain region. These observations suggest that generalization is affected significantly by experimental differences that are difficult to identify, such as indicator concentration or baseline calcium concentrations. However, this problem could be overcome by training networks on a diverse ground truth database, indicating that networks can learn to take these variations into account when sufficient information is provided during training. Highly efficient generalization across excitatory datasets was obtained by the unsupervised ‘global EXC model’ that was trained on all available excitatory ground truth datasets. This global model is therefore well-suited for practical applications of spike inference in unseen datasets.

In all investigated situations, our algorithm outperformed existing approaches (Fig. 4). Predictions were not only more precise, as measured by correlation metrics, but also less biased towards underestimates of true spike rates. We reason that the balance between spike detection and noise suppression is crucial for reliable spike inference. Our results suggest that less expressive models have to over-suppress if specific suppression is not possible. In contrast, the expressiveness of the employed deep networks enables CASCADE to better distinguish signal from noise, while their relatively small size prevents overfitting. It is however possible that other algorithms might perform better in regimes that are not covered by the current ground truth database (*e.g.,* recordings with extremely low noise levels that can easily resolve single spikes, or tonically spiking neurons that are transiently inhibited^59^).

CASCADE was not sensitive to user-adjustable hyper-parameters or the class of the deep networks tested. This insensitivity has two consequences. First, it seems more valuable to us to optimize the acquisition of more specific and diverse ground truth and the preprocessing of calcium data rather than to focus on improvements of the deep networks. Second, because hyper-parameters do not need to be adjusted by the user, the application of spike inference becomes simple in practice. While some previous studies assumed that user-adjustable parameters in model-based algorithms increase the interpretability of the model^10–13^, we argue here that (1) biophysical model parameters are often ambiguous^12^ and therefore not directly interpretable, and (2) it is more important to focus on the interpretability of the results rather than the model. To this end, our toolbox provides methods to estimate the errors made during spike inference. Moreover, we included a detailed documentation in the Colaboratory Notebook to help the user interpret the results.

Quantitative inference of spike rates is critical for the analysis of existing and future calcium imaging datasets^4, 5, 22^. The approach usually requires single-neuron resolution and is less well suited when the ΔF/F values are less well defined (*e.g.,* in endoscopic one-photon data with high background fluorescence, fiber photometry or wide-field calcium imaging). Moreover, ΔF/F can, in theory, only report spike rate changes. Nevertheless, we found that absolute spike rates can be reliably inferred when the baseline activity is sufficiently sparse to enable the determination of the fluorescence baseline level F_0_, which was the case in all datasets examined here (Fig. 5,6). The enhanced temporal resolution will be particularly useful for the analysis of neuronal activity during natural stimulus sequences and behaviors that occur on timescales shorter than typical durations of calcium transients. For example, the deconvolution of calcium signals can increase the effective temporal resolution to timescales below 100 ms, which will allow for the analysis of neuronal representations across theta cycles^60^ and for the resolution of early and late dynamics in cortical responses to sensory inputs that have been associated with different processing steps^61^. Moreover, the inference of absolute spike rates will help improve the calibration of precisely patterned optogenetic manipulations^62–64^ and the extraction of constraints, *e.g.,* absolute spike rates, for computational models of neural circuits.

The reliability of spike inference obviously depends on the recording quality of the calcium imaging data. To improve data quality of ΔF/F signals, future work should focus on the reduction of movement artifacts and neuropil contamination both by experimental design^45, 65^ and by extraction methods^40–43^, including the correct estimation of the F0 baseline despite unknown background fluorescence. In the long term, the development of more linear calcium indicators^66^ and especially the acquisition and integration of more specific ground truth, *e.g.*, for additional interneurons and subcortical brain regions, will enable quantitative spike inference for an even broader set of experimental conditions. We envision that our set of ground truth recordings will become enlarged over time, allowing to train more and more specific models for reliable inference of spike rates.

## ACKNOWLEDGEMENTS

We thank the members of the GENIE project, the Allen Institute and the Spikefinder project for publicly providing existing ground truth datasets together with excellent documentation. We thank Philipp Berens and Emmanouil Froudarakis for providing additional information on the Spikefinder datasets. We thank Gwendolin Schoenfeld for helpful discussions on dataset 18, and Hendrik Heiser, Nesibe Temiz, Chie Satou, Gwendolin Schoenfeld and Henry Luetcke for testing earlier versions of the toolbox. This work was supported by grants to F.H. from the Swiss National Science Foundation (Project grant 310030-127091; Sinergia grant CRSII5-18O316) and by the European Research Council (ERC Advanced Grant BRAINCOMPATH, grant agreement no. 670757), by grants to K.K. from MEXT, Japan (Scientific Research for Innovative Areas, no. 17H06313), by grants to R.W.F. from the Swiss National Science Foundation (Project grant 310030B-152833/1) and from the European Research Council (ERC Advanced Grant MCircuits, grant agreement no. 742576), by the Novartis Research Foundation, by a UZH Forschungskredit and a fellowship from the Boehringer Ingelheim Fonds to P.R..

## CONTRIBUTIONS

P.R. conceived the project, developed the algorithm, performed ground truth recordings (datasets 4-8), performed all analyses, developed the toolbox and wrote the paper. S.C. performed ground truth recordings (datasets 18 and 19). A.H. developed the toolbox. M.E. and K.K. (dataset 3), A.K. and Y.D. (datasets 2, 22 and 23), A.B. and S.H. (datasets 24-27) performed and preprocessed ground truth recordings. F.H. supervised ground truth recordings (datasets 18 and 19) and the development of the toolbox, and wrote the paper. R.W.F. supervised ground truth recordings (datasets 4-8) and the development of the algorithm, and wrote the paper.

## METHODS

### Simultaneous juxtacellular recordings and calcium imaging in adult zebrafish

All zebrafish experiments were approved by the Veterinary Department of the Canton Basel-Stadt (Switzerland). For the recordings in DS#04 and DS#05, the adult zebrafish brain was dissected *ex vivo* as described previously^23^ and OBG-1 AM or Cal-520 AM were injected and incubated in posterior Dp (pDp) as described previously^67^. During the dissection, the dura mater above pDp was carefully removed to prevent clogging of the patch pipette. Calcium indicators were injected for 1-2 min at two locations (injection 1: ∼210 μm dorsal from the ventralmost aspect of Dp and ∼130 μm from the lateral surface of Dp; injection 2: 180 μm and 60 μm) and was monitored by snapshot multiphoton images. The pressure was adjusted to avoid fast swelling of the tissue.

Juxtacellular recordings were performed >1h and <4h after the dye injection. Patch pipettes were pulled from 1 mm borosilicate glass capillaries (Hilgenrath), with a pipette resistance of 5-8 MΩ. Micropipettes were backfilled with ACSF (in mM: 124 NaCl, 2 KCl, 1.25 KH_2_PO_4_, 1.6 MgSO_4_, 22 D-(+)-Glucose, 2 CaCl_2_, 24 NaHCO_3_; pH 7.2; 300-310 mOsm) with 0.05 mM Alexa 594.

The explant preparation was rotated about the anterior-posterior axis to allow for optical access from the side (sagittal imaging). Using a multiphoton microscope, images generated from fluorescence and from the asymmetry of the signal on a four-quadrant detector for transmitted light were used to target the pipette to pDp, while continuous low pressure (30-40 mbar) was applied to prevent clogging of the pipette. The pipette then entered the tissue with initial high pressure (90-110 mbar) that was lowered after a few seconds. Neurons were approached using the shadow-patching technique described previously^51, 68^, but with lower pressure. Juxtacellular recordings were performed after establishing a loose seal (typically 30-50 MΩ) with a target neuron. In some cases, a small negative pressure was applied initially to improve the electrical contact with the target cell. In several cases, single micropipettes were used multiple times. Recordings were performed in voltage-clamp mode with the voltage adjusted such that the resulting current approximated zero^69^.

For DS#06-08, which were based on a transgenic line expressing GCaMP6f in the forebrain^49^, the experimental procedures were similar except for the injection of synthetic dyes. Because the baseline brightness of GCaMP6f is low it was often difficult to identify individual neurons. Upon application of odor stimuli, stimulus-responsive neurons that expressed GCaMP6f became brighter, which permitted reliable visual identification for targeted patching. For regions in dD (DS#07) with no obvious odor responses, cells were patched randomly based on shadow images generated by the blown-out Alexa dye^68^.

Simultaneous recordings of fluorescence and extracellular spikes of the same neuron were synchronized using *Scanimage 3.8* for imaging^70^ and *Ephus* for electrophysiology^71^. Calcium imaging was performed at intermediate zoom (see Fig. 1) with a frame rate of 7.5 or 7.8125 Hz for DS#04 and DS#05 and at high zoom with a framerate of 30 Hz for DS#06-08. Electrophysiological recordings were low pass-filtered at 4 kHz (4-pole Bessel filter) and sampled at 10 kHz.

Recordings were performed in 120-s episodes, with repeated stimulation of the nose with food extract odorants as described previously^51^. In pDp, spike rates are usually very low. When no spiking activity was observed, the holding potential of the pipette was set to higher values (between +5 and +30 mV), leading to an extracellular current that depolarized the neuron if the seal resistance was sufficiently high, resulting in artificially generated spikes. If no spikes could be elicited over the full duration of the recording, the recording was not included in the ground truth dataset.

### Anatomical location and further information about neurons in zebrafish ground truth datasets

*DS#04: OGB-1, injected in the posterior part of the olfactory cortex homolog (pDp) in adult zebrafish*. Recordings were performed throughout dorsal and ventral compartments of pDp and OGB-1 was injected as described previously^67^. Because OGB-1 localizes predominantly to the nucleus and because the resolution was high, neuropil contamination is negligible in this dataset.

*DS#05: Cal-520, injected in the posterior part of the olfactory cortex homolog (pDp) in adult zebrafish.* Same brain region as DS#03. Unlike OGB-1, Cal-520 is primarily cytoplasmic, resulting in considerable neuropil contamination. Cal-520 spread less than OGB-1 after injection and labeled only a small central volume in pDp.

*DS#06: tg(NeuroD:GCaMP6f), anterior part of the olfactory cortex homolog (aDp) in adult zebrafish.* In this transgenic line, GCaMP6f is strongly expressed throughout Dp. Recording location and framerate were chosen to match previous experiments^51^.

*DS#07: tg(NeuroD:GCaMP6f), dorsal part of the dorsal pallium (dD) in adult zebrafish.* All recorded neurons in dD were mapped onto brain regions Dm, Dl, rDc and cDc based on neuroD expression in the dorsal part of the dorsal pallium (Fig. S22, following Huang et al., 2020, ref. ^72^). Although the dorsal pallium is not known to be directly involved in olfactory processing, we noticed that several neurons were clearly inhibited during odor stimulation (duration, 10 - 30 s).

*DS#08: tg(NeuroD), olfactory bulb (OB) in adult zebrafish.* In the olfactory bulb of this transgenic line, GCaMP6f is restricted to a distinct, small subset of putative mitral cells and interneurons^49^. Neurons #1-#3, #5 and #7 were identified as interneurons based on their small size and morphology, while neurons #4, #6, #8 and #9 were classified as putative mitral cells.

### Simultaneous juxtacellular recordings and calcium imaging in anesthetized mice

For virus-induced expression of R-CaMP1.07 (DS#19), AAV1-EFα1-R-CaMP1.07 and AAV1-EFα1-DIO-R- CaMP1.07 were stereotactically injected under isoflurane anaesthesia in barrel cortex of C57BL/6J mice and hippocampal area CA3 of tg(Grik4-cre)G32-4Stl mice as described previously^25^. We performed combined electrophysiology and in vivo calcium imaging in acute experiments in anesthetized animals (n = 3; at least two weeks after virus injection) as described previously for barrel cortex recordings^25^. A stainless steel plate was fixed to the exposed skull using dental acrylic cement. A 1x1 mm^2^ craniotomy was made over barrel cortex. The dura mater was cleaned with Ringer’s solution (containing in mM: 135 NaCl, 5.4 KCl, 1.8 CaCl2, 5 HEPES, pH 7.2 with NaOH) and carefully removed. To reduce tissue motion caused by heart beat and breathing, the craniotomy was filled with low concentration agarose gel and gently pressed with a glass coverslip. For CA3 recordings (dataset #18), a 4-mm Ø craniotomy was centred over the injection site. The overlying cortex was aspirated until the corpus callosum became visible. The cavity was filled with 1% agarose gel to reduce tissue motion. Juxtacellular recordings from R-CaMP1.07-expressing neurons were obtained with glass pipettes (4–6 MΩ tip resistance) containing Ringer’s solution. For pipette visualization, Alexa-488 (Invitrogen) was added to the solution or pipettes were coated with BSA Alexa-594 (Invitrogen). Action potentials were recorded in current clamp at 10 kHz sampling rate using an Axoclamp 2B amplifier (Axon Instruments, Molecular Devices) and digitized using Clampex 10.2 software. All experimental procedures were conducted in accordance with the ethical principles and guidelines for animal experiments of the Veterinary Office of Switzerland and were approved by the Cantonal Veterinary Office in Zurich.

For dataset #03, C57BL6/J male mice were anesthetized by intraperitoneal injection of 1.9 mg/g urethane and the skull was partly exposed and attached to a stainless steel frame as described previously^26^. In a small craniotomy of the barrel cortex, we removed the dura, filled the cranial window with 1.5% agarose and placed a coverslip over the agarose to minimize brain movements^26^. Cal-520 AM together with an Alexa dye were bolus-loaded in layer 2/3 of the barrel cortex (200–300 μm deep below the surface) and monitored by two-photon imaging on the Alexa channel^26^. Calcium imaging was performed more than 30 min after dye ejection. For simultaneous calcium imaging and loose-seal cell-attached recordings, we filled glass pipettes (5–7 MΩ) with the extracellular solution containing Alexa 594 (50 μM), inserted pipettes into the barrel and targeted Cal-520-loaded somata. At about 10 min after the establishment of the cell-attached configuration, we performed simultaneous loose-seal cell-attached recording and high-speed line-scan calcium imaging (500 Hz) on the soma of cortical neurons as described^26^. The electrophysiological data were filtered at 10 kHz and digitized at 20 kHz by using Multiclamp 700B and Digidata 1322A (Molecular Devices), and acquired using AxoGraph X (AxoGraph).

Datasets #24-#26 were recorded in slices of mouse visual cortex as described previously^27^. Interneurons were targeted by injecting GCaMP6f-expressing AAV1 virus into PV-Cre, VIP-Cre or SOM-Cre mice. Coronal slices were cut with a thickness of 350 µm and loose patch recordings performed at 32°C in ACSF. To induce activity in otherwise quiet slices, a potassium-based solution was applied to the slice through a second pipette. Simultaneous calcium imaging was performed with a two-photon microscope recording at 34 Hz through a 16x water immersion objective (0.8 NA, Nikon) as described^27^. Dataset #27 was recorded in anaesthetized mice as described previously^73^. Adult (> 8 weeks) PV-tdTomato mice (cross between Rosa-CAG- LSL-tdTomato (JAX: 007914) and PV-Cre (JAX: 008069)) were injected with GCaMP6f-AAV (AAV1.Syn.GCaMP6f.WPRE.SV40, UPENN) in primary visual cortex (V1, ∼2.5 mm lateral, ∼0.7 mm anterior of the posterior suture). Acute recordings were performed at least 2 weeks after the initial injection. Mice were initially anaesthetized with a mixture of fentanyl (0.05 mg/ml), midazolam (5.0 mg/kg), and medetomidin (0.5 mg/kg), a metal headplate was fixed on the skull and a craniotomy was opened above V1. Anesthesia was maintained with a low concentration of isoflurane (0.5% in O2). Borosilicate glass pipettes (6–8 MΩ) filled with a solution containing 110 mM potassium gluconate, 4 mM NaCl, 40 mM HEPES, 2 mM ATP-Mg, 0.3 mM GTP-NaCl, and 0.03 mM Alexa 594 (adjusted to pH 7.2 with KOH, ∼290 mOSM) were lowered in the visual cortex. Neurons expressing GCaMP6f and tdTomato were targeted for juxtacellular recording in loose-cell configuration under a two-photon microscope. For simultaneous electrophysiological and optical recordings, calcium was recorded with Scanimage^70^ at 30 Hz, and juxtacellular voltage was recorded using a Multiclamp 700B amplifier (Axon Instruments, USA), filtered at 20 kHz and digitized at 10 kHz (National Instruments, USA). 50 Hz noise was reduced by using a noise eliminator (Humbug).

Datasets #02, #22 and #23 were recorded in mouse primary visual cortex as described previously^28^. GFP- GIN mice were used to target SOM interneurons, PV-Cre mice crossed with loxP-flanked tdTomato reporter mice were used to target PV interneurons, and CaMKIIα-Cre mice crossed with loxP-flanked tdTomato mice were used to target excitatory neurons. 1h after loading of OGB-1 into V1^28^, two-photon microscopy was used to target neurons 150-300 µm below the brain surface with the recording pipette, while the mouse was anesthetized with intraperitoneal injection of urethane and chlorprothixene. Two-photon imaging of neurons was performed with a 40x objective at a frame rate of 15.6 Hz while voltage was recorded in a loose-cell configuration from the same neuron as described^28^.

### Analysis of simultaneous juxtacellular recordings and calcium imaging

Movies of calcium indicator fluorescence images were corrected off-line for movement artifacts, i.e., slow drifts due to relaxation of the brain tissue for zebrafish data or fast movement artifacts for recordings in anesthetized mice. Ground truth recordings from DS#03 were not corrected for movement artifacts due to the scanning modality (line-scan). Afterwards, regions of interest (ROIs) were manually drawn using a custom-written software tool (https://git.io/vAeKZ)51 for each trial to select pixels that reflected the calcium activity of the neuron. Fluorescence traces were extracted either as average across the ROI or individually for each pixel to allow for both natural and artificial sub-sampling of calcium signal noise levels (Fig. S4).

Spike times were extracted from juxtacellular recordings using a custom-written template-matching algorithm. In brief, peaks of the first derivative of a 1 kHz-filtered electrophysiological signal were detected using a threshold that differed between recordings and that was manually adjusted to safely exclude false positives. The original waveforms of the detected events were then averaged and used in a second step as a template to detect all events across the full recording more precisely via cross-correlation of the template with the original signal. A threshold adjusted manually for each recorded neuron extracted action potential events. The process of first generating a template that was afterwards used to detect stereotypic signals allowed to increase the signal-to-noise of detected events, similar to previous usages of template matching in electrophysiology^74, 75^.

### Quality control

All electrical spiking events were inspected visually and compared to simultaneously recorded calcium transients. Any recordings that were ambiguous due to low electrophysiological signal-to-noise of action potentials were discarded and not used for any further analysis. Calcium recordings with excessive movement artifacts or apparent inconsistencies of juxtacellular and calcium recordings were discarded entirely. Excessive movement artifacts were defined as events when the neuron visibly moved out of the imaging plane, such that transients generated by these movements were almost as frequent and prominent as true calcium transients. Apparent inconsistencies of recordings were identified as recordings where no spike events corresponded to visible calcium transients and where a spike-triggered average (Fig. S1) did not show any signal, indicating that juxtacellular and calcium recordings were performed from different neurons. In addition, neurons were discarded when they did not spike at all even after application of currents, or when they became visibly brighter after establishing a loose seal due to unknown, possibly mechanical reasons. When the calcium recording clearly contained events without corresponding electrophysiological action potentials, the calcium trace of the manually drawn ROI and the calcium traces of adjacent neurons or neuropil were inspected together with the electrophysiological recordings in order to assess optical bleed-through, and ROIs were adjusted if necessary to avoid contamination. Occasionally, we also noted that mechanical stress exerted by the recording pipette can increase the brightness of the recorded neuron^30^, possibly by the release of calcium from internal stores. Recordings made during and after such events were discarded. Bursting can lead to adaptation of the extracellularly measured spike amplitude. Such recordings (e.g., in DS#18 with bursts of >10 APs with an inter-spike interval of ca. 5 ms) were carefully inspected for missed low-amplitude action potentials, in particular during these bursts.

### Extraction of ground truth from publicly available ground truth datasets

Additional ground truth was extracted from publicly available datasets and quality-controlled for each _neuron_^15,18,29,31–33^.

#### The Allen Institute datasets

For DS#10-13 from Huang et al. (2020)^29^, raw fluorescence traces were extracted from the processed datasets which were downloaded from https://portal.brain-map.org/explore/circuits/oephys. Neuropil signal was subtracted using the same standard scaling value for all neurons to make recordings comparable with other datasets (neuropil contamination ratio 0.7), despite the caveats associated with this procedure^29^. A 6- s running 10% lowest percentile window was typically used to compute F0 for ΔF/F0 calculation, but percentile values were adjusted to the noisiness of the recording and over window durations that were adjusted to the baseline activity. Simultaneous juxtacellular and calcium imaging recordings were inspected for each ground truth neuron together with the raw movie, described in the Methods section ‘Quality control’.

#### The Spikefinder datasets

For DS#01, DS#15 and DS#16 from Theis et al. (2016)^18^, the ground truth recordings at their native sampling rates as released during the Spikefinder challenge^15^ were processed. This Spikefinder dataset consists of 5 separate datasets. Datasets 1 and 4 were excluded since fluorescence baseline and scaling were unknown. The other datasets were extracted as fluorescence traces, F0 was computed as the 10^th^ percentile value (adjusted depending on the spike rate of each neuron) and used to compute ΔF/F0. Some ground truth neurons were discarded due to a highly unstable calcium recording baseline, but no strict quality control was possible since the raw calcium imaging data were not available. As found during a previous study, some datasets of the Spikefinder challenge come with calcium recordings that are delayed with respect to the electrophysiological recordings^15^. We therefore manually corrected for delays of the calcium recording with respect to the electrophysiological recording based on visual alignment of extracted linear kernels. The same correction delay was applied across all neurons of a given dataset.

#### The GENIE datasets

Datasets DS#09, DS#014, DS#017, DS#20 and DS#21 were downloaded from http://crcns.org/data-sets/methods^31–33, 76,77^. For DS#09 and DS#14 ^33^, ROIs were extracted from raw calcium imaging data using the same approach as described above for R-CaMP1.07 data. Recordings with excessive movement artifacts or apparent inconsistencies of juxtacellular and calcium recordings were discarded entirely. Neuropil was subtracted using the same standard scaling value for all neurons (neuropil contamination ratio 0.7)^33^. F0 values were computed using percentile values that were adjusted to the noisiness of the recording, and over window durations that were adjusted to the baseline activity.

For datasets DS#17, DS#20 and DS#21, no raw calcium imaging data were available, therefore not allowing for strict quality control using raw calcium recordings as additional feedback. Neuropil signal was subtracted from raw fluorescence using the same standard scaling value for all neurons (neuropil contamination ratio 0.7)^31, 32^. F0 values were computed using percentile values that were adjusted to the noisiness of the recording, and over window durations that were adjusted to the baseline activity.

#### Population calcium imaging with OGB-1 in zebrafish pDp

*Ex vivo* surgeries, OGB-1 AM injections and calcium imaging were performed as described for juxtacellular recordings. Calcium imaging in Dp was performed using a custom-built multiplane multiphoton microscope based on a voice-coil motor for fast z-scanning as described^49^. Laser power below the objective was 29-35 mW (central wavelength 930 nm, temporal pulse width below the objective 180 fs), with higher laser power for deeper imaging planes.

Imaging was performed in 8 planes (256x512 pixels, ca. 100x200 µm) at 7.5 Hz over a z-range of approximately 100 µm. Due to slowly relaxing brain tissue, movement correction was applied every 5 min by acquiring local z-stacks with a z-range of ±6 µm. The maximum cross-correlation between a reference stack acquired before the experiment and the local z-stack indicated the optimal positioning which was targeted using the stage motors of the microscope.

For odor stimulation, amino acids (His, Ser, Ala, Trp; Sigma) were diluted to a final concentration of 10^-4^ M and bile acid (TDCA; Sigma) was diluted to 10^-5^ M in ACSF immediately before the experiment. Food extract was prepared as described^51^. Odors were applied for 10 s through a constant stream of ACSF using a computer-controlled peristaltic pump^51^ in pseudo-random order with three repetitions of each odor presentation.

### Extraction of linear kernels from ground truth data

To extract linear kernels, we used simple regularized deconvolution using the *deconvreg*(*Calcium*,*Spikes*) function in Matlab (Mathworks). This function computes the kernel which, when convolved with the observed *Spikes*, results in the best approximation of the *Calcium* trace. Linear kernels were similar on average when extracted using different deconvolution algorithms (Wiener deconvolution, Lucy-Richardson algorithm; data not shown).

To compute the variability of linear kernels across neurons within and across datasets (Fig. S1), we split the ground truth recording of each neuron in five separate parts and computed the linear kernels for each of the segments separately. If the coefficient of variation (standard deviation divided by mean) across these five values was lower than 0.5, the computation of the kernel amplitude was considered reliable and included in the plots in Fig. S1.

### Computation of noise levels

In the shot-noise limited case the mean fluorescence *F_0_* scales with *N*, the number of photons collected by the detector per second, and the fluorescence baseline fluctuations *σ^F^* scale with 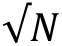. Thus, the *ΔF/F* baseline noise 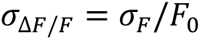 scales with 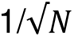. If the fluorescence signal is sampled at frame rate *f_r_*, the number of photons collected per frame reduces to *N/f_r_*, thus *σ*_Δ*F/F*_ scales with 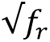. To define a noise measure that is independent of frame rate, we therefore normalized *σ*_Δ*F/F*_ for this shot-noise effect and defined the standardized noise ν as:

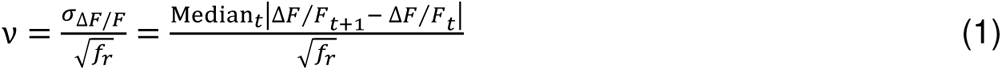

The units for ν are % · *Hz*^−1/2^, which for the purpose of readability we omit in the text. When computed for ΔF/F data in this way, ν is quantitatively comparable across datasets. A value of ν = 1 will always be a very low level, while ν = 8 will always be high, independent of frame rate.

### Metrics to quantify performance of spike inference algorithms: Correlation, Error and Bias

The ground truth spike rates were generated from detected discrete spikes by convolution with a Gaussian smoothing kernel (except in Fig. S21, where a non-Gaussian, causal kernel was applied). The precision of the ground truth was adjusted by tuning the standard deviation of the smoothing Gaussian (σ = 0.2 s for 7.5 Hz recordings and σ = 0.05 s for 30 Hz recordings). The ground truth spike rate was then compared to the inferred spike rate.

There is no single metric to reliably reflect the goodness of performance of a spike inference algorithm. Correlation between the inferred spike rate and the ground truth is widely used^15^ but does not contain information about different absolute scaling or offsets. *F_1_*-scores combine false positives and negatives^11^ but are difficult to compare across datasets when the baseline spike rates vary (which is the case for our database). Other metrics try to combine the strengths of the correlation measure with a sensitivity to the correct number of spikes^78^ but are less intuitive.

We defined three intuitive and complementary metrics (illustrated as color-coded equations in Fig. S5). First, we used the Pearson’s **correlation** between ground truth spike rate and inferred spike rates as a standard measure of the similarity. Second, the relative error (abbreviated as **error**) results from the sum of false positives and false negatives when subtracting the ground truth from the inferred spike rate, normalized by the absolute number of spikes in the ground truth. For example, an error of 0.7 would indicate that the number of either incorrectly inferred or omitted spikes is about 70% of the number of spikes in the ground truth. Third, the (relative) **bias** is defined as the difference of false positives and false negatives, again normalized by the absolute number of spikes in the ground truth. Algorithms that systematically underestimate spike rates will tend towards the minimum of the bias, -1, whereas other algorithms may tend to systematically overestimate spike occurrences (bias > 0). Importantly, the error can be very high when the number of false positives and false negatives is high, but the bias may still be zero. Error and bias are therefore two metrics that describe the absolute errors in terms of spike rates, thus complementing the correlation metric.

### Architecture of the default convolutional network

The default network consists of a standard convolutional network with in total 6 hidden layers, including 3 convolutional layers. The input consists of a window of 64 time points symmetric around the time point for which the inference is made. The three convolutional layers have relatively large but decreasing filter sizes (31, 19, 5 time points), with an increasing number of features (20, 30, 40 filters per layer). After the second and the third layer, maximum pooling layers are inserted. A final densely connected hidden layer consisting of 10 neurons relays the result to a single output neuron. While all neurons in hidden layers are based on rectified linear units (ReLUs), the output neuron is based on a linear identity transfer function. In total, the model consists of 18’541 trainable parameters.

The properties of the calcium imaging data are accounted for by resampling the ground truth with the appropriate noise levels and the matching frame rate. The ground truth is smoothed with a time-symmetric Gaussian kernel of standard deviation 0.2 s unless otherwise indicated for resampling at 7.5 Hz and 0.05 s for 30 Hz or a causal kernel (inverse Gaussian distribution) to facilitate gradient descent.

### Training deep networks for spike inference

To train the deep networks, the mean squared error between the smoothed ground truth spike rates and inferred spike rates was used as the loss function. This loss function not only optimizes the similarity of both signals (correlation), but also the absolute magnitude of the inferred spike rates. Based on errors computed via backpropagation, gradient descent was performed using a standard optimizer (*adagrad*; cf. Fig. S7). Based on a given resampled ground truth dataset, the network was trained using every single data point from this set, completing an epoch. Typically, training lasted for 10-20 epochs (except when analyzing overfitting in Fig. S7 and Fig. S8).

Crucially, in all spike inferences presented here, without exception, a leave-one-out strategy was employed. For example, to infer the spike rates of a given neuron in a dataset, the network was trained on all neurons of this dataset except the neuron of interest. To infer spike rates for a given set of datasets, the training set always excluded the dataset for which inferences were made. This strategy of cross-validation is absolutely crucial and is strictly distinct from the process of fitting parameters for a neuron or a dataset, which would yield better result for a given neuron but would fail to generalize to new data.

### Architecture of alternative deep learning networks

All deep learning architectures (Fig. S8) were trained with the same loss function, the same input and the same optimizer as the default network.

#### Small convolutional filters network

The same architecture as the default network, with the only difference that smaller convolutional filter sizes were used, (15, 9, 3) instead of (31, 19, 5). Total of 9’891 trainable parameters.

*Single convolutional layer network*

Consisting of the first convolutional layer of the default network, a single max pooling layer and a single dense layer of 10 neurons. Total of 1’021 trainable parameters.

#### Deeper convolutional network (5 CNN layers)

Consisting of 5 convolutional layers with filter sizes (11, 9, 7, 5, 3) and filter numbers (20, 30, 40, 40, 40), with three max pooling layers after the second, fourth and fifth convolutional layers, and a final dense layer expansion of 10 neurons. The reduction of the filter sizes compared with the default network is necessary since no zero-padding was applied, resulting in a decrease of the size of the 1D trace with increasing layer depth. Total of 27’421 trainable parameters.

#### Deeper convolutional network (7 CNN layers)

Consisting of 7 convolutional layers with filter sizes (7, 6, 5, 4, 3, 3, 3) and filter numbers (20, 30, 40, 40, 40, 40, 40), with three max pooling layers after the second, fifth and seventh convolutional layers, and a final dense layer expansion of 10 neurons. Total of 31’221 trainable parameters.

#### Batch normalization

Same as the default network, but with batch normalization^79^ for regularization after each convolutional and dense layer, but before the respective ReLU transfer functions of each network layer. Total of 18’741 trainable parameters.

#### Locally connected network

Same as the default network, but with locally connected filters instead of convolutional filters. For convolutional filters, filter weights are shared across each position in the image space (here, in the temporal window), while the filters are different for each position for locally connected networks. The rationale behind this architecture is that different filters can be learned for each position, which is intuitive given that spike detection is not invariant to the position of the calcium transient in the window. Using different weights for each position of the filter sets results in a total of 229’231 trainable parameters.

#### Naïve LSTM model

LSTM units are complex neuronal units with internal states and gates that are used in recurrent networks to overcome the problem of vanishing gradients when backpropagating through time^80, 81^. The time points of the input window are sequentially fed into the recurrent network, which are processed by the recurrent network, with earlier time points retained through recurrent activity or LSTM states and used to activate the network for processing of later time points. The investigated model consisted of two layers of each 25 LSTM units with ReLU as activation functions, followed by a simple dense expansion layer of 50 neurons with ReLU activation functions. Total of 4’051 trainable parameters.

#### Bi-directional LSTM model

The time points of the input window (64 data points) are split into past (32 data points) and future (32 data points) with respect to the time point used for spike inference (“presence”). Past and a reversed version of the future are each fed into a recurrent network based on a single layer of 25 LSTM units (with ReLU activations), such that the time point closest to “presence” is fed in last^82, 83^. The output of the two recurrent networks for past and future is concatenated and connected with a dense fully connected layer of 50 simple units (ReLU activations). Total of 8’001 trainable parameters.

#### Linear network

Same as the default network, but with linear activation functions instead of rectifying linear units (ReLUs). The network is therefore entirely linear, but is based on the same architecture in terms of connectivity. Total of 18’541 trainable parameters.

### Discretization of spiking probabilities

To obtain discrete spiking events from inferred probabilities, a brute-force fitting procedure was applied. The Gaussian kernel used to smooth the ground truth was used as a prior for the inferred spike rate that corresponds to a single action potential. The fit therefore consisted of optimally fitting a set of Gaussian kernels of the expected width and height to the inferred spike rate. We made a first guess that was then optimized by random modifications. The first guess was generated using Monte Carlo importance sampling, such that the overall number of discrete spikes matched the integral of inferred probabilities. Next, events were ranked in how they contributed to the fit by comparing the fit quality when single events were omitted. Lowest-ranking events were discarded and replaced by newly drawn events, again using importance sampling based on the residual probability distribution. Finally, each spike was shifted randomly over the entire duration and the best fit was used. This approach is relatively slow but results in a reliable fit. To speed up the procedure, spiking probabilities were divided in continuous sequences of non-zero support (divide-and-conquer strategy). For Fig. S12 and to allow for comparison against raw inferred spike rates, the resulting discrete spikes were convolved with the Gaussian smoothing kernel that had been used to generate the ground truth. We provide a demo script that infers discrete spikes from spike rates predicted with CASCADE (available on Github: https://git.io/JtZe4).

### Generalized linear model to fit predictability across datasets

To predict how well a model trained on a given ground truth dataset (*e.g.*, DS#08) is able to infer activity for another dataset (*e.g.*, DS#14), a set of descriptors (regressors) was extracted for each dataset, and a generalized linear model (GLM) was trained to predict this relationship based on the regressors of the two respective datasets (Fig. S14). In total, 8 predictors were used, separately or together.

First, *indicator species* was set to 1 if training and test dataset had the same indicator species (synthetic dyes vs. genetically encoded dyes) and 0 otherwise. *Animal species* was set to 1 if training and test dataset had the same animal species (zebrafish vs. mouse) and 0 otherwise. *Spike rate* was computed as the absolute difference between median spike rates across neurons from training and test datasets. *Burstiness* was computed as the number of spikes that were spike within 50 ms of the timing of a given spike. This metric quantifies the likelihood that a given spike is surrounded by other spikes. The *Fano factor* was computed by dividing the variance of inter-spike-intervals (ISIs) by the mean of ISIs^84^. Measured Fano factors were broadly distributed across datasets with a median of 3.7 and a standard deviation of 5.9, and an outlier dataset DS#18 in mouse CA3 with a Fano factor of 30.0. The *area of the linear kernel* was computed by summing up the area under the curve for the extracted linear kernel for each dataset. The *kernel decay constant* was computed without exponential fit by measuring the time between rise and decay time of the kernel directly. Rise and decay time points were identified by finding the first and last time point where the kernel surpassed 1/e of its maximum amplitude. The *correlation time course* was computed as the correlation of between the kernels of training and test dataset.

The GLM was fitted based on these regressors using the *glmfit()* command in Matlab with an identity linker function.

### Artificial ground truth generated with NAOMi

The package NAOMi was used to generate simulated two-photon calcium recordings of neurons with known spike patterns^45^. We used the default parameters, which had been optimized for the simulation of GCaMP6f based on previous calibrations^33^. Artificial ground truth was generated at 30 Hz with a detection NA of 0.6 and an excitation NA of 0.8 at a depth of 100 µm below the cortical surface, in a volume of 250x250x100 µm^3^. To increase the signal-to-noise ratio of the simulated recordings we used a relatively high simulated laser power of 70 mW. We simulated recordings of the central plane of five such volumes over a duration of 166 s. We extracted the cleanest components of each simulation by selecting the spatial components (from the *ideal components* returned by NAOMi) that correlated most highly with the known ground truth signals (correlation with a known somatic ground truth signal >0.80). We chose to only include the best-matching components since other components typically had much stronger neuropil contamination than our experimentally obtained ground truth recordings. Then, we extracted the fluorescence of the selected components and performed neuropil subtraction with a 2 pixel-ring around the detected component using a factor of 0.45 for neuropil subtraction. Afterwards, we computed the ΔF/F signal, using the 2^nd^ percentile across the entire recording to determine F0. This procedure resulted in ground truth recordings from a total of 250 simulated neurons.

### Adaptation of model-based spike inference algorithms

The MLSpike algorithm was downloaded from https://github.com/MLspike/spikes and used within Matlab 2017a ^11^. Parameter settings were manually explored for several datasets using the graphical demo user interface. Then, some parameters (noise level *sigma* and inverse frame rate *dt*) were fixed to the values constrained by the ground truth. The *drift* parameter was set to 0.1. For synthetic dyes (DS#01-05, DS#22- 23), a saturating non-linearity (*saturation* = 0.01) was used, whereas for all other datasets a GCaMP-like nonlinearity (*pnonlin* = [1.0 0.0]) was defined and kept the same across datasets, since predictions have been described to only slightly depend on the precise values of the non-linearity^11^. Based on manual exploration, the two parameters *tau* (decay time constant) and *amplitude* (amplitude of a single action potential) were explored in a grid search for all ground truth datasets and all noise levels separately. The grid search ranged from 0.1 to 5 s for *tau* and from 0.01 to 0.35 for *amplitude*.

The Peeling algorithm was downloaded from https://github.com/HelmchenLab/CalciumSim and used within Matlab 2017a ^10^. A single-exponential linear model with default values was used. A grid search was performed over two parameters for all ground truth datasets: time constant of the exponential decay (*tau1*) and the amplitude of a single spike (*amp1*). Grid search ranged from 0.25 – 5 s for *tau1* and from 2.5 – 35 for *amp1*. Discrete spike predictions were convolved with a Gaussian kernel such that the resulting trace optimized the loss function (mean squared error between predictions and ground truth).

The Python implementation of the L1-regularized OASIS algorithm in CaImAn was downloaded from https://github.com/j-friedrich/OASIS and used within Python 3.7 ^14^. The constrained version of OASIS was used to reduce the number of free parameters, with only one single free parameter, *g*. *g* relates to the exponential time fluorescence decay constant *τ* with the frame rate *f* via the formula: *g* = *e*^−1⁄*τf*^ . Grid search was performed for *g* in the range between 0.02 and 0.98, with a granularity of 0.02. The Python implementation of the FastL0SpikeInference algorithm from Jewell et al. (hence called Jewell&Witten) was downloaded from https://github.com/jewellsean/FastLZeroSpikeInference and used within Python 3.7 ^47^. A grid search was performed over two parameters for all ground truth datasets: Optimization was performed between 0.10 and 0.95 for the decay constant parameter *gamma*, and between 0.0001 and 0.75 for the L_0_ parameter *penalty*. Discrete spike predictions were convolved with a Gaussian kernel such that the resulting trace optimized the loss function (mean squared error between predictions and ground truth).

The Python implementation of the OASIS algorithm in Suite2p was downloaded from https://github.com/MouseLand/suite2p and used within Python 3.7 ^46^. Out of three tunable parameters (*tau*, *sig_baseline* and *win_baseline*), only the first two significantly affected the performance of the algorithm in our hands. *win_baseline* was set to 150 for all analyses. A grid search was performed over the two remaining parameters for all ground truth datasets: Optimization was performed between 0.5 and 3 for the decay time constant parameter *tau*, and between 2.5 and 20 for the parameter *sig_baseline*.

The optimal parameters resulting from the grid searches, which optimized the mean squared error between ground truth and inferred spike rates, are listed in Table 2 and provided as a csv file via Github (https://git.io/JtZe0). In addition to these parameters, we further used Gaussian smoothing kernels of variable standard deviation to find the amount of smoothing for each algorithm and dataset that optimized the mean squared error. Finally, to compensate for the propensity of several model-based algorithms to infer spike rates with a temporal lag compared to ground truth spike rates, we tested time shifts between - 1 and +1 s and used the respective shift that optimized the mean squared error for a given dataset to evaluate the algorithm in our analyses.

### Computational cost of spike inference

The six investigated algorithms exhibit different behaviors when scaling up the length of the calcium traces. For example, MLSpike and Peeling suffer from supra-linear cost when the duration of an analyzed calcium trace is increased, while CASCADE shows the opposite behavior due to its capability to parallelize spike inference. Therefore, all 26 full ground truth datasets, resampled at a noise level of 2 and a frame rate of 7.5 Hz, were used as a benchmark, consisting of recordings ranging from 10s of seconds up to several minutes. Processing time was averaged across all data points from all datasets. The time required to load the data from hard disk was not included. For CASCADE, the time for pre-processing the raw calcium data in order to generate a 64 point-wide segment for each time point was included in the benchmarking.

### Unsupervised sequence extraction using seqNMF

The Matlab-based toolbox seqNMF was used to extract temporal patterns for Fig. 5 in an unsupervised fashion^52^. Based on initial parameter exploration we used the following settings: K=7, L=20 and λ=0.002. K indicates the number of extracted patterns, L the number of time points for each pattern, λ serves as a regularizer to decorrelate the detected patterns^52^. The result of this unsupervised non-negative matrix factorization approach are K=7 temporal patterns that are each of them associated with a temporal loading which indicates when the temporal pattern became active. The temporal patterns and the temporal loadings provide low-complexity factors that break down the more complex population dynamics (Fig. 5).

### Allen Brain Observatory data

The complete calcium imaging data of the Allen Brain Observatory Visual Coding dataset were downloaded from http://observatory.brain-map.org/visualcoding via the AllenSDK with a Python interface. Layers were assigned based on imaging depth as described^53^. Imaging depth, transgenic lines, cortical areas and fluorescence traces were extracted from NWB files. For analysis, neuropil-corrected calcium traces from the Allen Brain Observatory dataset were used. Since all recordings were performed at an imaging rate of approximately 30 Hz, a single set of CASCADE models (,global EXC model‘ at 30 Hz, Fig. 3a) was used to predict spiking activity.

### Statistical tests and box plots

Statistical analysis was performed in Matlab 2017a and R. Only non-parametric tests were used. The Mann-Whitney rank sum test was used for non-paired samples (*e.g.*, comparison across datasets) and the Wilcoxon signed-rank test for paired samples (*e.g.*, comparison of predictions for the same set of neurons using two different algorithms). Two-sided tests were applied unless noted otherwise. Effect sizes Δ±CI (pseudo-median Δ and 95% confidence intervals CI unless otherwise indicated) were computed in R. Boxplots used standard settings in Matlab, with the central line at the median of the distribution, the box at the 25^th^ and 75^th^ percentile and the whiskers at the location of approximately 99.3 percent coverage for the case of a normal distribution.

## Supplementary information

⃞ Supplementary Note: Noise-matching of resampled ground truth data
⃞ Supplementary Note: Dependence of performance on hyper-parameters, overfitting and network architecture
⃞ Supplementary Note: Discrete spikes and single-spike precision
⃞ Supplementary Figures

## SUPPLEMENTARY NOTE: NOISE-MATCHING OF RESAMPLED GROUND TRUTH DATA

To ensure reliable inference of spike rates it is advantageous to train the supervised deep network with a training dataset that matches the noise level of the test neuron from the population imaging data. In practice, the noise level of each neuron from the population imaging data is determined and an existing model trained with approximately the same noise levels is loaded for spike inference. Noise-suppression is only effective if the noise statistics of the ground truth used for training the network resemble the noise statistics of the calcium data used for testing. To generate a ground truth that matches this requirement, we tested two different approaches to increase the noise of ground truth recordings to match population recordings (Fig. S4).

First, we used the raw imaging data to extract not only the mean fluorescence trace, but the fluorescence trace of each pixel of the region of interest (ROI) that defines the neuron. To achieve a given noise level ν, a random subset of pixels was drawn from the ROI pixels until the average fluorescence trace of this sub-ROI reached the desired noise level (Fig. S4a). This method generates realistic noise characteristics through spatial subsampling but is computationally costly and cannot be applied to ground truth datasets when raw fluorescence movies are not available.

Alternatively, we extracted only the mean fluorescence trace of the ROI of a ground truth neuron and added artificial noise until the overall high-frequency noise matched the test calcium dataset (Fig. S4a). This procedure can in theory be repeated to produce an infinite number of examples (replicas) from a simple ground truth recording. However, since the mean ΔF/F of a ground truth recording already is associated with a certain noise level, these noise patterns would be correlated across replicas. To avoid this undesired effect, which could lead to overfitting of correlated noise during training, the number of replicas was restricted to a number n that was computed with the noise level of the mean ΔF/F of a neuronal ROI, ν_ROI_, and the target noise level, ν_target_, by *n* = (ν_target_⁄ν_ROI_)^2^ and thresholded at a maximum of n = 500. We tested both simple Gaussian noise as well as Poisson noise, where the variance of the noise is proportional to the signal amplitude, as is typical for photon shot noise.

To test whether artificial noise enables the network to learn and suppress natural noise patterns we generated ground truth with either natural (spatial sub-sampling) or artificial (Gaussian or Poisson) noise for a subset of the available ground truth datasets (Fig. S4a-c). Using models trained on artificial rather than natural noise slightly but significantly decreased the correlation of predictions with the ground truth when applied to test datasets based on natural noise (decrease for Gaussian noise: Δ1 = 0.010±0.004, pseudo-median ± 95% C.I., p &lt; 1e-4, paired Wilcoxon test; decrease for Poisson noise: Δ1 = 0.006±0.004, p &lt; 0.005). This decrement indicates how much better a model trained with naturally sampled noise would likely perform. Conversely, testing models trained with artificially generated ground truth on artificially instead of naturally sampled ground truth increased the correlation of predictions slightly (Gaussian noise: Δ_2_ = 0.006±0.008, p = 0.13; Poisson noise: Δ2 = 0.005±0.009, p = 0.28). This increment indicates how much the correlations with the ground truth will be overestimated when training and testing is done with artificially generated ground truth only. Differences were generally more pronounced for larger noise levels, but overall remained very small compared to the absolute values (not shown). We therefore conclude that artificial noise allows deep networks to effectively learn noise patterns that can be applied to natural noise, with only minor performance loss compared to spatially subsampled recordings, and we further conclude that using Poisson-distributed noise yields slightly improved performance compared to artificial Gaussian noise.

## SUPPLEMENTARY NOTE: DEPENDENCE OF PERFORMANCE ON HYPER-PARAMETERS, OVERFITTING AND NETWORK ARCHITECTURE

We tested how spike inference performance depends on the choice of hyper-parameters and network architecture. Networks were trained on a specific ground truth dataset using all neurons except one, which was held out for testing (leave-one-out strategy). The algorithm turned out highly robust with respect to changes of the optimizer for gradient descent, the batch size during learning, the number of convolutional features per layer, the number of neurons in the dense layer, and the extent of the temporal window of the receptive field (Fig. S7a-e). All of these observations could be confirmed over a surprisingly large range, indicating that the network performance is highly robust with respect to any hyper-parameter choices.

In addition, we also investigated potential overfitting of the training dataset. We observed that the performance of the network was high already after one training epoch (i.e., as soon as every sample had been seen once by the network), then reached a maximum after 10-30 training epochs, and slightly decreased thereafter (Fig. S7f). This learning behavior suggests only moderate overfitting. At the same time the training loss decreased monotonically (Fig. S7g). We believe that the high abundance of noise and sparseness of events acts as a natural regularizer that prevents overfitting. More importantly, while the learning curve was smooth on average (Fig. S7f), individual network instances sometimes reached unfavorable states. As expected from known properties of deep networks85, this effect could be easily eliminated by ensemble averaging over 5 networks (Fig. S7h).

Finally, we also tested the performance when employing a network architecture different from the standard convolutional architecture (hence called ‘default’). We tested a large variety of standard deep learning architectures, including recurrent LSTM networks and non-convolutional deep networks with only the input, output and loss function remaining unchanged (see Methods for detailed descriptions and explanations). Most of these networks performed well, and the performance of several networks was statistically indistinguishable: the default convolutional network, a convolutional network with reduced filter size, a locally connected network, and a bi-directional LSTM network (Fig. S8a). Much smaller networks (single convolutional layer, Fig. S8a) tended to underfit the ground truth. On the other hand, networks with larger numbers of parameters (the deeper convolutional networks and the locally connected network) overfitted the data when training continued (dashed lines in Fig. S8b), consistent with previous observations^15^.

The locally connected network and the bi-directional LSTM network performed equally well compared to the default convolutional network despite very different architectures. However, some architectures that were not adapted to spike inference showed lower performance, for example the naïve LSTM network, which by its recurrent design prevents the network from looking precisely at the time point of interest (see Methods for details). Another example is a network identical to the default convolutional network but with purely linear activation functions (Fig. S8a,b), which prevents the algorithm from non-linearly adjusting decision boundaries.

## SUPPLEMENTARY NOTE: DISCRETE SPIKES AND SINGLE-SPIKE PRECISION

Many existing spike inference methods do not aim at the inference of spike rates, but rather of discrete spikes^10, 11, 14^. Previous publications reported that the average ΔF/F value triggered by a single spike is larger than zero, and that the calcium transients corresponding to single spikes can be detected in selected neurons^33^. However, the precise identification of individual spikes in practical situations is more challenging since a detection scheme should also work in unseen data. In addition, the task may depend in unknown ways on the variable expression of calcium indicators, on shot noise, on other noise sources, on low sampling rates and on the non-linear response of calcium indicators^3, 21, 29, 30, 86^. It is therefore not clear whether discrete spikes can be reliably inferred in realistic scenarios. We therefore devised two approaches to test whether single-spike precision can be achieved. First, we focused on single, isolated spikes in the ground truth and compared them with inferred spike rates for the same time window. This approach is discussed in the main text. Second, we transformed spike rates into discrete spikes and analyzed whether the discretization improved spike inference.

With respect to the second approach, we argue that the spike rates inferred by the deep network will exhibit a tendency to quantize if the predictions are close to single-spike precision. Therefore, a procedure that takes into account the prior about discretized spiking could improve the inferred spike rates. We therefore applied an algorithmic procedure that uses prior knowledge about the spike rate (spiking probability) waveform associated with a single spike to fill up the inferred probability trace with discrete spikes using an optimization procedure based on Monte-Carlo importance sampling (Methods).

We found that spiking probabilities that were almost correctly inferred by the deep network were optimized by suppression of noise or by rounding of close matches (blue arrows in Fig. S12a-d). However, this procedure can also enhance small false positive or small negative errors (red arrows in Fig. S12a-d). Over all ground truth datasets, the correlation metric slightly but consistently decreased when spike rates were discretized (Fig. S12e), indicating that the data quality did not allow for efficient use of the prior. Although discretized spike rates tended to decrease the error (Fig. S12e), this effect was primarily due to suppression of small noise events in the absence of spiking, and we found that this positive effect could be achieved without reduction of the correlation by thresholding the inferred spike rates (Fig. S12d,f). Together, this suggests that the available datasets do not permit discretization of predicted spike rates without performance loss.

Despite these caveats, discretization of spike rates might still be useful for two reasons. First, discrete spike events may be more intuitive visualizations of activity than smooth probabilities. Second, while spike rates are smoothed with a Gaussian kernel for each spike, the detection of a single spike that optimally explains this spike provides better temporal resolution. Here, we see a potentially useful application of discrete spike inference, which is, however, beyond the scope of this study. We include the algorithm to discretize spike rates as a script in the public repository (https://git.io/JtZe4).

## SUPPLEMENTARY FIGURES

**Figure S1.**
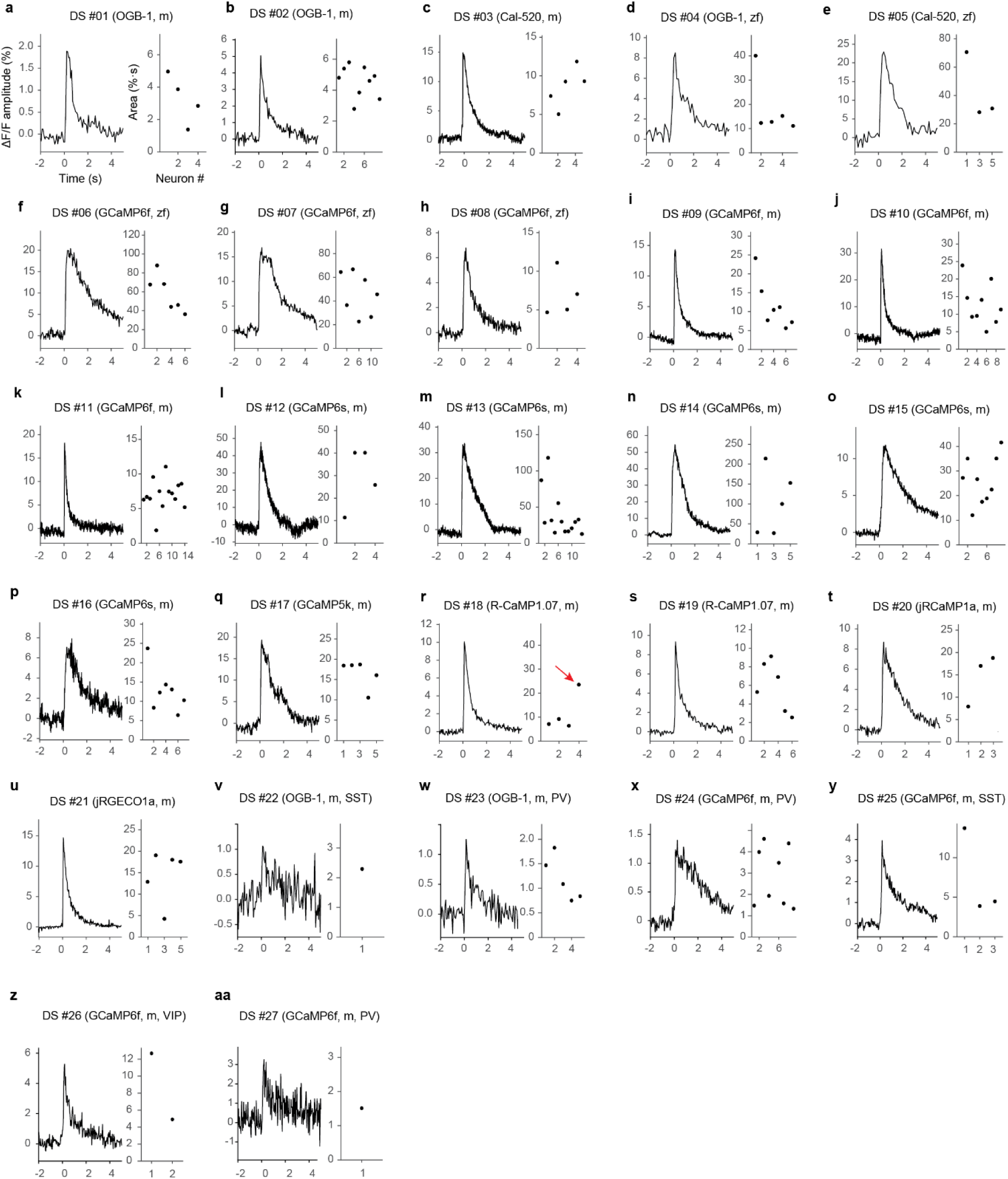
Linear kernels extracted from all ground truth datasets. The kernels are optimized such that when the ground truth spike times are linearly convolved with the kernel, the experimentally recorded ΔF/F trace is ideally approximated. In practice, this is achieved using regularized linear deconvolution of calcium traces based on spike times (Methods). Kernels vary both in amplitude and shape across datasets and within datasets. For single neurons, the kernel area (right panels) is only shown if the kernel could be reliably determined, as tested with the variability of the kernel across the recording (Methods). The red arrow in panel (r) indicates an outlier case that is discussed in Fig. S9a. m: Mouse, zf: Zebrafish.

**Figure S2.**
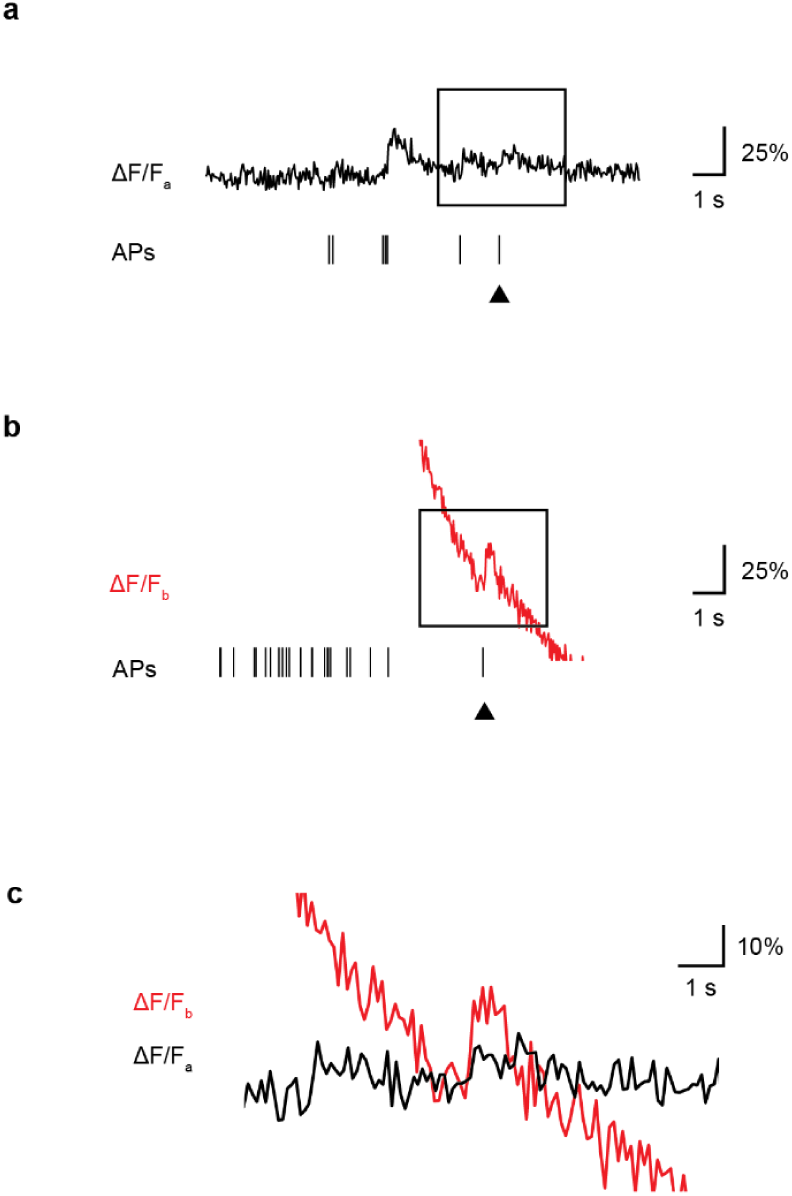
Example of history-dependence of spike-evoked fluorescence. Calcium recordings of the same neuron during a phase of sparse spiking **(a)** and after a burst **(b)** are shown. Single spikes are marked with a black arrowhead. The corresponding ΔF/F changes for cases a and b are overlayed and magnified in **(c)**, clearly showing a much larger fluorescence increase evoked by a single spike after the burst. This amplification is due to the non-linear cooperative calcium binding of GCaMP6f, with a larger fraction of indicator molecules being in a pre-bound state briefly after the burst. Example taken from DS#06.

**Figure S3.**
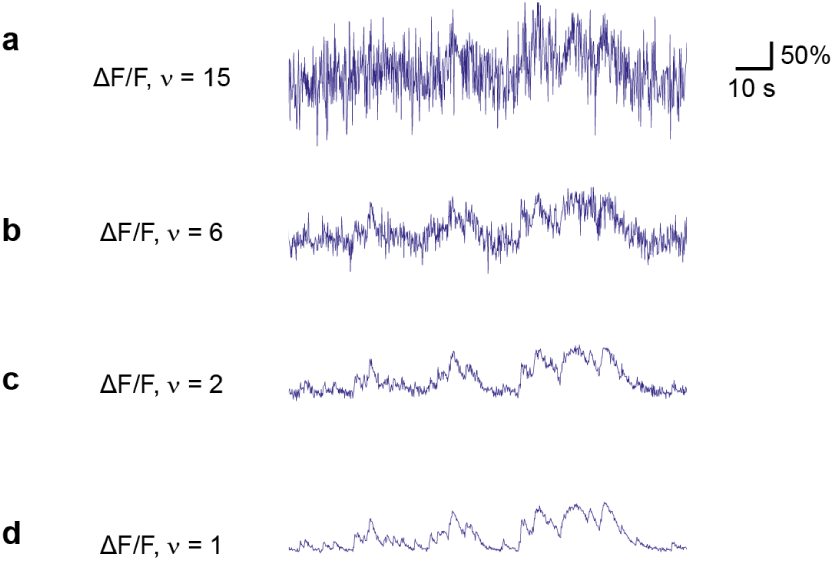
Illustration of different baseline noise levels. ΔF/F ground truth traces were resampled with added noise to reach the target noise level ν. **a-d**, Noise level illustration from ν = 15 (very high noise level) to ν = 1 (very low noise level). Standardized noise ν is given in units of % ·*Hz*^−1/2^.

**Figure S4.**
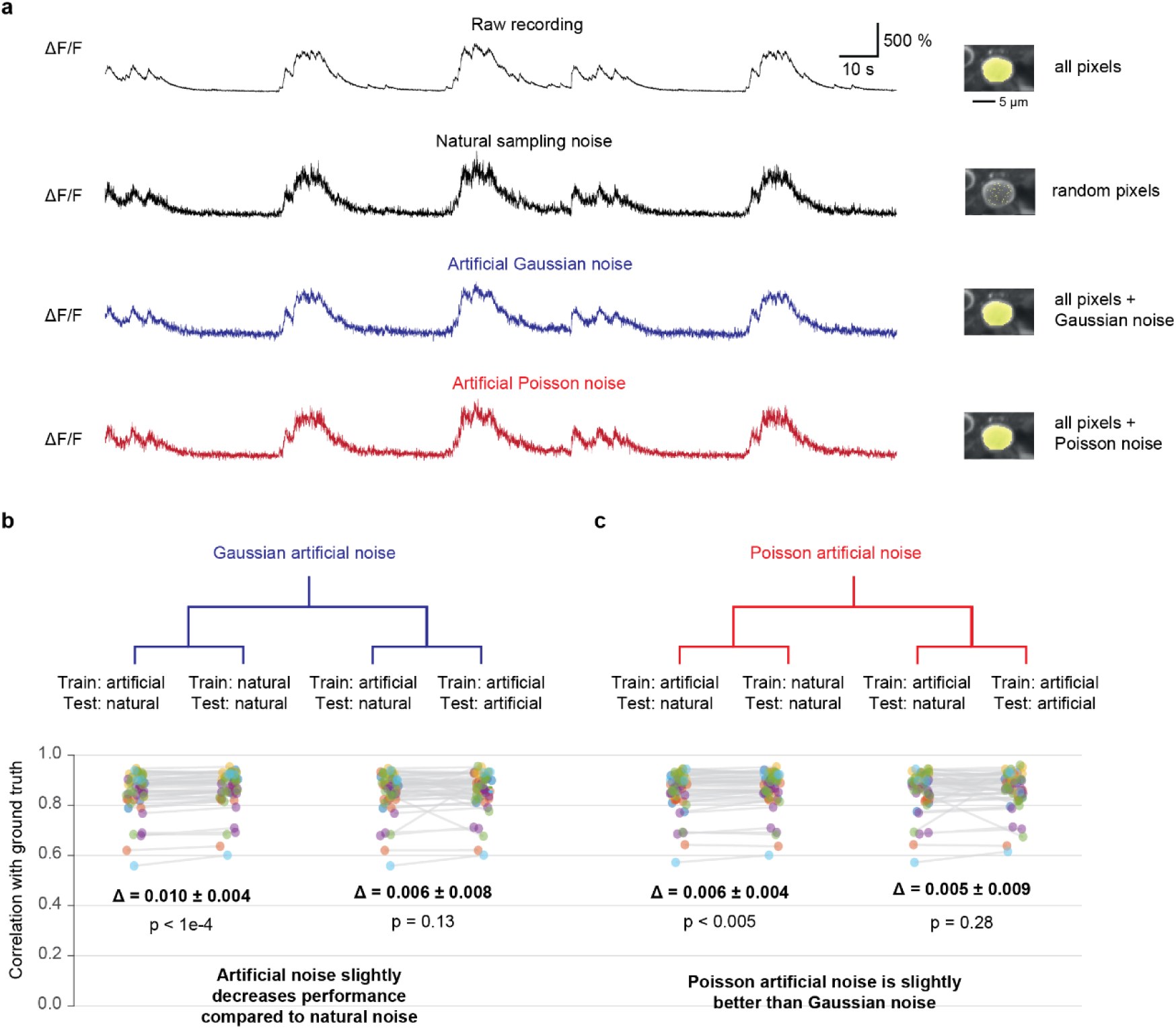
Natural resampling of the ground truth vs. artificially sampled noise. **a,** Row 1: Raw ground truth calcium recording, mean fluorescence. Row 2: Same ground truth recording, spatial subsampling (natural sampling) of random pixels in the ROI. Row 3: Same recording, mean fluorescence with added Gaussian noise. Row 4: Same recording, mean fluorescence with added Poisson noise. **b,** Testing different noise modes. For each condition, two comparisons between models are made: 1) Trained with artificial and tested with natural noise vs. trained and tested with natural noise (example with black arrow). This comparison shows how much worse the predictions become when applying the algorithm to naturally sampled calcium recordings while training the algorithm with artificially sampled noise. 2) Trained with artificial and tested with natural noise vs. trained and tested with artificial noise (example with red arrow). This comparison shows how much the procedure of training and testing with artificial noise overestimates the performance compared to the realistic case of training with artificial noise and applying the model to naturally sampled calcium recordings. Each data point represents a neuron, colors indicate ground truth datasets. Differences Δ were computed as pseudo- median ± 95% confidence intervals. P-values and pseudo-medians were computed using paired Wilcoxon signed-rank tests. All analyses were performed with a standardized noise level of ν = 2. Effect sizes increase slightly when increasing the noise level but remain around Δ ∼ 0.01 also for higher noise (data not shown).

**Figure S5.**
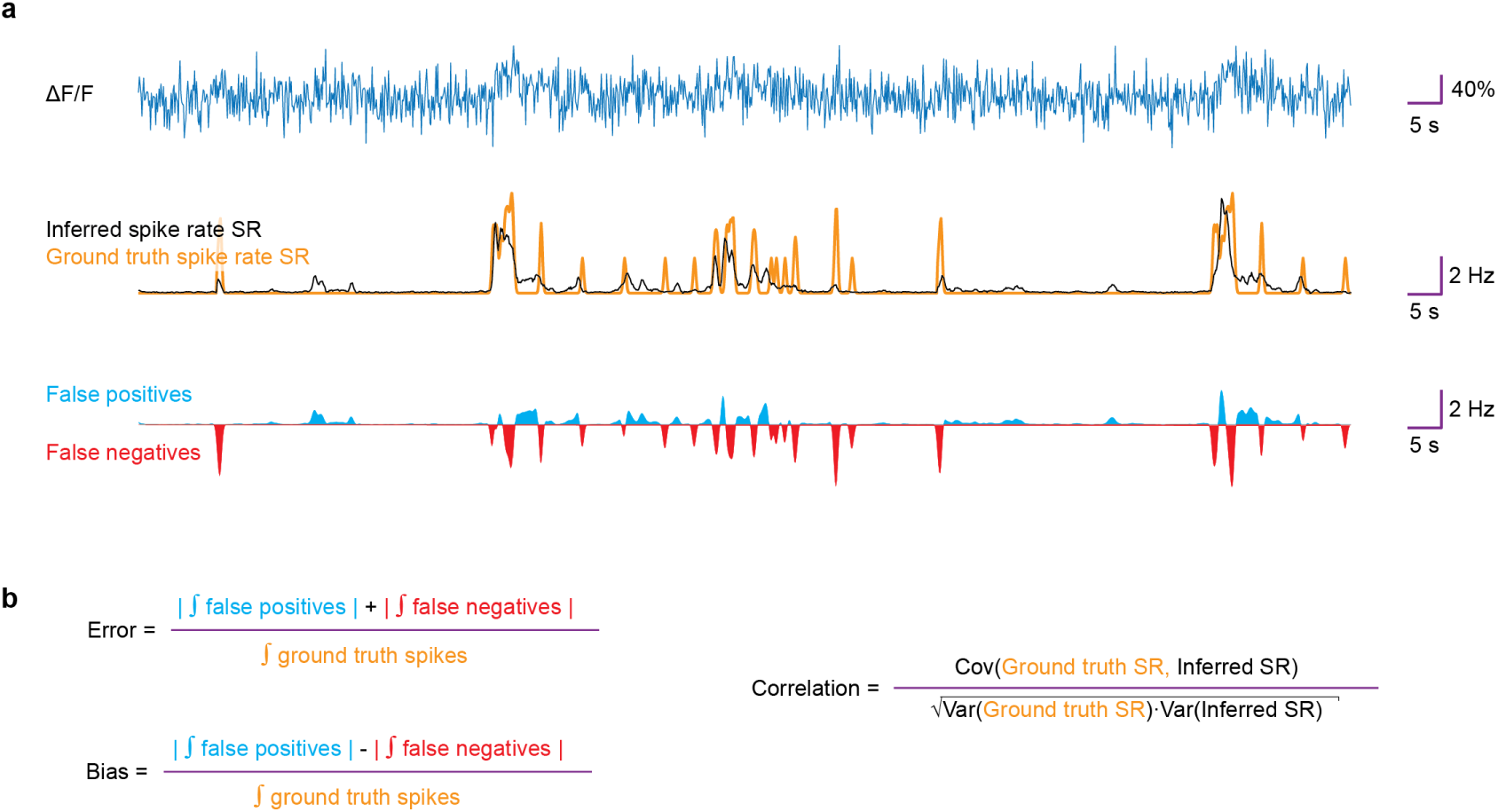
Illustration of error, bias and correlation metrics. As metrics that do not only measure the similarity (correlation) of ground truth spike rates and inferred spike rates, the error and bias also indicate absolute deviations from true spike rates. **a**, Example ΔF/F trace (dark blue), true spike rate (orange) and inferred spike rate prediction (black). The area under the spike rate traces corresponds to the number of true or inferred spikes. The difference between true and predicted spike rates gives false positives (light blue area) and false negatives (red area). **b**, The unsigned integral of the false positives and negatives, divided by the integral of true positives yields the error, while the difference yields the bias. The normalization by the integral of true positives, i.e., the true number of spikes makes errors and biases comparable across spike rate conditions. A side-effect of this normalization is that for neurons that spike only very sparsely, (relative) errors are systematically higher. Correlation is defined as the Pearson correlation coefficient computed with ground truth spike rates and inferred spike rates.

**Figure S6.**
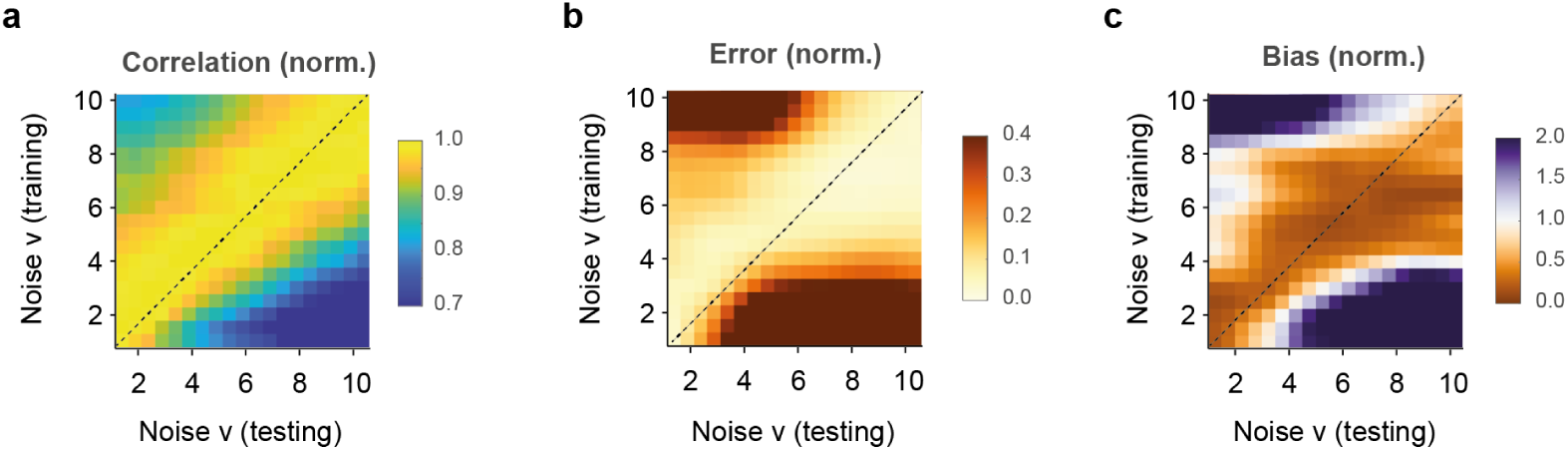
Matching standardized noise level *ν* of training and test data. Same as Fig. 2e-g, but with each column (testing level) normalized in order to highlight that the optimal training level for each testing noise level lies close to the diagonal. The correlation (a) was normalized by the maximum of each column, while error and bias metrics have been normalized by the minimum of each column. ν in units of standardized noise, % · *Hz*^−1/2^.

**Figure S7.**
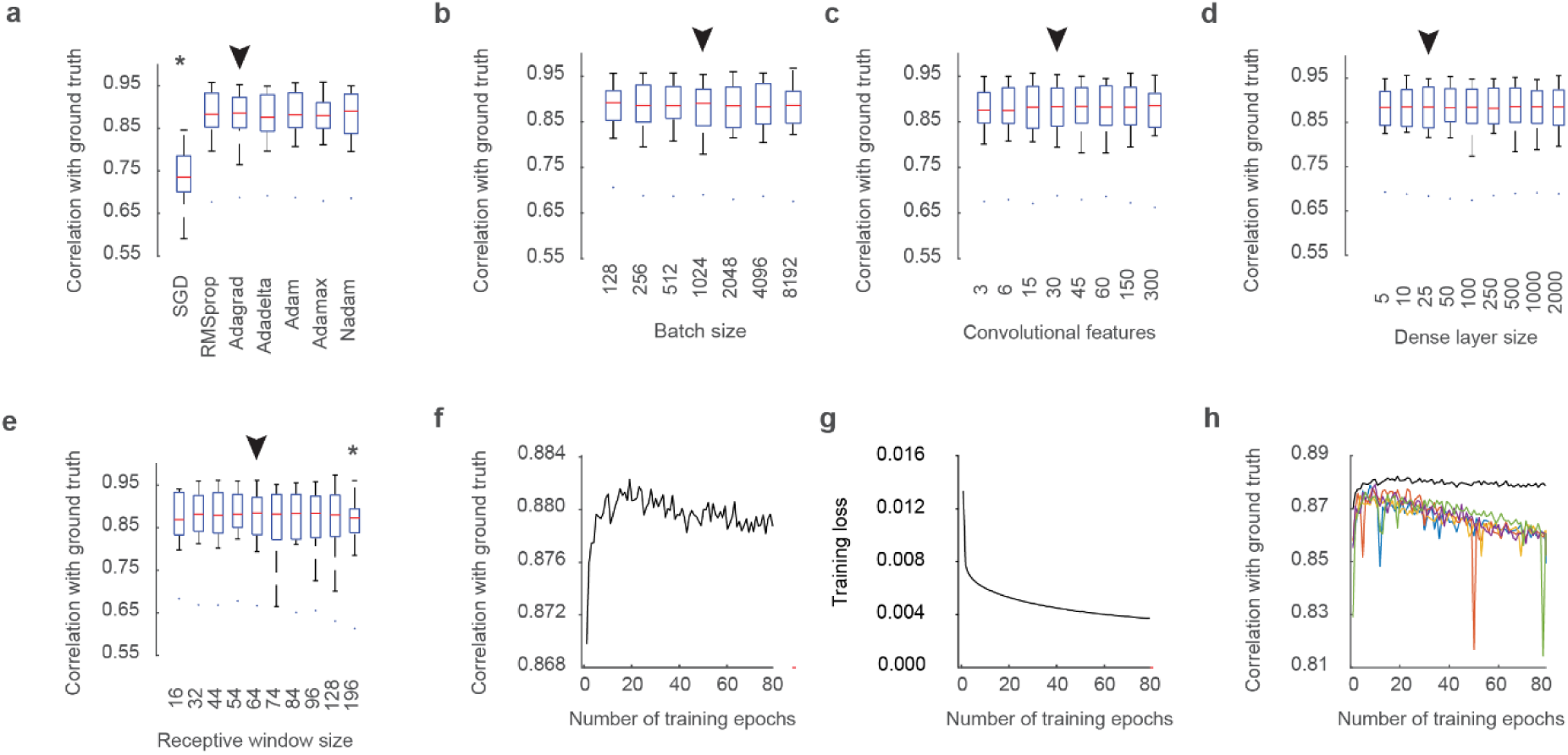
Parameter-robustness of CASCADE with respect to hyper-parameters. **a,** Performance is robust with respect to choice of the gradient descent optimizer, unless the naive stochastic gradient descent (SGD) is selected. **b,** Performance is robust with respect to the batch size used during learning. Batch sizes can influence the efficiency of gradient descent. **c,** Performance is robust against large variations in the number of features of the convolutional layers. Numbers indicate the mean number of features across the three convolutional layers. **d,** Performance is robust against large variations in the number of neurons in the dense layer. **e,** Performance is robust against large variations in the size of the receptive window; a receptive window of 64 data points corresponds *e.g.* to 64/7.5 ≈ 8.5 s in this setting. All parameter studies were performed in DS#04 at a calcium imaging rate of 7.5 Hz and a resampled noise level of 2. Networks were trained for 10 epochs with all data except for a single neuron, which was used for testing. Ensembles of 5 networks were used, and results were averaged across 3 iterations. No significant differences as observed by a Wilcoxon paired signed-rank test were found (exceptions indicated by asterisks). The parameter choices used as default values for the model are indicated by a black arrowhead. **f,** The mean correlation with the ground truth across neurons initially increases and subsequently decreases slightly during training. **g,** The training loss decreases monotonically during training. **h,** While the performance of the network is stable across epochs when using an ensemble of 5 instantiations (black), erroneous and unpredictable deviations can be observed for algorithms based on a single network (colored learning curves).

**Figure S8.**
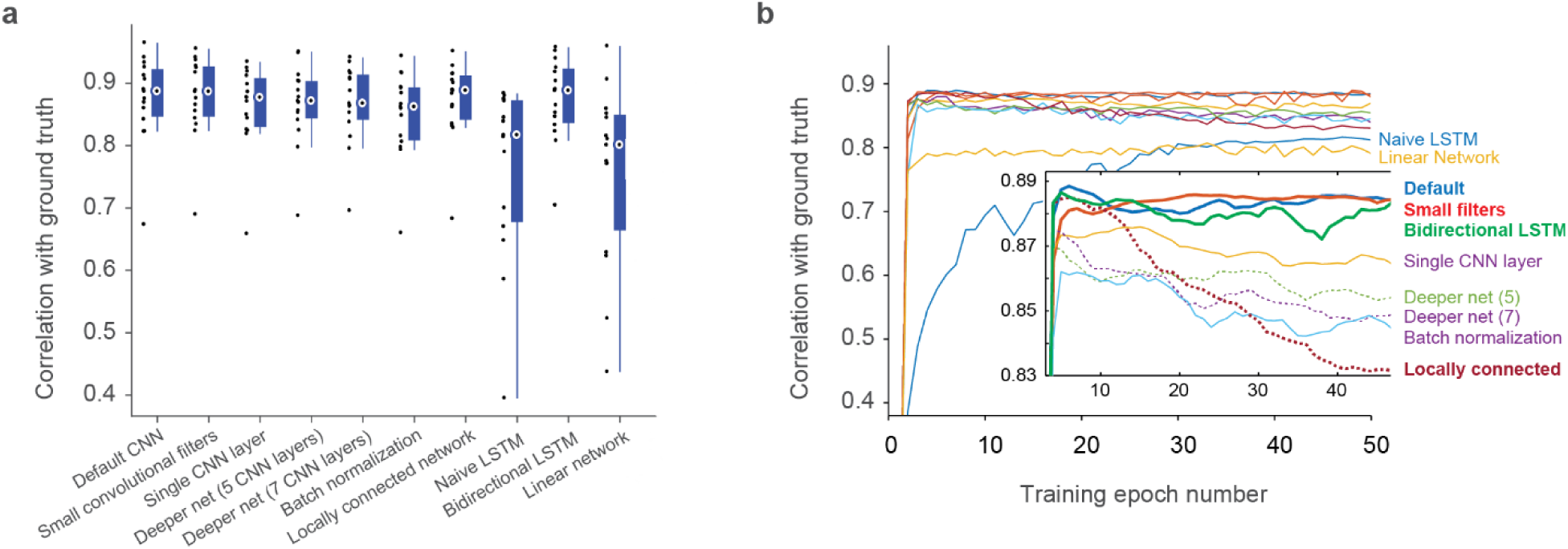
Comparison across deep learning architectures. **a,** Comparison of the default network with a network with smaller filters (Δ = -(0.002 ± 0.006), p = 0.28; paired Wilcoxon signed-rank test; pseudo-median Δ ± 95% confidence interval), with a single layer network (Δ = 0.013 ± 0.009, p = 0.008), with a deeper net (5 layers, Δ = 0.008 ± 0.011, p = 0.09), with a deeper net (7 layers, Δ = 0.008 ± 0.009, p = 0.07), with a network using batch norm (Δ = 0.023 ± 0.010, p = 0.0003), with a locally connected network (Δ = 0.002 ± 0.005, p = 0.33), with a naïve LSTM network (Δ = 0.070 ± 0.063, p = 0.00006), with a bidirectional LSTM network (Δ = 0.002 ± 0.008, p = 0.56) and with a simple linear network (Δ = 0.085 ± 0.080, p = 0.00006). Compared for a single dataset (DS#04), sampled at 7.5 Hz at a standardized noise level of 2. For detailed descriptions of all architectures, see Methods. **b,** Learning curves across epochs for all networks. The inset highlights the significant overfitting resulting from relatively large networks (deeper net 5, deeper net 7 and locally connected network; dashed lines).

**Figure S9.**
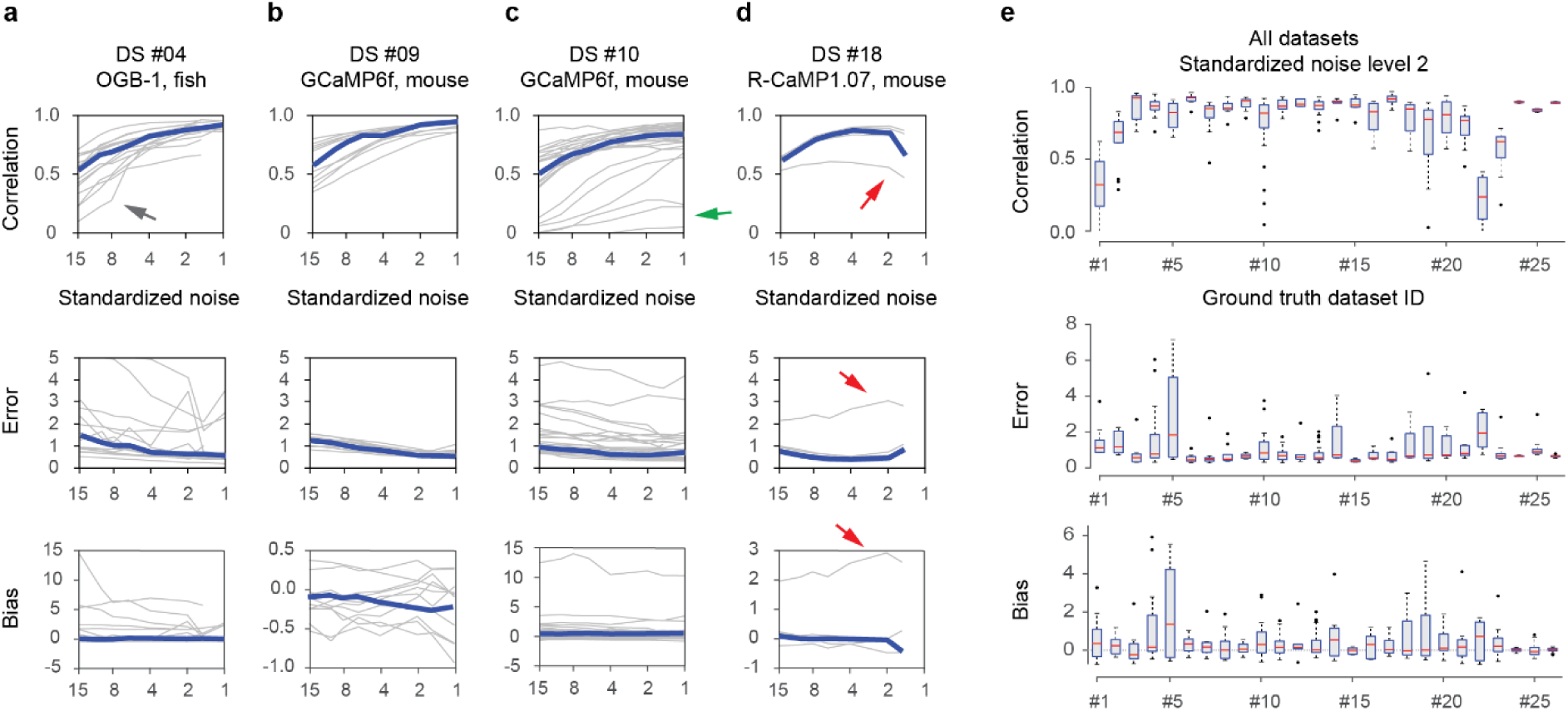
Generalization across neurons within a dataset. The deep network was trained on all neurons of a specific dataset except one, and then tested with the remaining neuron. This analysis shows how the network is able to generalize to new neurons recorded under the same conditions, as a function of the standardized noise level ν in % · *Hz*^−1/2^. **a-d,** Performance of the predictions for 4 selected ground truth datasets in terms of correlation, error and bias as a function of the standardized noise level. Error values were cropped at a value of 5 for display purposes. Single neurons in grey, median across neurons in blue. Grey lines highlighted by arrows indicate outlier neurons with particularly low spike rates (black and green arrows) and particularly distinct calcium response kernel (red arrow, see main text for discussion). **e,** Correlation, error and biases as a distribution across neurons within each dataset. All datasets were re-sampled at a frame rate of 7.5 Hz.

**Figure S10.**
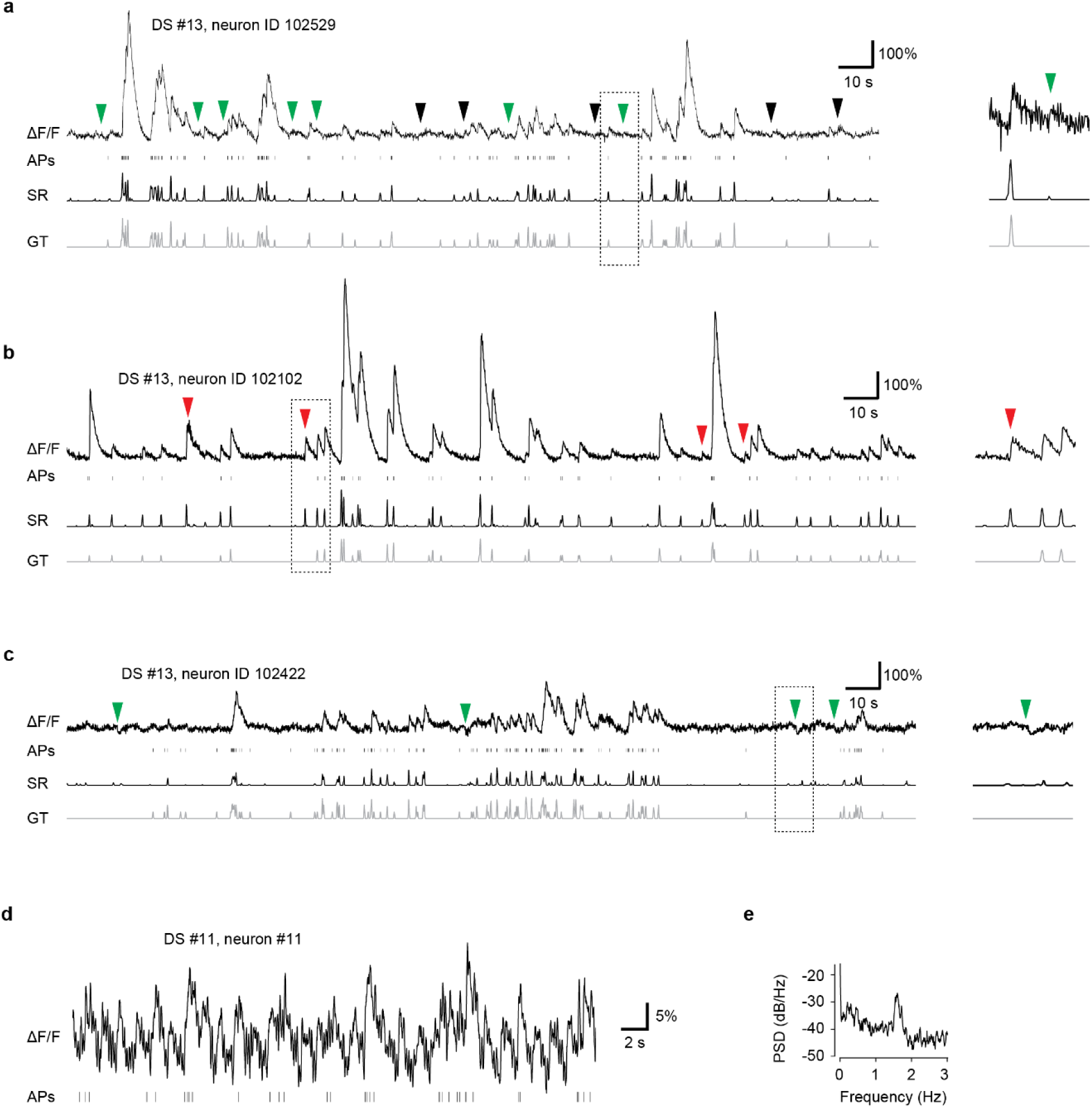
Typical artifacts in ground truth recordings. Calcium trace (ΔF/F), true action potentials (APs), inferred spiking activity (SR) and true ground truth spiking activity (GT). **a,** The baseline of this recording is unstable, exhibiting irregular bumps (arrowheads). The supervised deep network can learn to ignore these movement artifacts if their dynamics is dissimilar from the sharp onset of calcium transients. Predictions of the deep network are shown in black, ground truth in grey. Green arrowheads indicate movement artifacts that are not associated with high spiking acitivity (correct rejections of artifacts), while black arrowheads indicate movement artifacts that are not recognized as artifacts by the network (false positives). The zoom-in on the right shows an example where a movement artifact is associated with a negligeable spike rate (correct rejection). **b,** Fluorescence transients without corresponding action potentials are clearly visible (red arrowheads). These are induced by contamination through bright neuropil. The deep network is unable to distinguish this artifact from true calcium transients. **c,** Negative transients (arrowheads) are generated by standard neuropil decontamination (subtraction of the neuropil surround). The deep network can learn to partially ignore these events (correct rejections). **d,** Trace showing periodic movement artifacts that do not correspond to action potentials. **e,** A power spectral density of the recording in (d) exhibits a peak at ca. 1.5 Hz, suggesting breathing of the anesthetized animal underlying the movement artifact.

**Figure S11.**
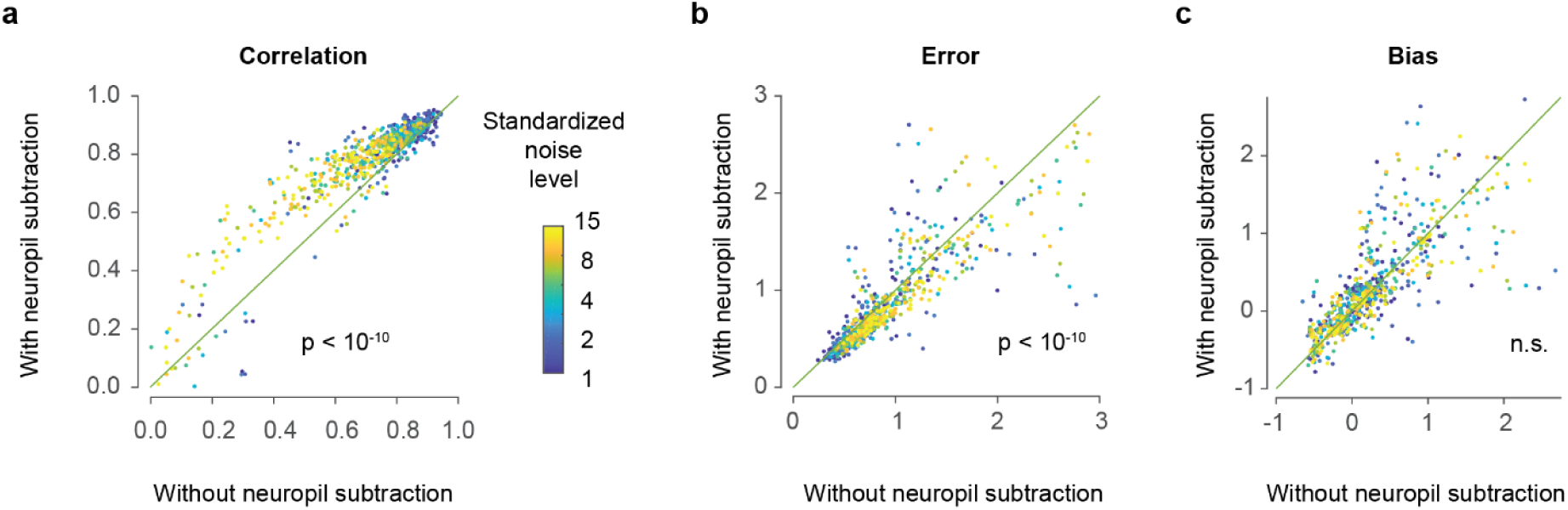
Neuropil decontamination improves spike inference. For datasets DS#10-13 (Huang et al. (2019), datasets from the Allen Institute), ground truth data were extracted both with and without simple subtractive neuropil decontamination. **a,** The performance (correlation) is improved by neuropil decontamination. The same calcium recordings were analyzed for several noise levels (color-coded). The p-value (paired signed-rank test) was <10^-10^ for every noise level but decreased for higher noise levels. **b**, Same for the error metric. Errors were significantly reduced after neuropil decontamination. **c**, Same for the bias metric. No significant effect of neuropil decontamination was observed. Color-coding indicates the standardized noise level ν (in % · *Hz*^−1/2^) for each data point.

**Figure S12.**
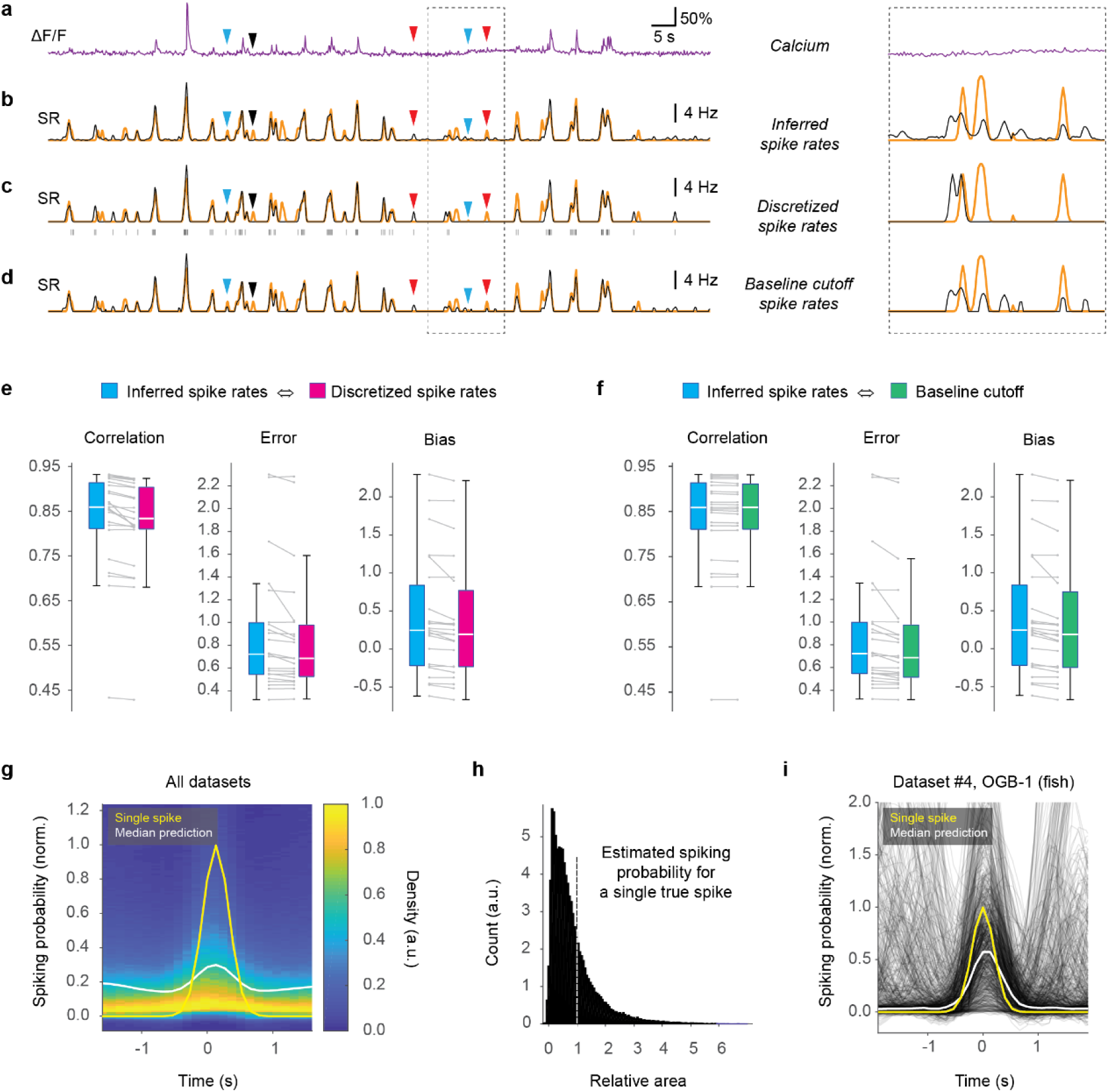
Analysis of single-spike resolution of spike inference. **a,** Calcium ΔF/F0 of an example neuron. **b,** Inferred spike rate (SR, black) and ground truth spike rate (orange). Inset to the right zooms into a section of the recording to highlight the noise floor. **c,** Discrete spikes were fitted into the inferred spiking probabilities from (b). Blue arrowheads indicate predictions that were therefore improved, red arrowheads indicate predictions that there degraded. **d,** Spiking probabilities from (b), but using a threshold (1/e of the magnitude of a single spike) as cutoff for the noise floor. **e,** Comparison of inferred probabilities (cf. (b)) vs. discretized spiking (cf. (c)). While errors and biases are slightly decreased for discretized spiking, also the correlations are reduced. Each data point is the median across a ground truth dataset. **f,** Thresholding the predictions as in (d) results in the same reduction of errors and biases, without negatively affecting the correlations. **g,** Distribution of predicted spiking probability for an isolated action potential in the ground truth across all datasets. Across all datasets, the estimate (white) of a single ground truth action potential is clearly lower than expected from ground truth (yellow). **h,** Distribution of the overall area under the curve associated with a single isolated ground truth action potential. In an ideal case, values would be narrowly distributed around 1 (dashed line). Instead, the distribution is broad and biased with a central value <1. **i,** Spiking probability for a single dataset that should be ideally qualified for single-spike precision (DS#04; no movement artifacts due to *ex vivo* preparation, synthetic indicator OGB-1). Even here, the median prediction is on average systematically lower than a single action potential, indicating that the network trades off single-spike precision in order to prevent false positive detection of spikes amidst of shot noise.

**Figure S13.**
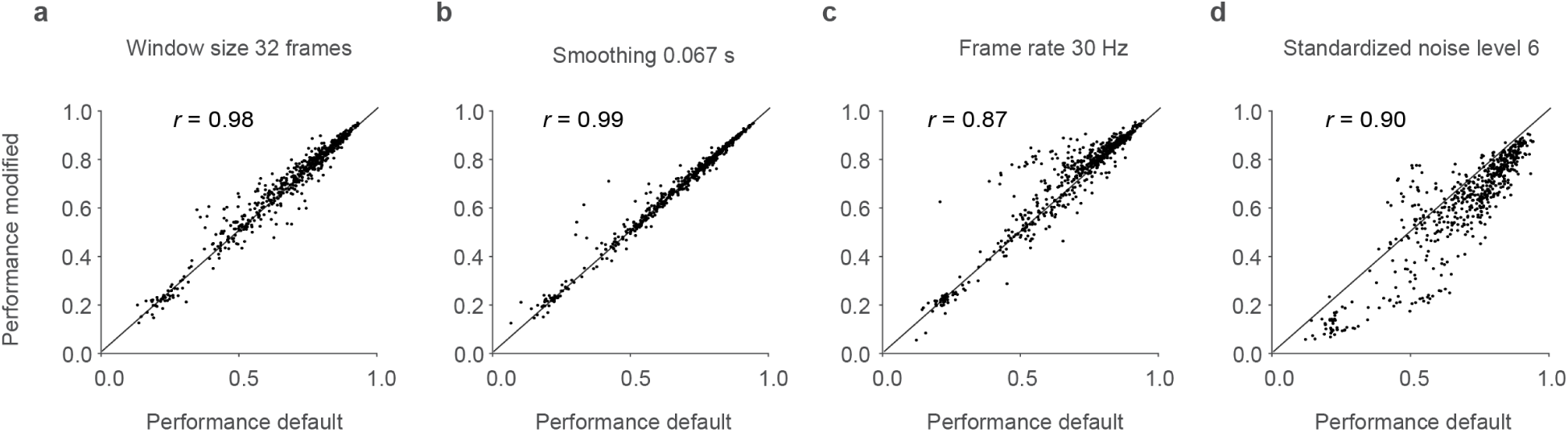
Parameter changes leave mutual predictability across datasets largely unchanged. The off-diagonal elements in Fig. 4a were recomputed using different parameter settings. The standard parameters were a smoothing kernel of 0.2 s, a framerate of 7.5 Hz, a standardized noise level of 2 and a window size of the deep network of 64 data points. The performance (correlation with ground truth) for the standard parameter analysis (“default”, x-axis) was plotted against the performance for modified parameters. Each data point corresponds to an off-diagonal matrix element in Fig. 3a. The red line indicates the unity function. Correlation between the two conditions is indicated as *r*. **a,** Window size decreased to 32. **b,** Smoothing kernel reduced to 0.067 s. **c,** Frame rate increased to 30 Hz. **d,** Standardized noise level increased to 6.

**Figure S14.**
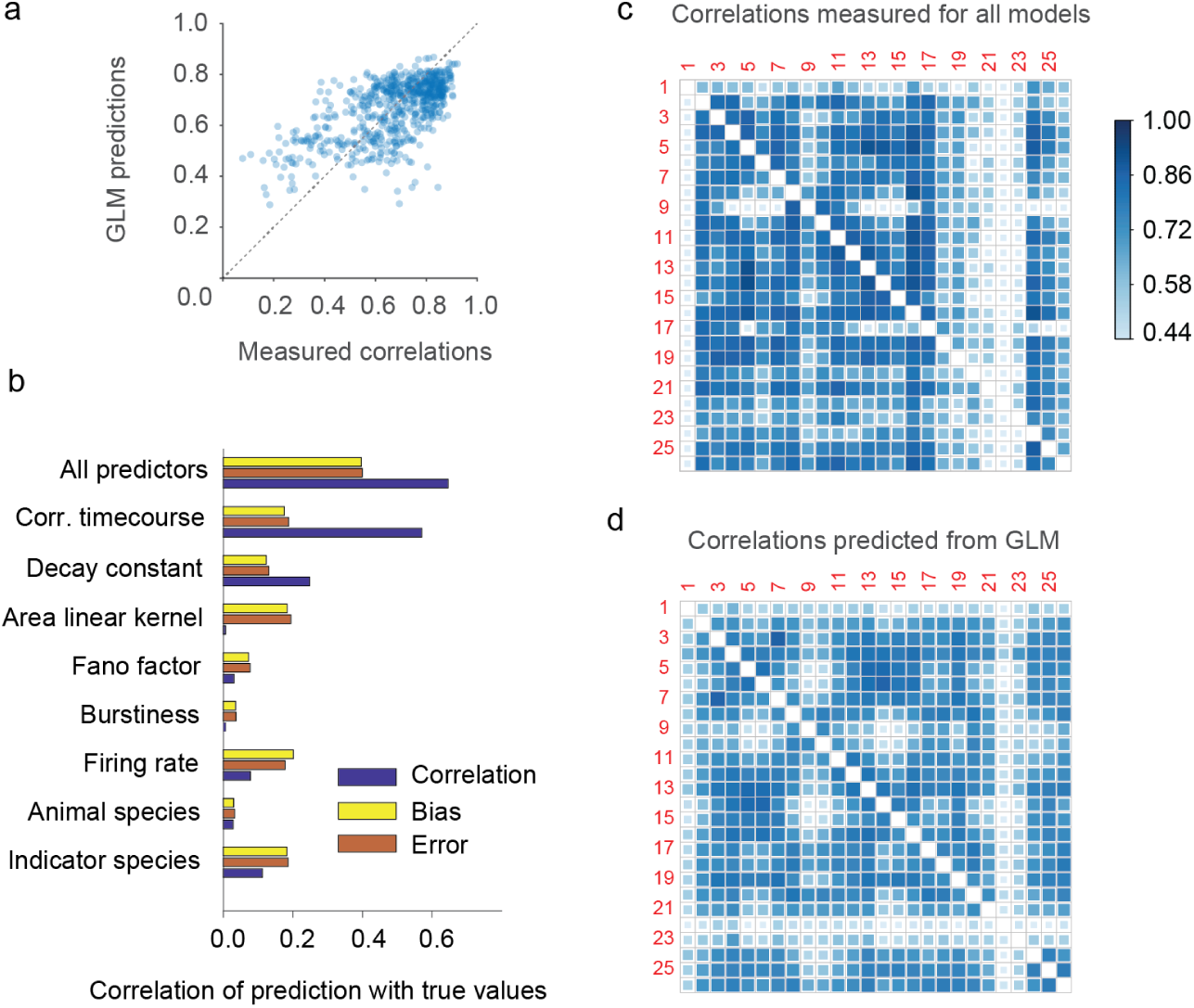
Predicting cross-dataset predictability with a generalized linear model (GLM). We considered several characteristics of ground truth datasets and evaluated whether they could improve the mutual predictability across ground truth datasets. This includes characteristics that are accessible without ground truth (indicator species, i.e., synthetic dyes vs. genetically encoded indicators; animal species, *i.e.,* mouse vs. zebrafish; median spike rate across neurons; the burstinesss; the Fano factor) and characteristics that are only accessible with available ground truth (area under the curve of the linear kernel, cf. Fig. S1; decay constant of the linear kernel; the correlation between the kernels of the training and test datasets), with detailed descriptions in the Methods section. These 8 predictors were used as regressors for a GLM to fit the mutual predictability matrix (correlations) among datasets. **a,** Correlations predicted from the GLM vs. measured correlations (see Fig. 3a). **b,** A GLM based on “all predictors” results in a correlation that recovers panel (a). In addition, this plot also shows values for fits to the bias and error matrices. Using only one of the predictors reduces the correlation, often very significantly. **c,** Measured cross-dataset predictability (reproducing Fig. 3a). **d,** Cross-dataset predictability as computed with the GLM. Together, this supplementary figure shows that the GLM is not able to explain a large fraction of the variance of the original matrix, and that the main predictor is the correlation between kernel time courses, which is not accessible without available ground truth. As consequence, we chose the use of a “global EXC model” that was trained on all reliable datasets (Fig. 3a)

**Figure S15.**
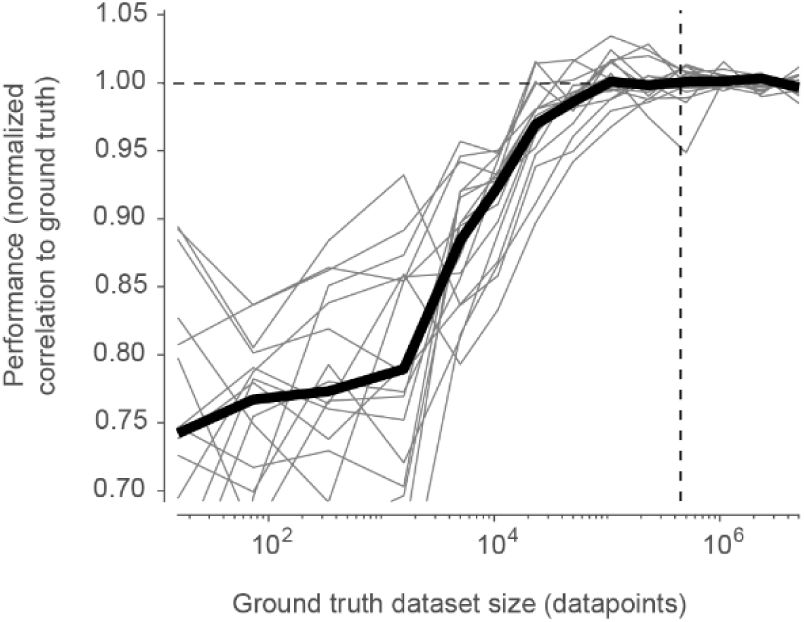
Improvement of performance with ground truth dataset size. The global EXC model (see Fig. 3) was trained as before, but using only a subset of the ground truth data points (x-axis). The performance (correlation) across each dataset was normalized to the performance with 5 million data points (horizontal dashed line). The performance approaches an asymptote at approximately 100,000 data points. A typical single ground truth dataset contains ca. 400,000 data points (median across all datasets; vertical dashed line). This result also indicates that a diverse but smaller training dataset sampled from all ground truth datasets results in better generalization than a larger training dataset from a single ground truth dataset.

**Figure S16.**
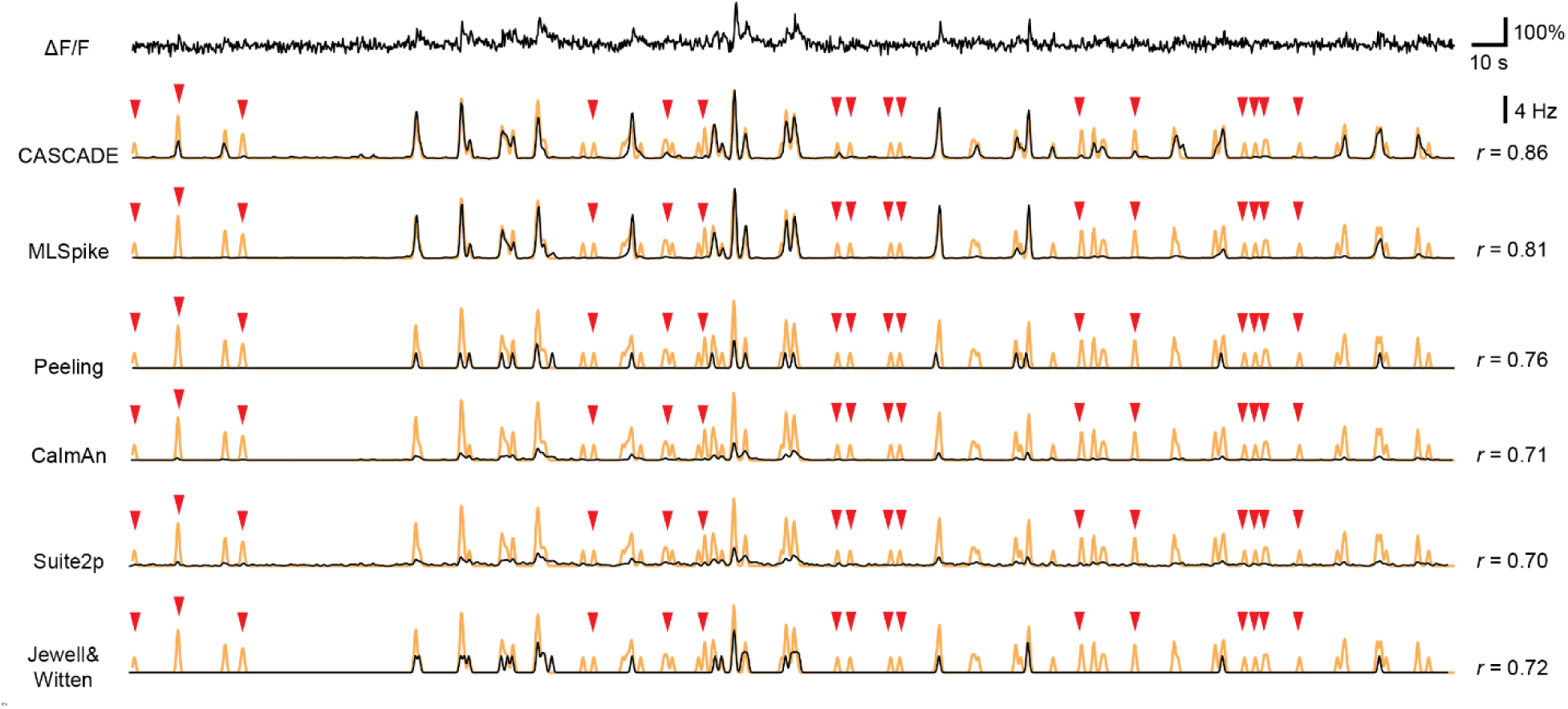
Comparison with model-based algorithms, extension of Fig. 4a. Example predictions from the deep-learning based method (CASCADE) and five model-based algorithms (MLSpike, CaImAn, Peeling, Suite2p, Jewell&Witten) of a ΔF/F recording. Inferred spike rates are in black, ground truth spike rates in orange. *r* indicates correlation of predictions with ground truth. Events that are not detected across all algorithms (false negatives) are labeled with red arrowheads. Compared to the example in Fig. 4a, the calcium recording here is rather noisy due to the insensitivity of GCaMP to single action potentials in this neuron.

**Figure S17.**
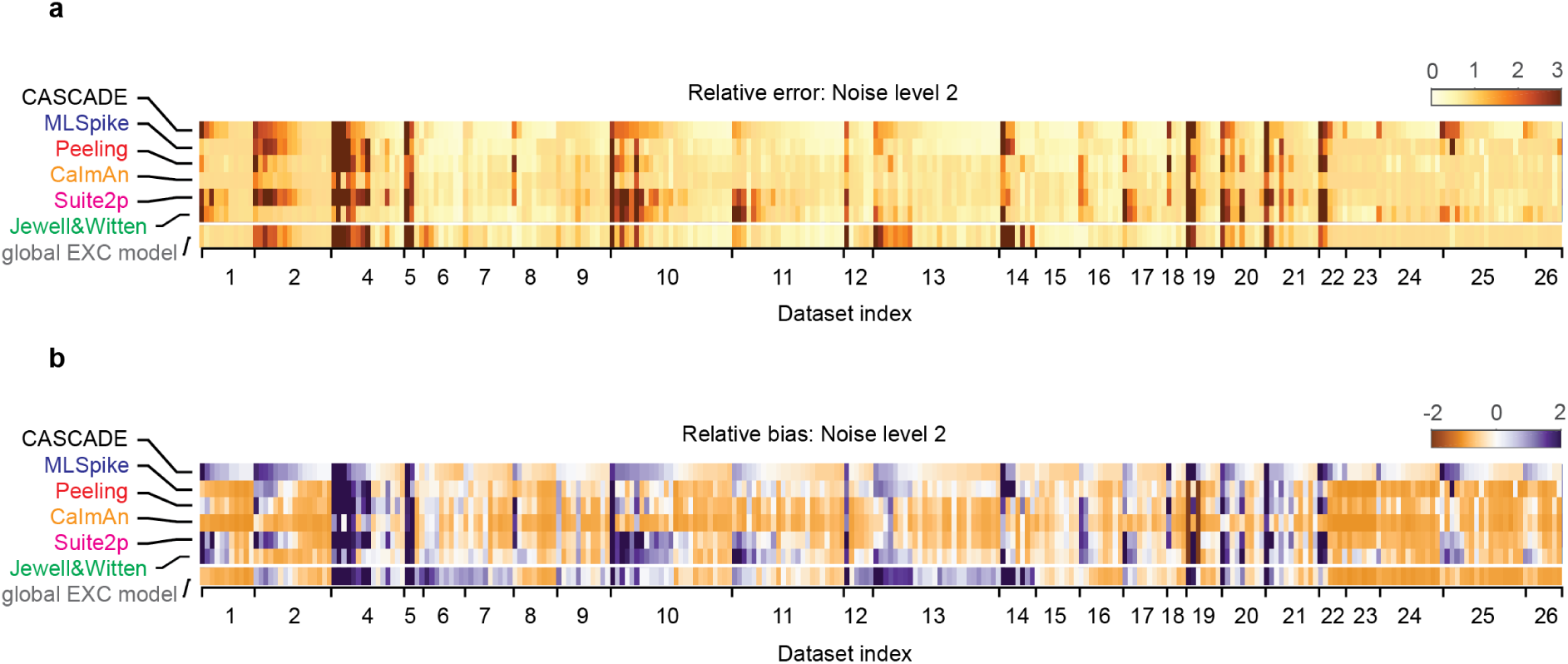
Comparison of CASCADE with model-based algorithms, extension of Fig. 4b. Comparison of the six algorithms when optimized for a single dataset, showing relative error and relative bias for all neurons, grouped by ground truth dataset.

**Figure S18.**
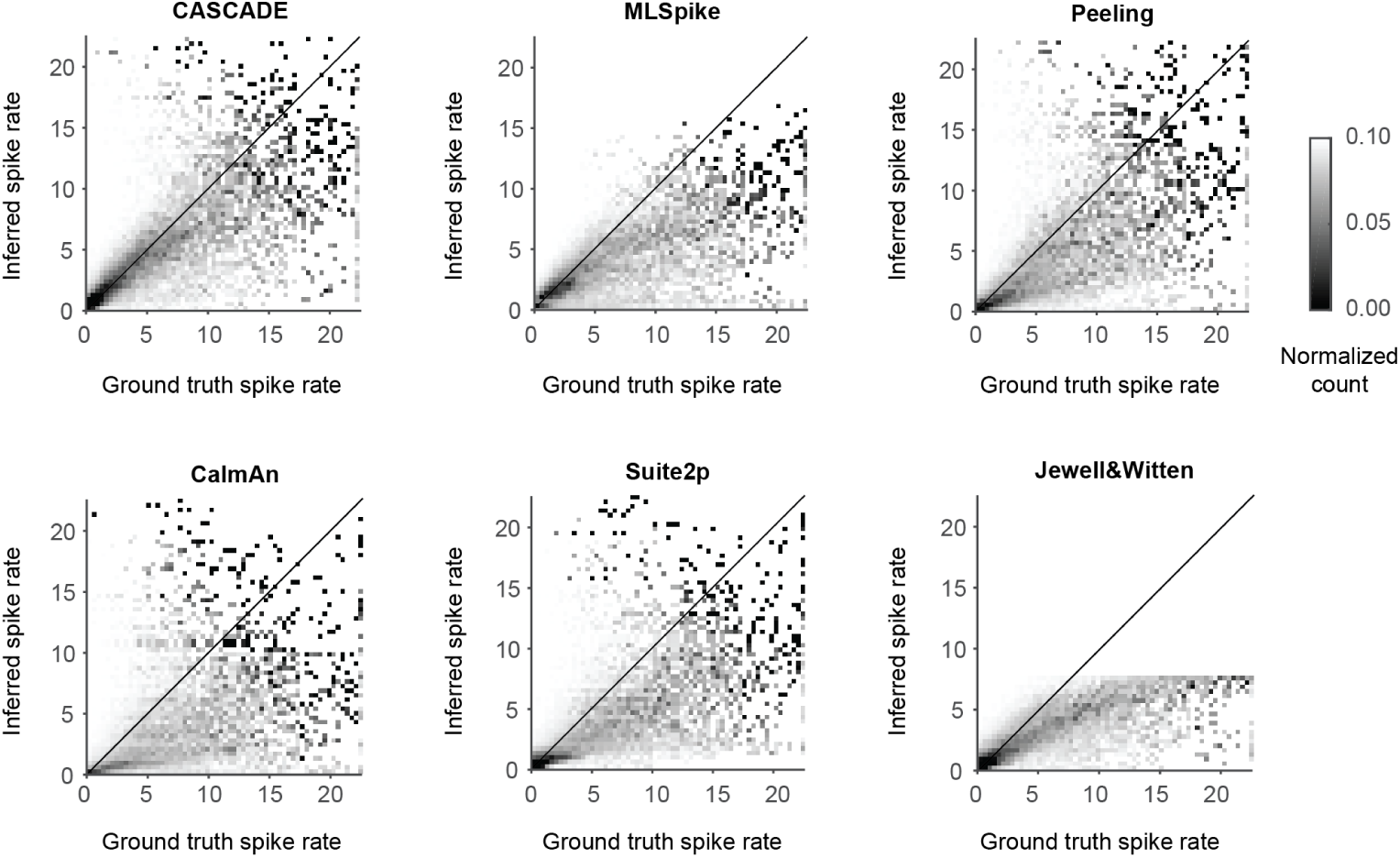
Bias of predictions across spike rates. Spike rates averaged for per 2 s-window, ground truth spike rate vs. Inferred spike rates for all algorithms, shown as a 2D histogram. The histogram was normalized to each column and each row to highlight the histogram shape also for higher spike rates. Fig. 4g shows the respective median lines across the histograms (median taken for each bin of the x-axis) for all algorithms.

**Figure S19.**
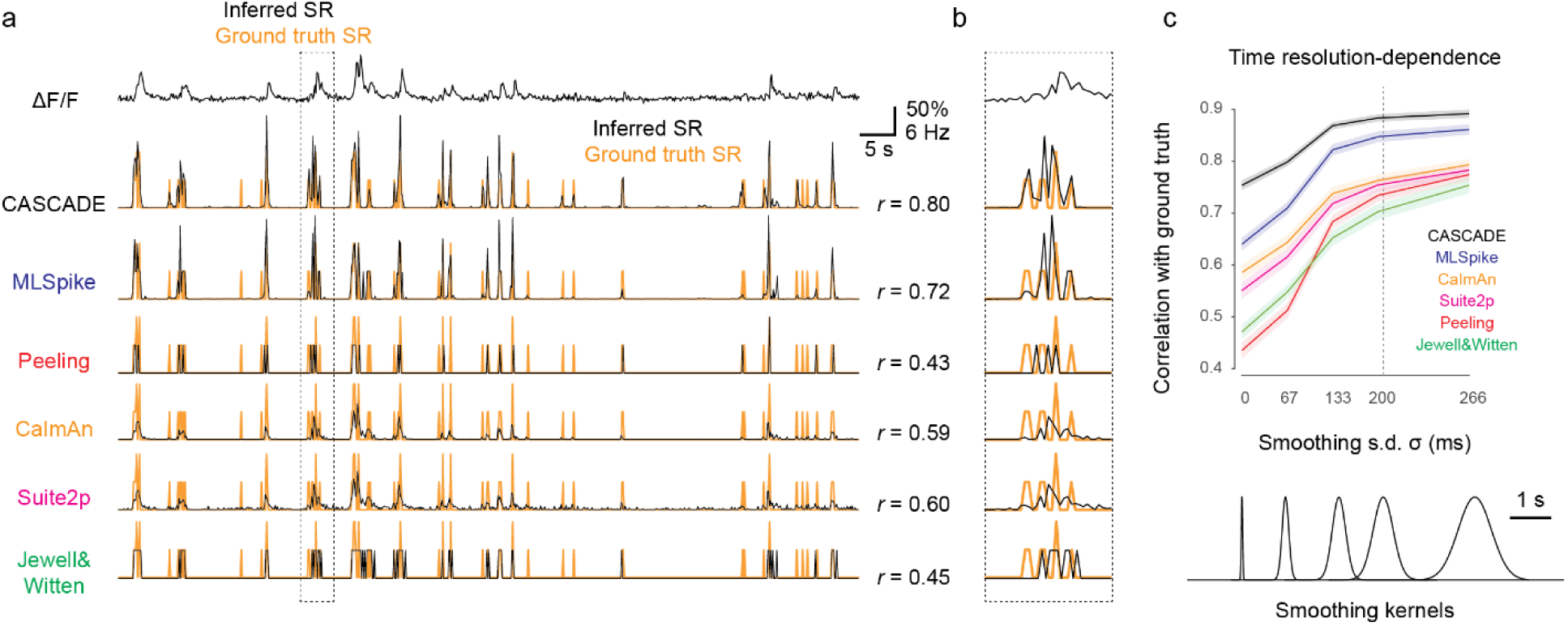
Performance dependence on temporal precision of predictions. All allgorithms were optimized via the mean squared error to infer spike rates at a specific temporal precision defined by the smoothing of the ground truth (default: Gaussian smoothing with kernel of σ = 200 ms). For all model-based algorithms, the inferred spike traces were shifted in time to optimize the mean squared error. **a**, Predictions from an example ΔF/F trace (top; dataset #09). Ground truth spike rates are shown in orange, inferred spike rates as black overlay. Correlation values are indicated at the right. The scale bars for ΔF/F and time are the same as in Fig. 4a. **b**, Highlighted excerpt from (a). Due to the high temporal precisions of the inferred spike rates, small time shifts lead to low performance (clearly visible for the Peeling algorithm in this example). The CaImAn and Suite2p algorithms deconvolve less aggressively, therefore making less dramatic errors. CASCADE and MLSpike perform best for this example neuron, with CASCADE detecting more events than MLSpike. **c**, Overall performance (correlation) change with temporal precision of predictions (smoothing kernels shown below) on a subset of datasets (datasets #4, #6, #9, #11-14 and #18). As expected, correlation with ground truth decreased with higher temporal resolution of the desired temporal resolution. This decrease was especially prominent for algorithms that, by design, aim at the inference of precise (discrete) spike rates (Peeling, Jewell&Witten). The decrease was less pronounced for CASCADE compared to *e.g.* MLSpike. All recordings resampled at a noise level of 2 with a frame rate of 7.5 Hz.

**Figure S20.**
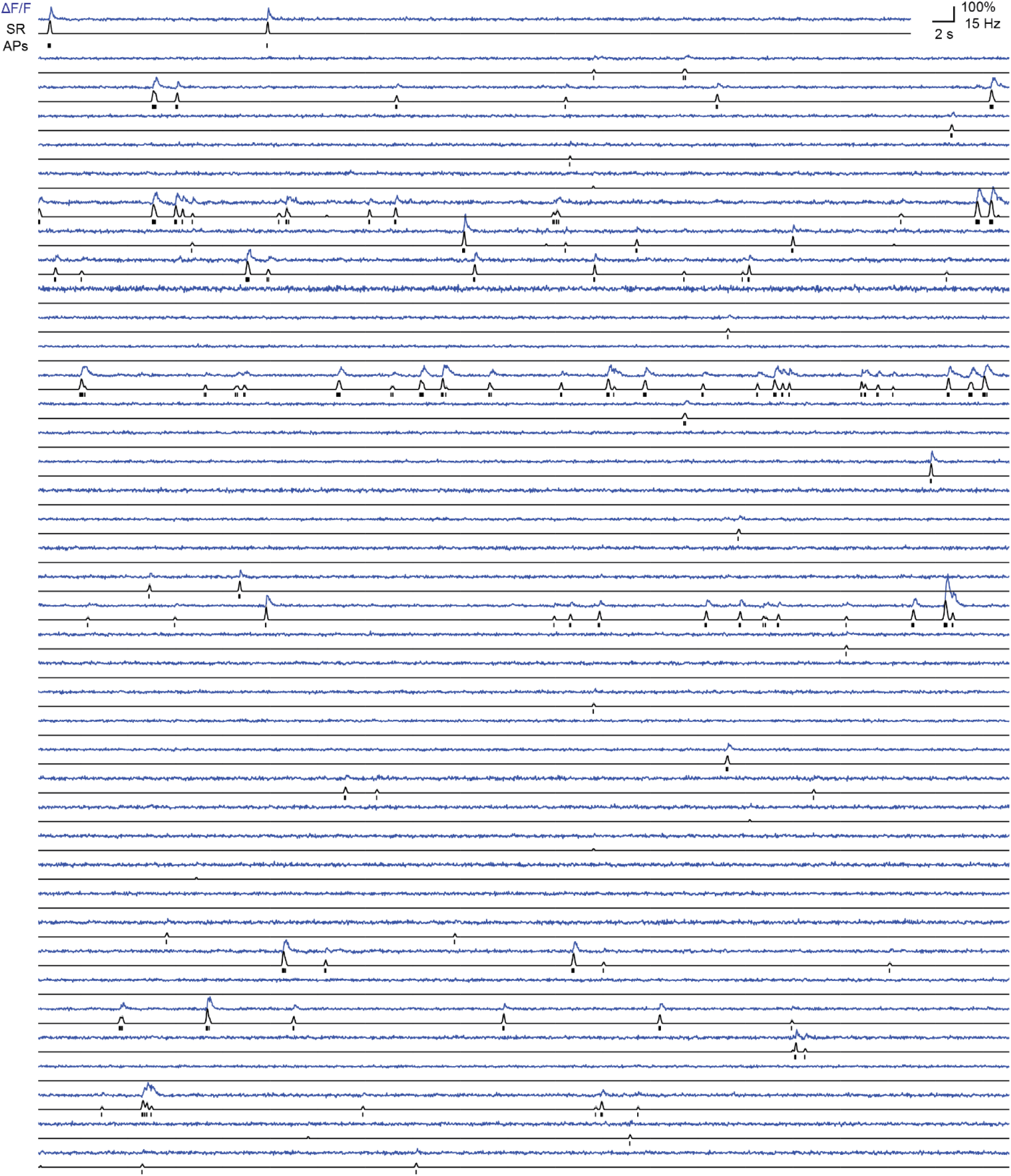
Predictions of spiking probabilities and discrete spikes from the Allen Brain Institute Visual Coding dataset. Predictions were produced with the global EXC model trained at 30 Hz. From dataset ID ‘552195520’, plotting a total of 40 neurons out of 74, approximately 1 minute out of 63.2 minutes of recording for this dataset. Discrete spikes are the most likely fit, generated with an algorithm using Metropolis-Monte Carlo sampling as starting point (see Methods).

**Figure S21.**
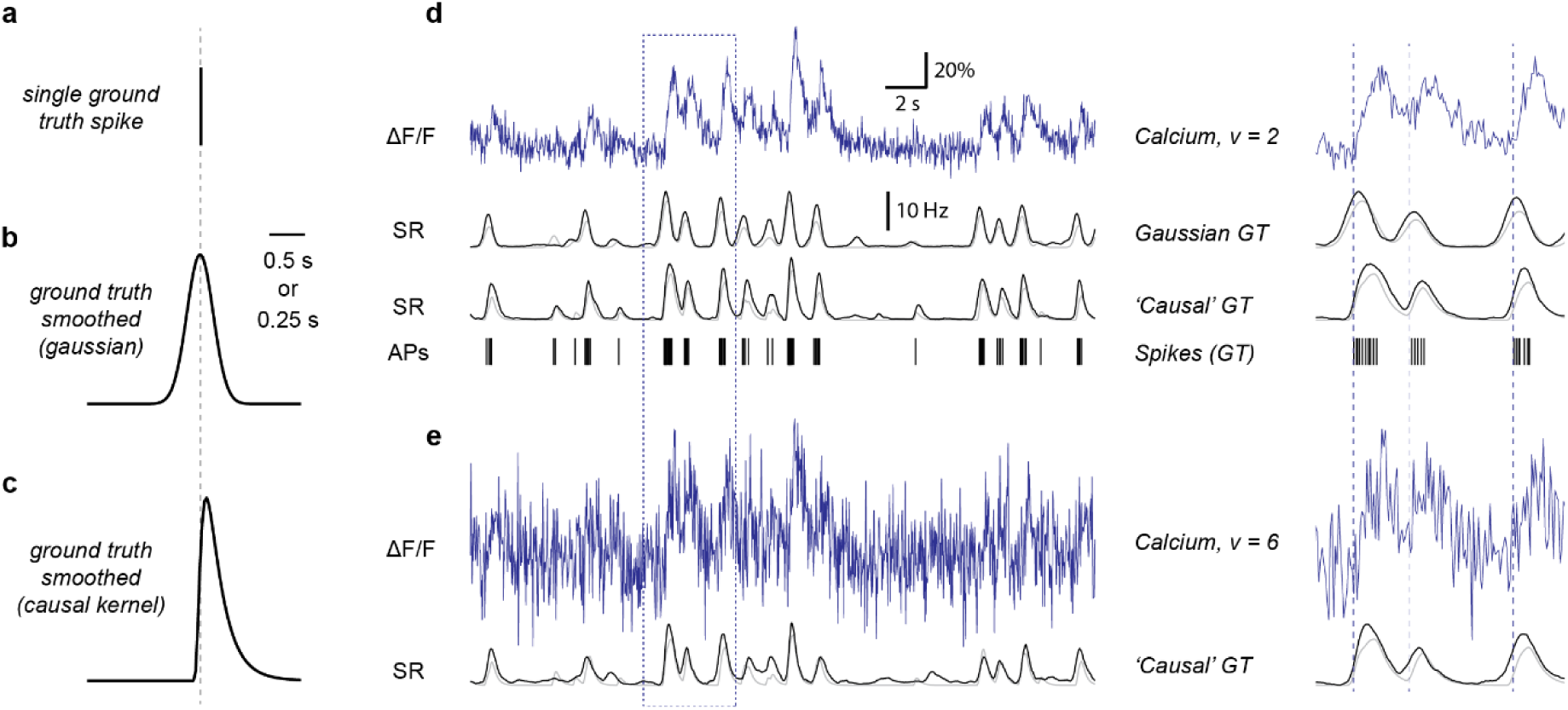
Non-Gaussian ground truth smoothing kernel. Since calcium signals rise after action potentials, the task for spike inference is to re-assign ΔF/F activity to the true spike time. Due to noise and imperfect match between model and calcium trace, the re-assignment is typically slightly off, into both the past and the future of the true spike. In principle, both re-assignment to past and future is equally undesirable. In some scenarios, however, re-assignment of the activity into the past of the spike can be particularly adverse, for example, when a precise external stimulus (*e.g.*, an auditory tone) is used to generate a peri-stimulus activity trace. While the ground truth used for training **(a)** is typically smoothed symmetrically in time with a Gaussian function **(b)**, by design activity that happened briefly after the stimulus will be deconvolved to time points both after and before the stimulus. To circumvent these a-causal events, it is possible to use a more causal filter **(c)**, using a highly skewed inverse Gaussian distribution. **d,** The deep network can use the ‘causal’ ground truth, resulting in inferred spike rates that re-assign activity in a more causal way (lower prediction trace; see zoom-in). However, noise sources result in re-assignment of activity to time bins prior to ground truth spikes. **e,** The predictions become necessarily more sloppy and smeared out for higher noise levels. This shows that even when using a causal kernel to smooth the ground truth, a-causal effects can be induced by spike inference. Depending on the desired temporal resolution of the algorithm, different width of the smoothing kernels can be chosen. A Gaussian with σ = 0.2 s will lead to a FWHM of the smoothing Gaussian of ca. 0.5 s (b). For panel (d), a smoothing kernel with σ = 0.2 s was used.

**Figure S22.**
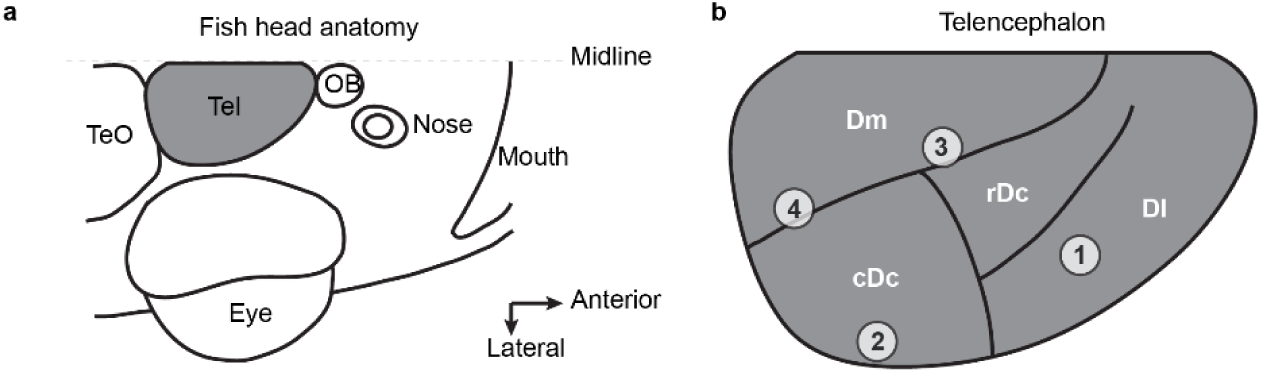
Locations of neurons in the dorsal telencephalon of adult zebrafish from ground truth dataset DS#08. **a,** Fish head anatomy, highlighting the Telencephalon from a dorsal aspect. TeO = optic tectum, OB = olfactory bulb, Tel = Telencephalon. **b,** Recording locations in the telencephalon (see Methods for details). Neuron #1 of the ground truth dataset was located in Dl, ca. 120 μm below the dorsal surface of Dm (position 1). Neurons #2-#5 were located in cDc, ca. 50 μm below the dorsal surface of Dm (position 2). Neurons #6 and #7 were located close to the sulcus ypsilonformis at the interface of Dm and rDc, at a depth of ca. 80 μm below the dorsal surface of Dm (location 3). Neurons #8, #9 and #10 were located at the interface of Dm and cDc, ca. 20-30 μm below the dorsal surface of Dm. Nomenclature and subdivisions follow Huang et al. (2020).

## REFERENCES

1. Göbel, W. & Helmchen, F. In Vivo Calcium Imaging of Neural Network Function. Physiology 22, 358–365 (2007).

2. Harris, K. D., Quiroga, R. Q., Freeman, J. & Smith, S. L. Improving data quality in neuronal population recordings. Nat. Neurosci. 19, 1165–1174 (2016).

3. Rose, T., Goltstein, P. M., Portugues, R. & Griesbeck, O. Putting a finishing touch on GECIs. Front. Mol. Neurosci. 7, (2014).

4. Sabatini, B. L. The impact of reporter kinetics on the interpretation of data gathered with fluorescent reporters. bioRxiv 834895 (2019) doi:10.1101/834895.

5. Wei, Z. et al. A comparison of neuronal population dynamics measured with calcium imaging and electrophysiology. PLOS Comput. Biol. 16, e1008198 (2020).

6. Yaksi, E. & Friedrich, R. W. Reconstruction of firing rate changes across neuronal populations by temporally deconvolved Ca2+ imaging. Nat. Methods 3, 377–383 (2006).

7. Greenberg, D. S., Houweling, A. R. & Kerr, J. N. D. Population imaging of ongoing neuronal activity in the visual cortex of awake rats. Nat. Neurosci. 11, 749–751 (2008).

8. Vogelstein, J. T. et al. Spike inference from calcium imaging using sequential Monte Carlo methods. Biophys. J. 97, 636–655 (2009).

9. Vogelstein, J. T. et al. Fast Nonnegative Deconvolution for Spike Train Inference From Population Calcium Imaging. J. Neurophysiol. 104, 3691–3704 (2010).

10. Lütcke, H., Gerhard, F., Zenke, F., Gerstner, W. & Helmchen, F. Inference of neuronal network spike dynamics and topology from calcium imaging data. Front. Neural Circuits 7, (2013).

11. Deneux, T. et al. Accurate spike estimation from noisy calcium signals for ultrafast three-dimensional imaging of large neuronal populations in vivo. Nat. Commun. 7, 1–17 (2016).

12. Greenberg, D. S., et al. Accurate action potential inference from a calcium sensor protein through biophysical modeling. Preprint at www.biorxiv.org/content/10.1101/479055v1 (2018).

13. Pachitariu, M., Stringer, C. & Harris, K. D. Robustness of Spike Deconvolution for Neuronal Calcium Imaging. J. Neurosci. 38, 7976–7985 (2018).

14. Friedrich, J., Zhou, P. & Paninski, L. Fast online deconvolution of calcium imaging data. PLOS Comput. Biol. 13, e1005423 (2017).

15. Berens, P. et al. Community-based benchmarking improves spike rate inference from two-photon calcium imaging data. PLoS Comput. Biol. 14, (2018).

16. Jewell, S. & Witten, D. Exact spike inference via L0 optimization. Ann. Appl. Stat. 12, 2457–2482 (2018).

17. Sasaki, T., Takahashi, N., Matsuki, N. & Ikegaya, Y. Fast and Accurate Detection of Action Potentials From Somatic Calcium Fluctuations. J. Neurophysiol. 100, 1668–1676 (2008).

18. Theis, L. et al. Benchmarking Spike Rate Inference in Population Calcium Imaging. Neuron 90, 471–482 (2016).

19. Sebastian, J., Sur, M., Murthy, H. A. & Magimai.-Doss, M. Signal-to-signal networks for improved spike estimation from calcium imaging data. Preprint at www.biorxiv.org/content/10.1101/2020.05.01.071993v1 (2020).

20. Hoang, H. et al. Improved hyperacuity estimation of spike timing from calcium imaging. Sci. Rep. 10, 17844 (2020).

21. Éltes, T., Szoboszlay, M., Kerti-Szigeti, K. & Nusser, Z. Improved spike inference accuracy by estimating the peak amplitude of unitary [Ca2+] transients in weakly GCaMP6f-expressing hippocampal pyramidal cells. J. Physiol. 597, 2925–2947 (2019).

22. Evans, M. H., Petersen, R. S. & Humphries, M. D. On the use of calcium deconvolution algorithms in practical contexts. Preprint at www.biorxiv.org/content/10.1101/871137v1 (2019).

23. Zhu, P., Fajardo, O., Shum, J., Zhang Schärer, Y.-P. & Friedrich, R. W. High-resolution optical control of spatiotemporal neuronal activity patterns in zebrafish using a digital micromirror device. Nat. Protoc. 7, 1410–1425 (2012).

24. Schoenfeld, G., Carta, S., Rupprecht, P., Ayaz, A. & Helmchen, F. In vivo calcium imaging of CA3 pyramidal neuron populations in adult mouse hippocampus. Preprint at https://www.biorxiv.org/content/10.1101/2021.01.21.427642v1 (2021).

25. Bethge, P. et al. An R-CaMP1.07 reporter mouse for cell-type-specific expression of a sensitive red fluorescent calcium indicator. PLOS ONE 12, e0179460 (2017).

26. Tada, M., Takeuchi, A., Hashizume, M., Kitamura, K. & Kano, M. A highly sensitive fluorescent indicator dye for calcium imaging of neural activity in vitro and in vivo. Eur. J. Neurosci. 39, 1720–1728 (2014).

27. Khan, A. G. et al. Distinct learning-induced changes in stimulus selectivity and interactions of GABAergic interneuron classes in visual cortex. Nat. Neurosci. 21, 851–859 (2018).

28. Kwan, A. C. & Dan, Y. Dissection of cortical microcircuits by single-neuron stimulation in vivo. Curr. Biol. CB 22, 1459–1467 (2012).

29. Huang, L., et al. Relationship between spiking activity and simultaneously recorded fluorescence signals in transgenic mice expressing GCaMP6. Preprint at www.biorxiv.org/content/10.1101/788802v1 (2019).

30. Ledochowitsch, P., et al. On the correspondence of electrical and optical physiology in in vivo population-scale two-photon calcium imaging. Preprint at https://www.biorxiv.org/content/10.1101/800102v1 (2019).

31. Dana, H. et al. Sensitive red protein calcium indicators for imaging neural activity. eLife 5, e12727 (2016).

32. Akerboom, J. et al. Optimization of a GCaMP calcium indicator for neural activity imaging. J. Neurosci. Off. J. Soc. Neurosci. 32, 13819–13840 (2012).

33. Chen, T.-W. et al. Ultrasensitive fluorescent proteins for imaging neuronal activity. Nature 499, 295–300 (2013).

34. Russakovsky, O. et al. ImageNet Large Scale Visual Recognition Challenge. Int. J. Comput. Vis. 115, 211–252 (2015).

35. Lecun, Y., Bottou, L., Bengio, Y. & Haffner, P. Gradient-based learning applied to document recognition. Proc. IEEE 86, 2278– 2324 (1998).

36. Sinz, F. H., Pitkow, X., Reimer, J., Bethge, M. & Tolias, A. S. Engineering a Less Artificial Intelligence. Neuron 103, 967–979 (2019).

37. Mathis, A. et al. DeepLabCut: markerless pose estimation of user-defined body parts with deep learning. Nat. Neurosci. 21, 1281–1289 (2018).

38. Nath, T. et al. Using DeepLabCut for 3D markerless pose estimation across species and behaviors. Nat. Protoc. 14, 2152–2176 (2019).

39. Deng, J. et al. ImageNet: A large-scale hierarchical image database. in 2009 IEEE Conference on Computer Vision and Pattern Recognition 248–255 (2009). doi:10.1109/CVPR.2009.5206848.

40. Denis, J., Dard, R. F., Quiroli, E., Cossart, R. & Picardo, M. A. DeepCINAC: A Deep-Learning-Based Python Toolbox for Inferring Calcium Imaging Neuronal Activity Based on Movie Visualization. eNeuro 7, (2020).

41. Gauthier, J. L., et al. Detecting and Correcting False Transients in Calcium Imaging. Preprint at www.biorxiv.org/content/10.1101/473470v1 (2018).

42. Giovannucci, A. et al. CaImAn an open source tool for scalable calcium imaging data analysis. eLife 8, e38173 (2019).

43. Keemink, S. W. et al. FISSA: A neuropil decontamination toolbox for calcium imaging signals. Sci. Rep. 8, 3493 (2018).

44. Ali, F. & Kwan, A. C. Interpreting in vivo calcium signals from neuronal cell bodies, axons, and dendrites: a review. Neurophotonics 7, (2020).

45. Charles, A. S., Song, A., Gauthier, J. L., Pillow, J. W. & Tank, D. W. Neural Anatomy and Optical Microscopy (NAOMi) Simulation for evaluating calcium imaging methods. Preprint at https://www.biorxiv.org/content/10.1101/726174v1 (2019).

46. Pachitariu, M., et al. Suite2p: beyond 10,000 neurons with standard two-photon microscopy. Preprint at www.biorxiv.org/content/10.1101/061507v2 (2017).

47. Jewell, S., Hocking, T. D., Fearnhead, P. & Witten, D. Fast Nonconvex Deconvolution of Calcium Imaging Data. Biostatistics (2019).

48. McMahon, S. M. & Jackson, M. B. An Inconvenient Truth: Calcium Sensors Are Calcium Buffers. Trends Neurosci. 41, 880– 884 (2018).

49. Rupprecht, P., Prendergast, A., Wyart, C. & Friedrich, R. W. Remote z-scanning with a macroscopic voice coil motor for fast 3D multiphoton laser scanning microscopy. Biomed. Opt. Express 7, 1656–1671 (2016).

50. Blumhagen, F. et al. Neuronal filtering of multiplexed odour representations. Nature 479, 493–498 (2011).

51. Rupprecht, P. & Friedrich, R. W. Precise Synaptic Balance in the Zebrafish Homolog of Olfactory Cortex. Neuron 100, 669–683.e5 (2018).

52. Mackevicius, E. L. et al. Unsupervised discovery of temporal sequences in high-dimensional datasets, with applications to neuroscience. eLife 8, e38471 (2019).

53. de Vries, S. E. J. et al. A large-scale standardized physiological survey reveals functional organization of the mouse visual cortex. Nat. Neurosci. 23, 138–151 (2020).

54. Lin, I.-C., Okun, M., Carandini, M. & Harris, K. D. The Nature of Shared Cortical Variability. Neuron 87, 644–656 (2015).

55. Moreno-Bote, R. et al. Information-limiting correlations. Nat. Neurosci. 17, 1410–1417 (2014).

56. Williams, A. H. et al. Unsupervised Discovery of Demixed, Low-Dimensional Neural Dynamics across Multiple Timescales through Tensor Component Analysis. Neuron 98, 1099–1115.e8 (2018).

57. Kaifosh, P., Zaremba, J. D., Danielson, N. B. & Losonczy, A. SIMA: Python software for analysis of dynamic fluorescence imaging data. *Front*. Neuroinformatics 8, (2014).

58. Siegle, J. H., et al. Reconciling functional differences in populations of neurons recorded with two-photon imaging and electrophysiology. Preprint at https://www.biorxiv.org/content/10.1101/2020.08.10.244723v1 (2020).

59. Vanwalleghem, G., Constantin, L. & Scott, E. K. Calcium Imaging and the Curse of Negativity. Front. Neural Circuits 14, (2021).

60. Kay, K. et al. Constant Sub-second Cycling between Representations of Possible Futures in the Hippocampus. Cell 180, 552–567.e25 (2020).

61. Bourg, A. van der et al. Temporal refinement of sensory-evoked activity across layers in developing mouse barrel cortex. Eur. J. Neurosci. 50, 2955–2969 (2019).

62. Chen, I.-W., Papagiakoumou, E. & Emiliani, V. Towards circuit optogenetics. Curr. Opin. Neurobiol. 50, 179–189 (2018).

63. Packer, A. M., Russell, L. E., Dalgleish, H. W. P. & Häusser, M. Simultaneous all-optical manipulation and recording of neural circuit activity with cellular resolution in vivo. Nat. Methods 12, 140–146 (2015).

64. Pégard, N. C. et al. Three-dimensional scanless holographic optogenetics with temporal focusing (3D-SHOT). Nat. Commun. 8, 1228 (2017).

65. Griffiths, V. A. et al. Real-time 3D movement correction for two-photon imaging in behaving animals. Nat. Methods 1–8 (2020) doi:10.1038/s41592-020-0851-7.

66. Inoue, M. et al. Rational Engineering of XCaMPs, a Multicolor GECI Suite for In Vivo Imaging of Complex Brain Circuit Dynamics. Cell 177, 1346–1360.e24 (2019).

67. Frank, T., Mönig, N. R., Satou, C., Higashijima, S. & Friedrich, R. W. Associative conditioning remaps odor representations and modifies inhibition in a higher olfactory brain area. Nat. Neurosci. 22, 1844–1856 (2019).

68. Kitamura, K., Judkewitz, B., Kano, M., Denk, W. & Häusser, M. Targeted patch-clamp recordings and single-cell electroporation of unlabeled neurons in vivo. Nat. Methods 5, 61–67 (2008).

69. Perkins, K. L. Cell-attached voltage-clamp and current-clamp recording and stimulation techniques in brain slices. J. Neurosci. Methods 154, 1–18 (2006).

70. Pologruto, T. A., Sabatini, B. L. & Svoboda, K. ScanImage: Flexible software for operating laser scanning microscopes. Biomed. Eng. OnLine 2, 13 (2003).

71. Suter, B. A., et al. Ephus: Multipurpose Data Acquisition Software for Neuroscience Experiments. Front. Neural Circuits 4, (2010).

72. Huang, K.-H. et al. A virtual reality system to analyze neural activity and behavior in adult zebrafish. Nat. Methods 17, 343–351 (2020).

73. Pecka, M., Han, Y., Sader, E. & Mrsic-Flogel, T. D. Experience-Dependent Specialization of Receptive Field Surround for Selective Coding of Natural Scenes. Neuron 84, 457–469 (2014).

74. Pernía-Andrade, A. J. et al. A Deconvolution-Based Method with High Sensitivity and Temporal Resolution for Detection of Spontaneous Synaptic Currents In Vitro and In Vivo. Biophys. J. 103, 1429–1439 (2012).

75. Guzman, S. J., Schlögl, A. & Schmidt-Hieber, C. Stimfit: quantifying electrophysiological data with Python. *Front*. Neuroinformatics 8, (2014).

76. GENIE project, J. F. C., Hhmi & Svoboda, K. Simultaneous imaging and loose-seal cell-attached electrical recordings from neurons expressing a variety of genetically encoded calcium indicators. CRCNS.org. (2015).

77. Boaz, M., Dana, H., Kim, D. S., Svoboda, K. & GENIE project, J. F. C., HHMI. jRGECO1a and jRCaMP1a characterization in the intact mouse visual cortex, using AAV-based gene transfer, 2-photon imaging and loose-seal cell attached recordings, as described in Dana et al 2016. CRCNS.org. (2016).

78. Reynolds, S., Abrahamsson, T., Sjöström, P. J., Schultz, S. R. & Dragotti, P. L. CosMIC: A Consistent Metric for Spike Inference from Calcium Imaging. Neural Comput. 30, 2726–2756 (2018).

79. Ioffe, S. & Szegedy, C. Batch Normalization: Accelerating Deep Network Training by Reducing Internal Covariate Shift. Preprint at https://arxiv.org/abs/1502.03167 (2015).

80. Hochreiter, S. & Schmidhuber, J. Long Short-Term Memory. Neural Comput. 9, 1735–1780 (1997).

81. Gers, F. A., Schmidhuber, J. & Cummins, F. Learning to forget: continual prediction with LSTM. 850–855 (1999) doi:10.1049/cp:19991218.

82. Schuster, M. & Paliwal, K. Bidirectional recurrent neural networks. Signal Process. IEEE Trans. On 45, 2673–2681 (1997).

83. Graves, A., Fernandez, S. & Schmidhuber, J. Bidirectional LSTM Networks for Improved Phoneme Classification and Recognition. 6.

84. Eden, U. T. & Kramer, M. A. Drawing inferences from Fano factor calculations. J. Neurosci. Methods 190, 149–152 (2010).

85. Fort, S., Hu, H. & Lakshminarayanan, B. Deep Ensembles: A Loss Landscape Perspective. Preprint at https://arxiv.org/abs/1912.02757 (2019).

86. Beaulieu-Laroche, L., Toloza, E. H. S., Brown, N. J. & Harnett, M. T. Widespread and Highly Correlated Somato-dendritic Activity in Cortical Layer 5 Neurons. Neuron 103, 235–241.e4 (2019).

